# Alpha diversity analysis of hepatic transcriptome reveals novel pathways in alcohol-related hepatitis

**DOI:** 10.1101/2025.10.08.681263

**Authors:** Sudrishti Chaudhary, Jia-Jun Liu, Silvia Liu, Marissa Di, Juliane I Beier, Ramon Bataller, Josepmaria Argemi, Panayiotis V Benos, Gavin E Arteel

**Author notes:** Send all correspondence to: Gavin E. Arteel, PhD, FAASLD, Thomas E. Starzl Biomedical Science Tower West 1143, 200 Lothrop Street, Pittsburgh, PA 15213. Conflict of interest: The authors have declared that no conflict of interest exists.

## Abstract

Next generation sequencing can identify novel gene expression patterns in disease. Beyond differentially expressed genes analysis, we investigated the ability of within-population diversity (α-diversity) of the transcriptome to reveal new biological information in alcohol-related liver disease (ALD), comparing Differential Shannon diversity (DSD) to transcriptome heterogeneity changes. RNA sequencing data from normal livers and patients with early silent ALD and severe AH were analyzed. α-diversity indices and Percent Shannon Diversity of a gene, which refers to this gene’s contribution to total Shannon entropy were calculated. Ingenuity pathway analysis identified canonical pathways determined by differentially expressed genes (DEG) and DSD approaches. ALD significantly decreased hepatic transcriptome α-diversity correlating with increased relative contribution of select genes. These changes were driven by lower abundance gene expression loss. DEG and DSD analyses showed overlapping genes and canonical pathways, but DSD also identified novel genes and pathways not highlighted by DEG. Importantly, DSD more effectively identified differences between preclinical ALD and AH severity stages. ALD decreases hepatic transcriptome heterogeneity, favoring pathways associated with organ damage or damage response. Preclinical and clinical ALD led to differential heterogeneity patterns that may provide new disease insights. DSD analysis identified enriched pathways missed by standard DEG analyses, potentially yielding novel insight into disease mechanisms and biomarkers.

**Graphical abstract:** 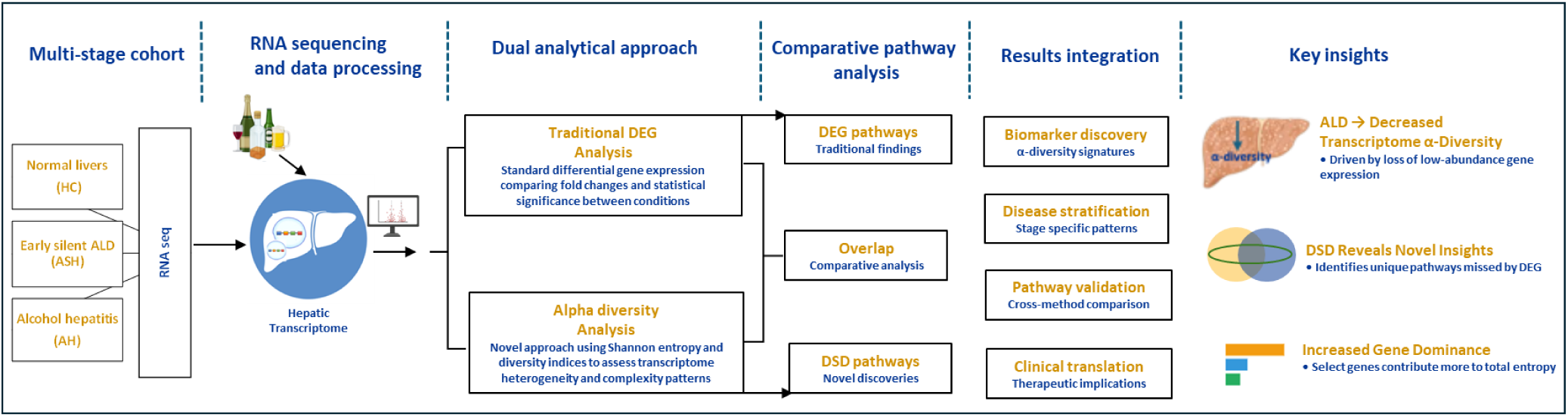

## Introduction

Alcohol-related hepatitis (AH) is a subacute form of alcohol-related liver disease (ALD), characterized by jaundice, ascites and other sequelae associated with severe hepatic decompensation. AH has a high mortality rate of 30-50% at 3 months and 40% at 6 months.(1) Although the clinical progression of AH has been well-described for decades, the underlying mechanisms that drive AH are incompletely understood. The only FDA-approved therapy for AH (corticosteroids), has been used since the 1950s, despite having only limited efficacy in AH patients.(2) Hence, the need arises for developing better therapeutic options, for which it is crucial to understand the underlying mechanisms of AH.

To address these gaps in understanding, the National Institute on Alcohol Abuse and Alcoholism (NIAAA) launched AlcHepNet in 2012; this multi-center translational and clinical research program was designed to accelerate the understanding of AH. This program also led to the storage of large cohorts containing clinical data and biobanked samples, which have been available to researchers in the field for primary and secondary analyses. These studies and the establishment of clinical biobanks have paved the way for future research and development of new information and insights for AH. For example, multi-platform next generation sequencing (NGS) of transcription from liver biopsies highlighted expression changes in AH patients that suggest a pseudo-fetal hepatic function.(3)

NGS is a powerful tool for detecting gene expression patterns, also uncovering unrecognized or cryptic gene expression in disease states.(4) Differentially expressed genes (DEG) analysis has been established as the common gene analysis method, focusing on genes that are significantly upregulated (expressed at higher levels) or downregulated (expressed at lower levels) under different conditions.(5) These approaches have been useful in detecting targets or biomarkers and helping generate hypotheses and mechanisms behind diseases. However, DEG analysis assumes transcriptional independence, as it focuses on individual genes. For this reason, DEG analysis is often coupled with pattern recognition algorithms (e.g., Ingenuity Pathway Analysis; IPA) to identify changes in enrichment in specific biological pathways. However, pattern recognition approaches rely on curated knowledge, which is incomplete, and do not agnostically identify novel patterns.(6)

In addition to analysis at the individual gene level, transcriptomic data are often analyzed using algorithms developed from ecosystem diversity analyses. For example, β-diversity indices and multivariate analyses are often employed to describe how the overall transcriptome differs between two different states (e.g., diseased vs. healthy). β-Diversity compares the gene expression pattern between conditions and quantifies the similarity or dissimilarity of the different groups. For example, Principal Component Analysis (PCA) clusters samples that have similar expression profiles (i.e., low β-diversity). These approaches build on the understanding that environmental stressors often impact the totality of gene expression and provide a more holistic view of comparative transcriptome dynamics. However, these approaches are used predominantly for visualizing the overall changes in gene expression and not for identifying novel mechanisms and pathways.(7)

In contrast to between-sample diversity (i.e., β-diversity), within-sample diversity (i.e., α-diversity) is rarely used to analyze transcriptomic data sets. Moving away from the traditional DEG approach, this study employs indices of α-diversity and introduces a novel analysis of gene expression based on α-diversity (i.e., Differential Shannon Diversity; DSD). Unlike DEG analysis, which focuses on individual gene expression changes and can be sensitive to outliers, or indices of β-diversity, which measure variation between samples but may not pinpoint specific genes, α-diversity indices like Shannon diversity can provide a more holistic view of transcriptome complexity while still allowing for gene-level analysis. The DSD approach allows for the quantification of each gene’s contribution to overall transcriptome diversity, potentially revealing subtle but important changes that might be missed by traditional methods. The purpose of this study was therefore to determine the effect of AH on indices α-diversity of the hepatic transcriptome, as well as investigate the pattern of gene expression determined by DSD.

## Results

### Alcohol-related liver disease decreases the α-diversity of the hepatic transcriptome

Figure 1A outlines a transcriptome study from the InTEAM Consortium (NLM study code phs001807.v1.p1) comparing gene expression profiles across disease groups: healthy controls (10 subjects), early silent alcoholic liver disease (11 subjects), and alcohol hepatitis patients (18 subjects). We further performed the secondary analysis which specifically focused on RNA sequencing data to understand how gene expression patterns change during alcoholic liver disease progression. α-diversity analysis was performed to assess transcriptome diversity through Shannon index, evenness, and dominance metrics, along with abundance analysis that rank genes from lowest to highest expression and calculates rank differences between groups. We also conducted pathway analysis using IPA (Ingenuity Pathway Analysis) to identify differentially regulated biological pathways across the progression from healthy liver to alcohol hepatitis. To evaluate potential differences in α-diversity profiles of individuals with the disease spectra (ASH, AH) and healthy controls (HC), α-diversity was quantified by richness, evenness, dominance, and related indices (Figure 1B-G). The Menhinick‘s index attempts to estimate species richness independent of sample size. It is calculated here as the number of genes with non-zero expression in the sample divided by the square root of the sum of read counts of all genes in the sample. ASH significantly decreased the Menhinick index, compared to HC and AH groups (Figure 1B). The Brillouin (Figure 1C) and Shannon (Figure 1D) diversity indices showed a stepwise reduction from HC to ASH to AH. Indices of Equitability (Figure 1E) and Evenness (Figure 1F) followed a similar trend, indicating that the distribution of level of gene expression became less uniform with ALD disease severity. The Dominance_D index (Figure 1G) shows an inverse pattern with AH having the highest value. Taken together, these results collectively suggest a progressive decrease in hepatic gene expression diversity and evenness, coupled with an increase in dominance, as liver disease severity progresses from healthy to ASH to AH.

**Figure 1.**
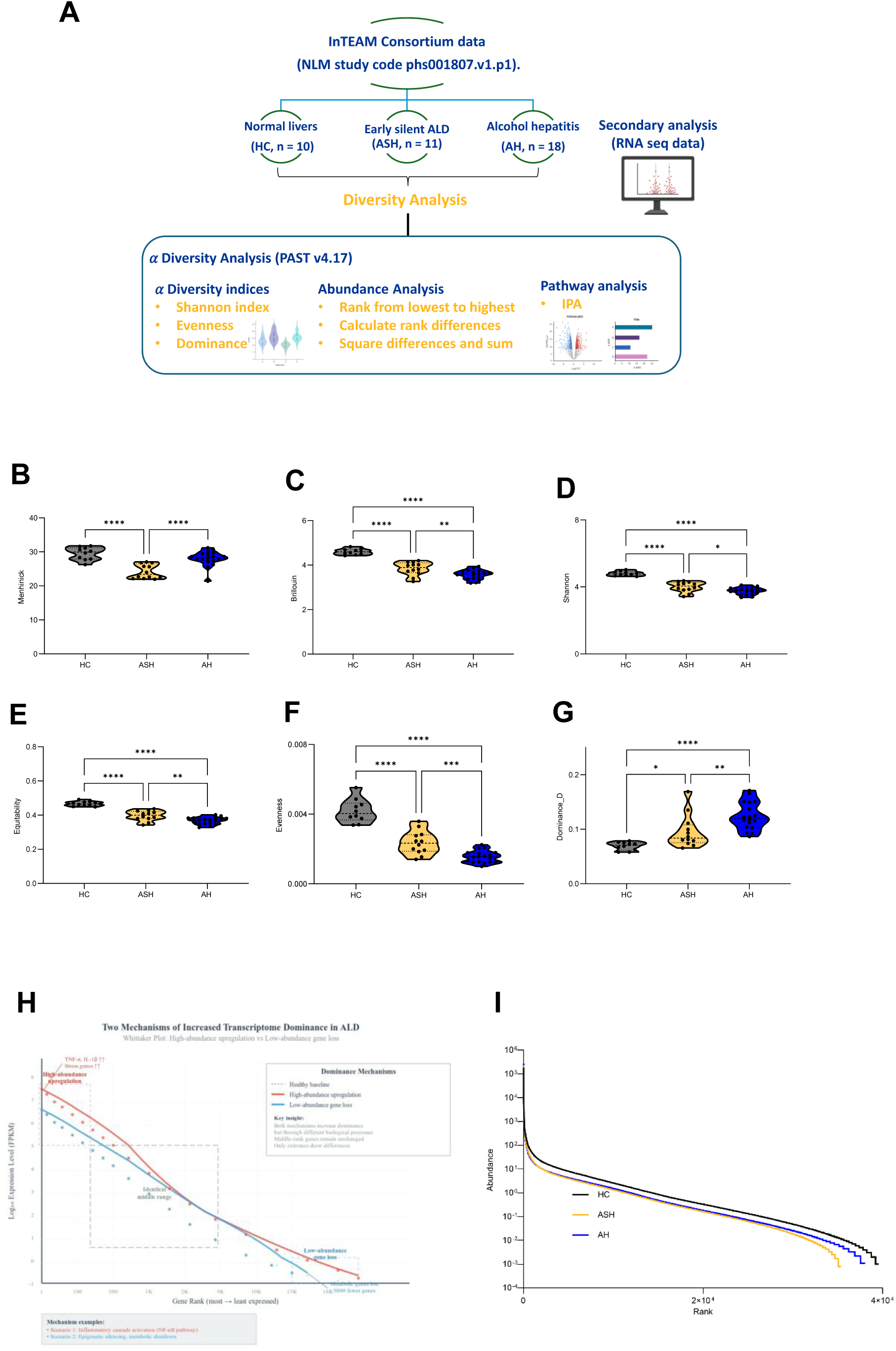
Study design and analytical workflow for transcriptome analysis of alcohol liver disease progression. **(A)**Secondary analysis on RNA sequencing data to characterize transcriptional changes during disease progression, examining gene expressions across healthy controls (HC, n=10), early silent alcoholic liver disease (ASH, n=11), and alcohol hepatitis patients (AH, n=18). Analysis included α-diversity assessment using PAST v4.17 (Shannon index, evenness, dominance), abundance analysis with gene ranking and rank differences, and pathway analysis using IPA. **(B-G). Indices of alpha diversity.** Violin plots showing the distribution of alpha-diversity as measured by (A) Menhinick index (B) Dominance (C) Shannon index (D) Brillouin (E) Evenness and (F) Equitability index for the four groups: HC (Healthy control), ASH (Alcohol-related steatohepatitis) AH (Alcohol-related hepatitis responders). One way ANOVA analysis revealed significant grouping between groups(p<0.05). **(H, I). Diversity and changes by abundance.** (H) Mechanisms of Transcriptome Dominance in Alcohol Liver Disease. Whittaker plot shows two opposing routes to increased dominance: inflammatory upregulation of highly expressed genes (red) and loss of low-abundance metabolic genes (blue) relative to healthy liver (gray dashed line). Mid-ranked genes remain stable. Colored boxes indicate distinct biological processes with unique therapeutic implications. (I) Whittaker plots showing changes by abundance and prevalence rank. Relative abundance-rank curve of alpha diversity in different groups based on average of each gene per group.

### Diversity and changes by abundance

The above-described changes to indices of diversity indices (Figure 1B-1G) can be generally described by either an increase in the relative expression of highly abundant genes or via a loss of expression in low abundance genes. The schematic represented in Figure 1H demonstrates two distinct mechanisms by which Alcohol Liver Disease achieves increased transcriptome dominance. The red curve shows high-abundance gene upregulation, where already highly expressed genes are further amplified through inflammatory cascades, effectively concentrating transcriptional resources among dominant genes. The blue curve illustrates low-abundance gene loss, where weakly expressed genes are systematically silenced through epigenetic suppression or metabolic shutdown, reducing the total transcriptome diversity. Importantly, middle-ranked genes remain unchanged in both scenarios, indicating that dominance increases through changes at expression extremes rather than global shifts. This reveals that disease-associated transcriptome dominance can result from either amplifying dominant pathways or eliminating minor contributors, with significant implications for targeted therapeutic strategies. Figure 1I provides a comprehensive analysis of gene expression as a factor of expression abundance to visualize these potential mechanisms. It shows the pattern of abundance as a function of prevalence rank (Whittaker plot). This plot shows a similar overall pattern of gene expression between the groups, with subtle differences apparent only at the lower abundance genes as shown in schematic represented in Figure 1H.

### Pathway analysis of significantly changed genes determined by DEG and DSD

The above-described analyses (Figure 1) indicate an interesting effect of ALD disease severity on indices of transcription diversity, indicating a stepwise deregulation of lower abundance genes in ASH and AH. However, these analyses describe global changes in the transcriptome and do not highlight specific changes in genes. The effect of ALD disease severity on changes in gene expression was therefore determined using Differential Shannon Diversity (DSD; see Methods for details) and compared with traditional DEG analysis. The volcano plots (-Log_10_PV as a function of Log_2_FC) and Venn diagrams of the DEG and the DSD data sets are presented in Figure 2(A-I). DEG and DSD analyses yielded quantitatively similar results, as depicted by volcano plots for ASH vs. HC (Figure 2A and B), AH vs. HC (Figure 2D and E) and AH vs. ASH (Figure 2G and H). Figure 2C, F and I depict Venn diagrams of genes identified by DEG and/or DSD that were significantly upregulated or downregulated and were common or unique to both the approaches. Venn diagrams for each of the diseases ASH/HC (Figure 2C), AH/HC (Figure 2F) and AH/ASH (Figure 2I) depict significant overlap between genes chosen by both DEG and DSD techniques. Moreover, there were genes uniquely identified by both analytical approaches. For example, both DEG and DSD analysis identified more unique genes in the AH vs. ASH group compared to other groups with respect to healthy controls; thus, the uniquely identified genes were more balanced between the approaches in the AH vs. ASH group. The above results revealed an overall pattern of gene expression between the groups, with subtle differences apparent only at the lower abundance genes. The prevalence percentage was calculated for the unique as well as common set of genes derived from the Venn analysis in all the disease states (Figure 2J-L), which also supported the above results that show that the high prevalence genes were generally less impacted by both disease states, and the middle and lower show higher variability in response to ASH and AH. The percent of coefficient variation (CV), calculated by the standard deviation divided by the mean, showed no difference between DEG and DSD analysis in gene expression across the groups. (Supplemental Figure S1).

**Figure 2.**
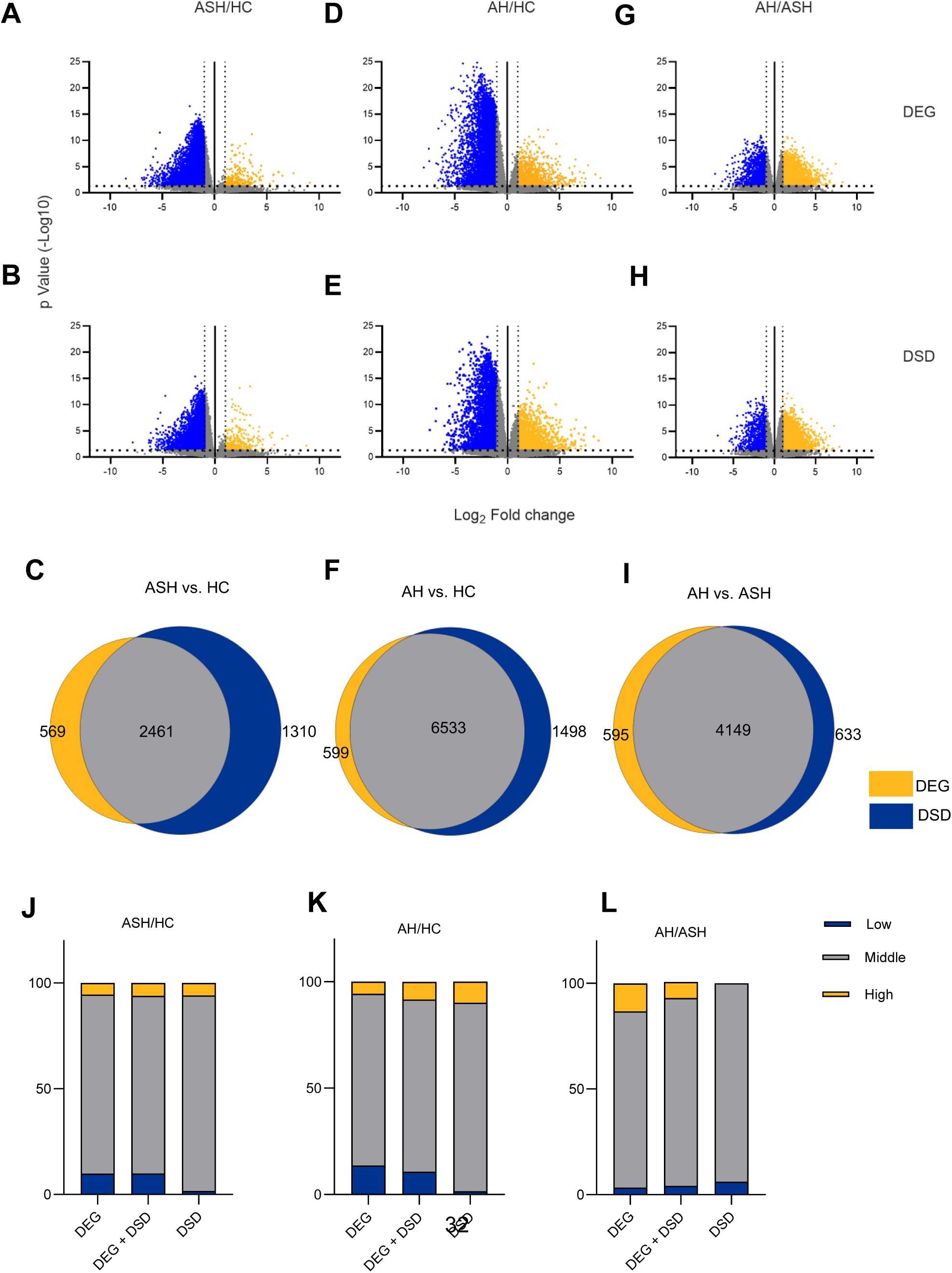
DEG vs. DSD: Shared and unique changes. Volcano plot representation of differentially expressed genes (DEG) and the Differential Shannon Diversity (DSD) data sets in ASH vs. HC (A, B), AH vs. HC (D, E) and AH vs. ASH (G, H). The yellow and blue points mark the increased or decreased gene expression respectively. The x-axis shows log_2_fold-changes and the y-axis the -log of p value (p>=1.3) of the gene expression. Venn diagrams depicting unique and common subsets of genes shared by both differentially expressed genes (DEGs) and Differential Shannon diversity (DSD) approaches in (C) ASH vs HC, (F) AH vs HC and (I) ASH vs AH groups. Bar diagrams (J, K & L) depicting percentage of low, medium and high abundance genes across the groups, common and unique to DEG and DSD.

Figure 3 (A-I) shows a modified volcano plot depicting the results of Ingenuity Pathway Analysis (IPA) of significantly changed genes determined by DEG and DSD in the various comparisons where the various GO terms enriched (-Log_10_PV) were plotted as a function of the Z-score. These Bubble plots showed significantly enriched pathways both common and exclusive for the DEGs and DSDs in disease groups ASH, AH vs HC and AH vs ASH. IPA identified both overlapping and unique canonical pathways by both analyses in each of the groups (Supplemental table S2-S10). The DSD analysis identified novel genes and pathways not highlighted by DEG approach specifically, in the AH vs ASH group. Some of these pathways have been established to potentially contribute to ALD e.g., Fatty acid oxidation. GO: 0019395, p=1.34×10^-5^ (Supplemental table S8-S10).

**Figure 3.**
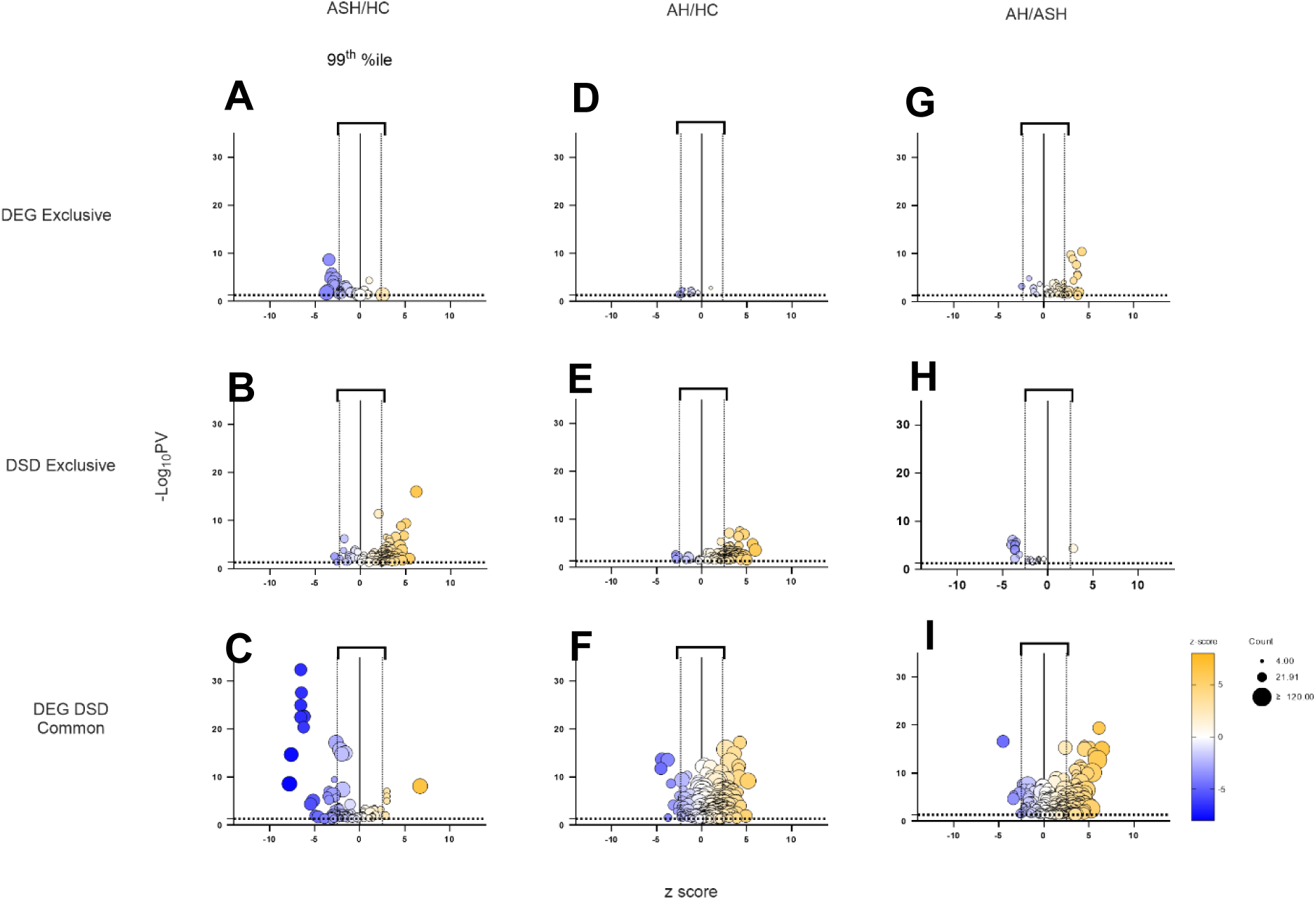
Pathway enrichment analysis of DEG and DSD. Bubble plot showing significantly enriched pathways both common and exclusive for the differentially expressed genes (DEGs) and Percent Shannon Diversity (DSD) in disease groups ASH vs HC**(A-C)** and AH vs HC**(D-F)** and AH vs. ASH**(G-I).** Size of the bubbles is proportional to the gene count. The y-axis represents the negative logarithm of the Log_10_ p value for the genes, and the x-axis displays the z-score. The threshold for displaying the bubble labels was set to a Z score of 2.3. Bubbles for genes belonging to high z score are depicted in yellow and low z score in blue.

IPA analysis of significantly changed genes determined by DSD analysis identified differences between preclinical ALD (ASH), and more severe disease state (AH) more effectively than DEG analysis. The pathways from their scores clustered into categories were identified. Canonical Pathway scores plotted as pathway category based on gene expression level for the DSD exclusive approach in the AH vs ASH group is presented in Figure 4. (The other groups data are presented in the supplementary Figure S2, S3). The top genes identified by DEG did not include a significant number of disease specific pathways in contrast to the genes identified by DSD. The enrichment analysis using DSD approach revealed pathways most significantly enriched in categories related to Degradation of extra cellular matrix, Inhibition of matrix metalloproteases, Cellular stress and injury, apoptosis and Cellular immune response (Figure 4).

**Figure 4.**
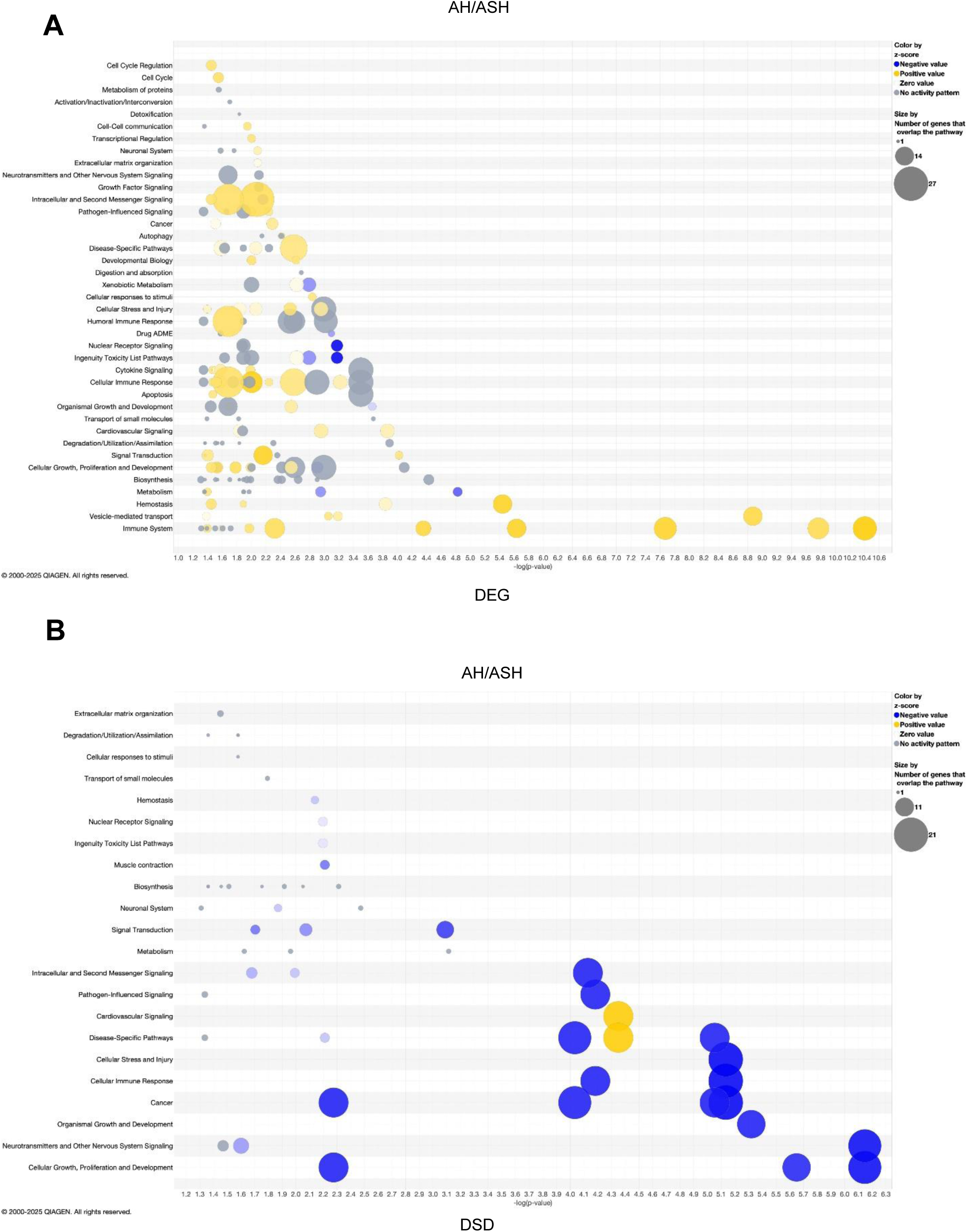
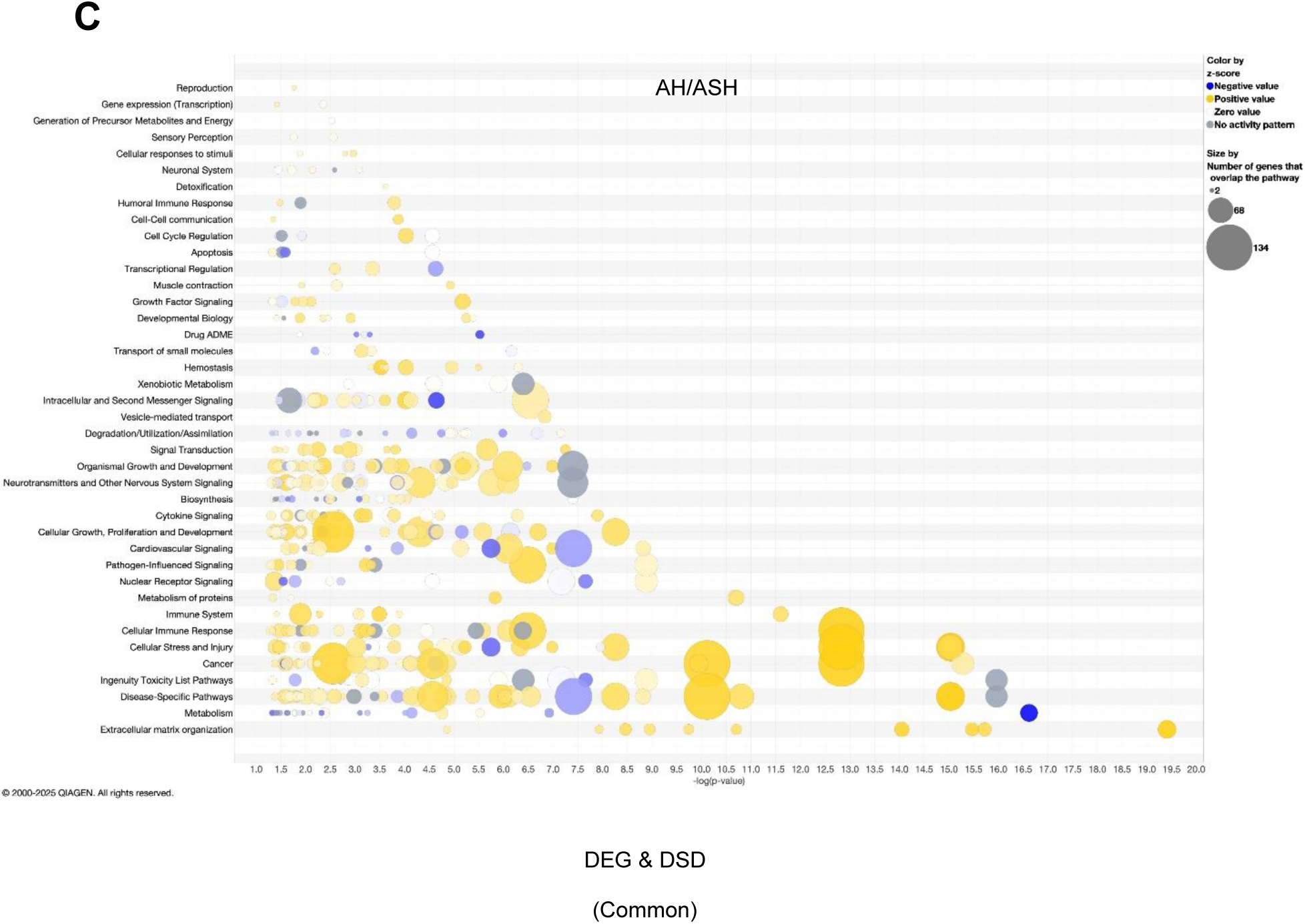
Canonical Pathway bubble chart. Canonical Pathway scores plotted as pathway category based on gene expression level for the DEG exclusive **(A)** and DSD exclusive **(B)** and common pathways **(C)** in the AH vs ASH group. The log of the p-value for each pathway is plotted on the x-axis versus the pathway categories plotted along the y-axis. By default, the coloring relates to the pathway’s z-score, and the bubble size relates to the number of dataset genes that overlap each pathway, as shown in the legend.

### Alpha Diversity Analysis of Hepatic Transcriptome in Alcohol-Related Liver Disease

This figure illustrates the application of ecological diversity concepts to analyze hepatic gene expression changes across the spectrum of alcohol-related liver disease (ALD). An ecological framework demonstrates parallel patterns of biodiversity loss between natural ecosystems and liver transcriptomes (Figure5A). Environmental stressors drive ecosystem degradation from diverse communities (left) to homogeneous landscapes with reduced species diversity (right), mirroring transcriptome changes in disease progression. Disease progression from healthy controls (HC, green) to early silent ALD (ASH, yellow) to alcohol-related hepatitis (AH, red) involves progressive changes in liver architecture, with increasing lipid accumulation and disruption of normal hepatic structure. Shannon diversity index, a measure of transcriptome heterogeneity, shows stepwise reduction with disease severity, from high in HC to medium in ASH to low in AH. Gene expression evenness/dominance patterns illustrate how transcriptome composition shifts from high evenness (multiple similarly expressed genes in HC) to low evenness/high dominance (few genes dominating expression in AH). Disease-induced transcriptome changes disproportionately affect genes based on their abundance levels, with low abundance genes showing the most dramatic expression changes while high abundance genes remain relatively stable. This ecological perspective demonstrates how chronic alcohol exposure reduces transcriptome diversity in the liver, similar to how environmental stress reduces biodiversity in natural ecosystems.

### Cellular composition changes across ALD disease progression

Cellular deconvolution analysis using 528 validated marker genes across 9 major liver cell types was performed to estimate cellular composition (Figure 5B). ALD caused progressive changes in cellular composition across the disease spectrum. Hepatocyte proportions decreased stepwise: HC (64.3% ± 3.2) → ASH (58.7% ± 4.1) → AH (52.1% ± 5.8) (p < 0.001), representing a 19% total reduction from healthy to severe disease consistent with established ALD progression.(8, 9) Kupffer cell proportions increased across disease stages: HC (12.1% ± 1.8) → ASH (16.8% ± 2.3) → AH (22.3% ± 3.1) (p < 0.001), representing an 84% increase in severe disease that aligns with scRNA-seq studies demonstrating macrophage activation in ALD.(10, 11) Stellate cell activation occurred progressively: HC (8.9% ± 1.3) → ASH (12.4% ± 1.8) → AH (16.2% ± 2.4) (p < 0.001), likely reflecting the fibrotic response characteristic of advanced ALD, while endothelial cell proportions showed modest decreases with disease progression: HC (7.2% ± 1.1) → ASH (6.9% ± 1.0) → AH (6.1% ± 1.2). Adaptive immune cells showed progressive depletion, with T cell proportions declining from 4.1% ± 0.8 in controls to 2.4% ± 0.6 in severe disease (p < 0.01), B cells decreasing from 2.3% ± 0.5 to 0.8% ± 0.2 (p < 0.001), and NK cells showing an 86% reduction from 0.7% ± 0.2 to 0.1% ± 0.1 (p < 0.001). Neutrophils and monocytes were virtually absent in severe disease (< 0.1%) (Supplemental table S1). These cellular composition changes occurred in parallel with the transcriptome diversity reduction described in the α-diversity analysis. The observed patterns of hepatocyte loss, Kupffer cell expansion, and adaptive immune cell depletion are consistent with findings from recent human scRNA-seq studies in alcohol-related liver disease,(9–11)validating the utility of bulk RNA deconvolution approaches for detecting cellular remodeling in liver disease progression.

**Figure 5.**
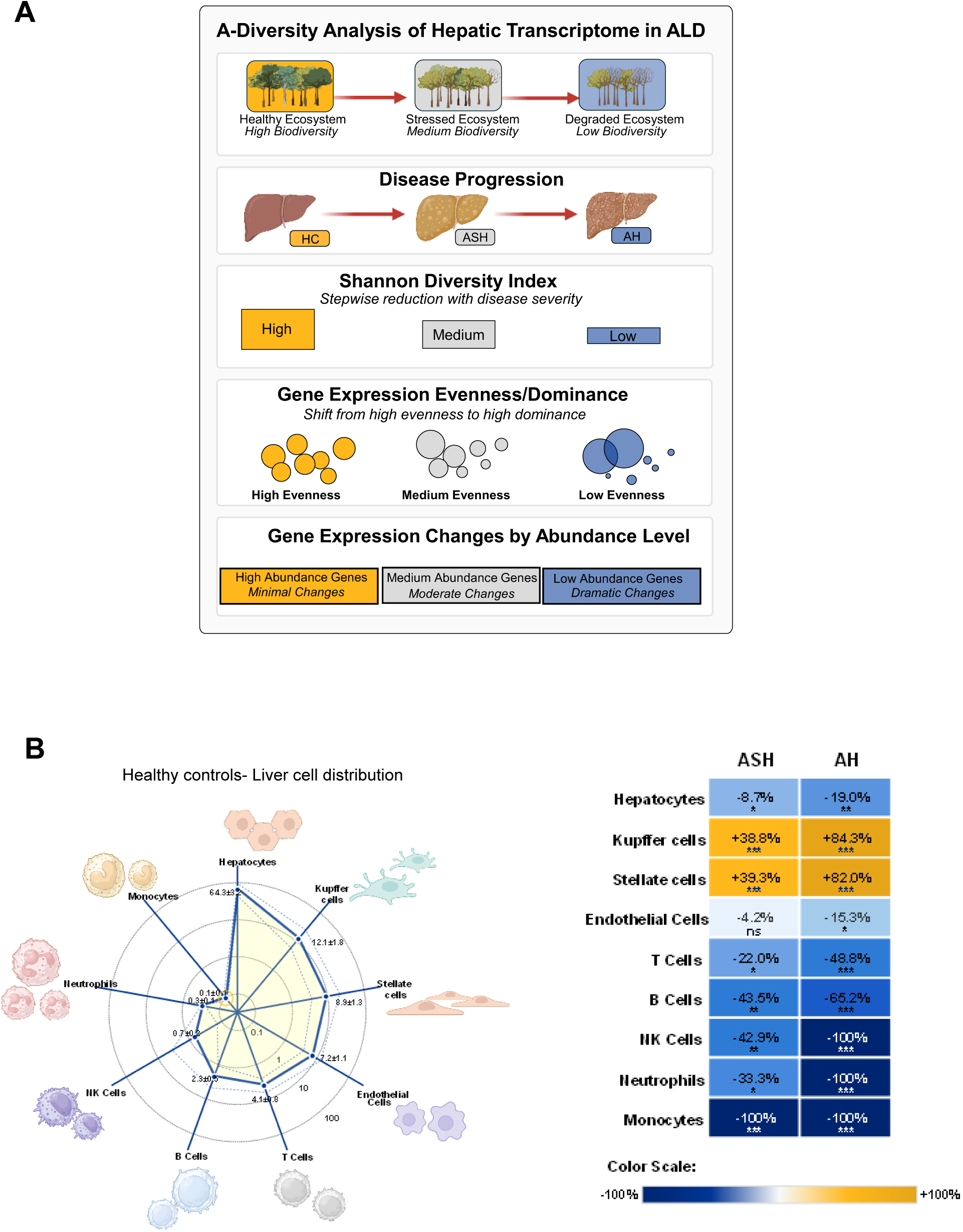
Conceptual framework for α-diversity analysis of hepatic transcriptome in alcoholic liver disease progression. **(A)** Schematic illustration of transcriptome diversity changes during liver disease progression using ecological diversity principles. Ecosystem analogy shows transition from healthy (high biodiversity) to stressed (medium biodiversity) to degraded (low biodiversity) states. Corresponding liver disease progression from healthy controls (HC) through early silent alcoholic liver disease (ASH) to alcohol hepatitis (AH). Shannon Diversity Index demonstrates stepwise reduction in transcriptome diversity with increasing disease severity. Gene expression evenness and dominance patterns shift from high evenness (many genes with similar expression levels) to high dominance (few highly expressed genes). Abundance-dependent gene expression changes showing minimal alterations in high abundance genes, moderate changes in medium abundance genes, and dramatic changes in low abundance genes during disease progression. **(B)** Liver Cell Type Distribution. Panel B is a radar plot displaying liver cell type distribution in healthy controls (n=10) on log₁₀ scale. Data points show mean values positioned on radial axes with error bars representing ±1 SEM. Dashed lines indicate upper and lower confidence boundaries (mean ± SEM). Vector length is proportional to log₁₀(cell count), effectively visualizing the 643-fold dynamic range from 0.1 (Monocytes) to 64.3 (Hepatocytes) cells per field. Gold circle around Monocytes indicates high measurement uncertainty (±100% CV). Alongside is a heatmap showing percent change from healthy controls across disease conditions with integrated statistical significance indicators. ASH (n=12) and AH (n=18) columns display changes for each cell type. Complete cell loss represented as -100% change in dark blue. White indicates minimal change (∼0%). Statistical significance displayed below each percentage value: *** p<0.001, ** p<0.01, * p<0.05, ns = not significant (Welch’s t-test vs. healthy controls).

## Discussion

Our study explored whether ecological alpha diversity metrics could provide novel insights in analyzing differential gene expression in transcriptomic data. Our work suggests that chronic alcohol exposure decreases overall transcriptome diversity in the liver, inducing various stress response pathways. While diversity is a fundamental concept in community ecology,(12–14) its application to transcriptomics remains relatively unexplored. Ecological diversity measures have long been used to quantify species distribution patterns, with alpha diversity defined as local diversity within a single area or ecosystem and beta diversity describing changes in species composition across areas.(12) We hypothesized that applying these established ecological frameworks to transcriptomic data—conceptualizing the transcriptome as an ecosystem and individual genes as species—could reveal patterns and mechanisms missed by traditional analytical approaches. This novel perspective allows us to investigate how environmental stressors, such as alcohol consumption, might affect the overall “ecosystem health” of the transcriptome, potentially disrupting expression patterns in ways analogous to how environmental stresses impact natural ecosystems. While this ecological lens has been applied in microbial community analysis(15, 16) its application to human transcriptomics represents an innovative cross-disciplinary approach that may provide deeper insights into disease pathophysiology and progression, particularly in complex conditions like Alcohol-related Liver Disease (ALD).

Beta diversity measures, particularly through principal component analysis (PCA), have been extensively used in transcriptomics to examine differences between populations or sample groups.(16–18) These approaches help visualize variation across different conditions and identify major sources of transcriptome heterogeneity. Beta diversity metrics have proven particularly valuable in RNA-seq studies for detecting global patterns of gene expression changes across experimental conditions, disease states, and tissue types.(5, 10, 11, 19) Researchers have employed distance matrices and ordination techniques like PCA, non-metric multidimensional scaling (NMDS), and t-SNE to relate gene expression patterns to phenotypic variables, treatment effects, and developmental stages.(8, 10, 20) In contrast, alpha diversity metrics in transcriptomics have typically been limited to describing and quantitating within-group variability(21), serving primarily as quality control measures or indicators of overall transcriptome complexity. Few studies have explored their potential for between-group comparisons in disease states and progression monitoring, despite their established utility in ecology for measuring ecosystem health and stability. This gap represents a significant opportunity to develop new analytical frameworks that could provide complementary insights to traditional differential expression analyses.

Analyzing various alpha diversity indices across Healthy Control (HC), Alcohol-related Steatohepatitis (ASH), and Alcohol-related Hepatitis (AH) samples revealed a clear pattern: highest diversity in HC, followed by ASH, with lowest in AH, confirming that ALD decreases hepatic transcriptome diversity. We systematically evaluated multiple ecological diversity metrics including Menhinick richness index, Shannon entropy, Brillouin index, and dominance measures to quantify transcriptome heterogeneity across disease states. Our analysis demonstrated a stepwise reduction in diversity metrics that correlated with disease severity, suggesting a progressive disruption of normal gene expression patterns with advancing ALD. Whittaker plots and Log_2_FC scatter plots demonstrated that lower abundance genes show more pronounced expression changes under disease conditions, consistent with our gene prevalence analysis. This finding is particularly significant as it indicates that disease-related transcriptional changes disproportionately affect genes expressed at lower levels, potentially explaining why traditional differential expression approaches—which often favor highly expressed genes—might miss important pathological mechanisms. The pattern we observed mirrors ecological systems under stress, where rare species often exhibit greater sensitivity to environmental perturbations than dominant ones.(22–24)

The healthy liver presents a rich, diverse transcriptome that chronic alcohol exposure disrupts, similar to how environmental stress decreases genetic diversity in natural populations. Chronic alcohol consumption creates a cellular environment characterized by oxidative stress, inflammation, and metabolic dysregulation that places ‘selective pressure’ on gene expression patterns. Several recent studies have examined alcohol-induced transcriptomic changes using RNA sequencing,(5) multi-omics approaches,(11) and single-cell transcriptome research,(9, 10) but few have applied diversity metrics as analytical tools. RNA sequencing studies in murine models exposed to both chronic and binge ethanol feeding have documented widespread changes in gene networks governing lipid metabolism, inflammation, and cellular stress responses, paralleling many of the pathways we identified through our diversity-based approach.

Comprehensive omics studies(20) and research on transcriptional dynamics(25) have further expanded our understanding, with recent single-cell RNA-seq studies mapping liver cell heterogeneity in alcohol models and revealing cell-type specific vulnerability to alcohol toxicity. This growing body of literature confirms alcohol’s profound effects on hepatic gene expression but has largely overlooked the ecological perspective of transcriptome diversity that our study introduces.

A novel method to analyze gene expression data called Differential Shannon Diversity (DSD) was developed in this study. DSD relies on Shannon diversity index, capable of measuring both richness and evenness of gene abundance for analyzing expression changes. When compared to Differential Expression Gene (DEG) analysis, both methods generated overlaps and differences in gene sets. To look macroscopically, the patterns between DEG and DSD were very similar implying that most of the genes were following the same pattern, but important unique changes were revealed through deeper analysis. Looking at the Venn diagrams (Figure 2C, F, I), while most of the genes followed the same pattern, there were genes uniquely identified by DEG analysis, and notably, genes that were uniquely identified by DSD. Interestingly, our analysis revealed that while gene richness (number of different transcripts) initially decreased in ASH compared to normal, it increased again in AH, yet Shannon diversity continued to decline. This suggests that AH is characterized by the emergence of more transcript types (possibly including low-expressed fetal transcripts) but with decreased evenness due to dramatic expression changes in a subset of transcripts responding to liver failure. The DSD technique helped identify genes not found by differential expression analysis, with each metric independently finding disease-relevant genes, such as MITF/p300/CBP complex (p=3.48×10^-4^) a CREB binding protein, which acts as a transcription regulator) and Spring1 (p=6.89×10^-5^) involved in SREBP signaling and cholesterol metabolism. Markedly, as compared to DEG, DSD proved to be a more sensitive method in detecting changes in highly expressed genes, even with small fold changes. Unlike DEG analysis, DSD was capable of detecting shifts in gene proportions, even when the absolute gene expression levels did not change.

Our study thus suggests that prolonged alcohol exposure reduces the variety of RNA transcripts in liver tissue while activating multiple stress response mechanisms.

Integrating diversity methodologies through DSD reveals enriched pathways not highlighted by standard approaches. Looking forward, changes in transcriptome diversity could potentially serve as biomarkers for ALD progression or treatment response, though future research should determine whether these findings are specific to AH or generalized to all exogenous stresses.

## Methods

*Sex as a biological variable.* This study performed secondary analysis on publicly available RNA sequencing data with both male and female patients. Sex was not considered as a biological variable in the study.

### Publicly available Liver RNA Sequencing Analysis

RNA sequencing data were obtained from normal livers (n = 10) and from biopsies of patients with early silent ALD (ASH, n = 11), non-severe AH (n = 9), and severe AH (n = 9) from the InTEAM Consortium – Alcohol-related Hepatitis Liver RNA Sequencing study, sponsored by the National Institute of Alcohol Abuse and Alcoholism (NIAAA, USA). The study details and sequencing data can be found in the Database of Genotypes and Phenotypes (dbGAP, phs001807.v1. p1) of the National Institutes of Health (NIH, USA). The basic clinical and laboratory data of the patients included in this study, the methods used to extract RNA and perform deep RNA sequencing, and the bioinformatic pipelines used to determine transcript counts have been described previously.(8) Indices of α-diversity (e.g., Shannon index, evenness and dominance) were calculated using PAST (v. 4.17) software. The abundance rank correlation coefficient was calculated, each data set was ranked from lowest to highest, then rank differences and square of each of those differences was calculated, followed by the calculation of the sum of all the squared differences. This was calculated as a Spearman rank correlation function in excel.

### Differentially expressed gene (DEG) and Differential Shannon Diversity (DSD) analyses

The impact of ALD and AH on gene expression was measured by the Log2 fold-change between groups. We tested two differential metrics; one based on the differentially expressed gene (DEG) and the other on the normalized Differential Shannon Diversity (DSD). The probability of observing gene *i* in this sample (*p*_*i*_) was calculated as detailed in Equation 1:

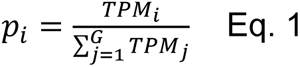

Where TPM represents the transcript per million, *G* is defined as the total number of expressed genes for a given sample.

Next, the Log_2_-based Shannon entropy (*H*) was used to measure the RNA diversity for a given library, which is calculated as detailed in Equation 2:

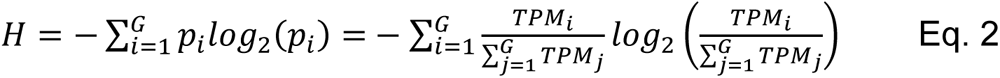

*Here H* ranges from 0 to *log*_2_(*G*). If *H* = 0 , only one gene is expressed for that library. And if *H* = *log*_2_(*G*), all the genes are evenly expressed.

In order to quantify the expression weight of gene *i* among all the genes of a given sample, the Percentage Shannon Entropy (PSE) was defined to illustrate the percentage of gene *i* in terms of Shannon entropy. This measure represents the contribution of each individual gene to the overall Shannon entropy of the transcriptome, normalized by the total entropy.

Mathematically:

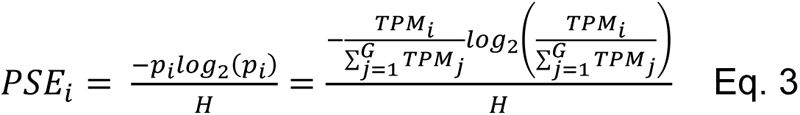

PSE values range from 0 to 1, where 0 indicates that gene *i* is not expressed and PSE will be close to 1 if only gene i is expressed for a given sample.

Based on PSE per gene, the Differential Shannon Diversity (DSD) was calculated as the log2 fold change of PSE values in the case sample over PSE value in the control sample. Thereby, it allows DSD to be directly compared to DEG values, which is defined by the log2 fold change of the gene expression intensity. Finally, the PSE was calculated for all the genes across all the samples. Per gene, the average PSE was calculated across the RNA libraries with the same condition. Wilcox tests were performed for pairwise conditions.

### Cellular deconvolution analysis

Liver cellular composition was estimated using a reference-based deconvolution approach with 528 validated marker genes across 9 major liver cell types (hepatocytes, Kupffer cells, stellate cells, endothelial cells, T cells, B cells, NK cells, neutrophils, and monocytes; supplemental Table S1). Cell type proportions were calculated for each sample and expressed as percentages of total liver cell composition. Statistical comparisons across disease groups were performed using one-way ANOVA followed by post-hoc analysis, with significance set at p < 0.05.

### Bioinformatics Pathway Analyses

Pathway analyses were conducted using Ingenuity Pathway Analysis (IPA) software from Qiagen in Valencia, CA (http://www.ingenuity.com). This software was utilized for canonical pathway analysis and network discovery. IPA’s core analyses rely on existing knowledge of the relationships between upstream regulators and their downstream target genes, which are stored in the Ingenuity Knowledge Base. Fisher’s exact test was employed to calculate p-values for the analysis. Ingenuity pathway analysis (IPA; QIAGEN) was used to identify top canonical and enriched biological pathways determined by DEG and DSD approaches. The cutoff for Log2 Fold change values was assigned >=1 and <=-1 for the upregulated and downregulated genes respectively. Log p-value >=1.3 was selected for the study as the level of significance.

### Data processing and statistical analysis

For the DEG and DSD analysis the Wilcox tests were performed. Spearman’s rank correlation test was employed to assess correlations between gene expression and prevalence rank. All statistical analyses were conducted, and graphs were generated using GraphPad Prism software from GraphPad in La Jolla, CA, USA.

### Study approval

This study involved secondary analysis of existing sequencing data obtained from the Database of Genotypes and Phenotypes (dbGAP, phs001807.v1.p1) maintained by the National Institutes of Health (NIH, USA). As this research utilized only de-identified, publicly available data, institutional review board (IRB) approval was not required. The original studies contributing data to dbGAP received appropriate IRB approval from their respective institutions prior to data deposition.

## Data availability statement

Data available upon request.

## Authors’ contributions

**Sudrishti Chaudhary:** Conceptualization, Methodology, Formal analysis, Investigation, Writing – original draft, Visualization. **Jia-Jun Liu:** Formal analysis, Software, Writing – original draft, Writing – review & editing. **Silvia Liu:** Formal analysis, Software, Supervision, Writing – original draft, Writing – review & editing. **Marissa Di**: Formal analysis, Software**. Juliane I Beier:** Writing – original draft, Writing – review & editing. **Ramon Bataller:** Resources, Data curation, Funding acquisition, Writing – review & editing. **Josepmaria Argemi:** Resources, Data curation, Writing – review & editing. **Panayiotis V Benos:** Formal analysis, Software, Methodology, Writing – review & editing. **Gavin E Arteel:** Conceptualization, Supervision, Project administration, Funding acquisition, Writing – original draft, Writing – review & editing.

## Funding support

This study was supported in part, by grants from NIH (R01 DK130294, R01 AA028436, P30 DK120531, R01 HL159805).

| Institution | Grant Number |
| --- | --- |
| NIH grant awarded by the National Institute of Diabetes and Digestive and Kidney Diseases (NIDDK) | R01<br>DK130294 |
| National Institute on Alcohol Abuse & Alcoholism | R01<br>AA028436 |
| Center Core Grant from the National Institute of Diabetes and Digestive and Kidney Diseases (NIDDK) that funds the Pittsburgh Liver Research Center (PLRC) at the University of Pittsburgh. | P30<br>DK120531 |
| National Heart, Lung, and Blood Institute (NHLBI) | R01<br>HL159805) |

## Acknowledgments

We acknowledge the use of data from the Database of Genotypes and Phenotypes (dbGaP, accession phs001807.v1.p1). We thank the consortium led by Principal Investigator Ramon Bataller (University of Pittsburgh) and all contributing members from Institut national de la santé et de la recherche médicale (France), Cliniques Universitaires Saint-Luc (Belgium), Institut d’Investigacions Biomèdiques August Pi i Sunyer (Spain), and Hospital Clinic, University of Barcelona (Spain). The original study was funded by the National Institute on Alcohol Abuse and Alcoholism (NIAAA), NIH, Bethesda, MD, USA.

