## Supplemental material for "Alpha diversity analysis of hepatic transcriptome reveals novel pathways in alcohol-related hepatitis"

^1^Department of Medicine, Division of Gastroenterology, Hepatology and Nutrition, ^2^Pharmacology and Chemical Biology, University of Pittsburgh, **^3^**Organ Pathobiology and therapeutics institute, University of Pittsburgh, ^4^Pittsburgh Liver Research Center, University of Pittsburgh. ^5^Department of Epidemiology, University of Florida, Gainesville, Florida, USA. ^6^Institut d’Investigacions Biomediques August Pi i Sunyer (IDIBAPS), University of Barcelona, Barcelona, Spain, ^7^Department of Internal Medicine, Liver Unit, Clinical University of Navarra, Navarra, Spain.

Key Words: Alpha diversity, Hepatic transcriptome., EtOH, Liver disease

Send all correspondence to:	Gavin E. Arteel, PhD, FAASLD

Thomas E. Starzl Biomedical Science Tower

West 1143

200 Lothrop Street

Pittsburgh, PA 15213

Abbreviations: DEG, differentially expressed genes, DSD; Differential Shannon Diversity; ALD, alcohol-related liver disease; ASH; alcohol-related steatohepatitis, AHR; alcohol-related hepatitis responders, AHNR; alcohol-related hepatitis non-responders.

**Supplemental results**

The supplement table S1 represents the liver cellular composition estimated using a reference-based deconvolution approach with 528 validated marker genes across 9 major liver cell types in ALD disease progression. The supplement table S2-S10 lists the significant canonical pathways enriched across all the groups. The tables S2, S3 and S4 represents top canonical pathways enriched in ASH/HC group exclusively by DEG, DSD and commonly by both DEG and DSD approaches respectively. The tables S5, S6 and S7 represents top canonical pathways enriched in AH/HC group exclusively by DEG, DSD and commonly by both DEG and DSD approaches respectively. The tables S8, S9 and S10 represents top canonical pathways enriched in AH/ASH group exclusively by DEG, DSD and commonly by both DEG and DSD approaches respectively.

Supplemental Tables

Table S1: Liver cell type marker genes.

| Cell type | Number of genes^1^ | Genes employed.^1^ |
| --- | --- | --- |
| Hepatocytes | 57 | ALB, APOB, CYP3A4, CYP2E1, CYP1A2, SERPINA1, TF, AFP, HNF4A, CYP2C9, CYP1A1, CYP2A6, CYP2B6, CYP2C8, CYP2C19, CYP2D6, CYP3A5, CYP3A7, G6PC, PCK1, CPS1, FABP1, APOA1, APOA2, APOA4, APOC1, APOC2, APOC3, SLCO1B1, SLCO1B3, ABCB11, ABCC2, SLC22A1, SLC22A7, HNF1A, HNF1B, FOXA2, FOXA3, CEBPA, CEBPB, ONECUT1, ONECUT2, TAT, HPX, FGA, FGB, FGG, AHSG, RBP4, IGF1, ANGPTL3, FTL, AGT, C3, CFB, ORM1, ORM2 |
| Kupffer Cells | 58 | CD68, CD163, CSF1R, MARCO, MSR1, CD14, LYZ, ITGAM, ADGRE1, FCGR1A, CLEC4F, TIMD4, VSIG4, CD5L, HMOX1, SPIC, ID3, CLEC2D, C1QA, C1QB, C1QC, C3, C4A, C4B, CFB, CFD, CFH, CFI, IL1B, IL6, TNF, IL10, IL12A, IL12B, CXCL10, CCL2, CCL3, MERTK, AXL, TYRO3, FCGR2A, FCGR2B, FCGR3A, SCARA1, SCARA5, IRF4, IRF5, IRF8, STAT1, NFKB1, TYROBP, TREM2, CX3CR1, P2RY12, SIGLEC1, LILRB4, CD206, MRC1 |
| Stellate Cells | 59 | ACTA2, COL1A1, PDGFRB, TIMP1, MMP2, LRAT, RBP1, TAGLN, CNN1, PDGFRA, COL1A2, COL3A1, COL4A1, COL4A2, COL5A1, COL5A2, COL6A1, COL6A2, COL6A3, MMP1, MMP3, MMP9, MMP13, MMP14, TIMP2, TIMP3, TIMP4, SMTN, MYH11, CALD1, DES, VIM, MSN, TGFB1, TGFBR1, TGFBR2, SMAD2, SMAD3, SMAD4, CTGF, NOX4, PLIN2, PLIN3, ADIPOQ, LEP, PPARG, CEBPA, FABP4, FASN, GFAP, NESTIN, THY1, NT5E, ENG, PECAM1, MCAM, HAND2, TWIST1, SNAI1 |
| Endothelial Cells | 59 | PECAM1, VWF, CDH5, KDR, FLT1, ICAM1, VCAM1, TEK, ENG, PLVAP, CLDN5, OCLN, TJP1, TJP2, JAM2, JAM3, ESAM, NOS3, EDN1, EDNRA, EDNRB, ANGPT1, ANGPT2, TIE1, ROBO4, LYVE1, STAB1, STAB2, CLEC14A, FCN2, FCN3, CLEC4G, CLEC4M, VEGFA, VEGFB, VEGFC, VEGFD, FLT4, NRP1, NRP2, PDGFA, PDGFB, CD34, CD31, MCAM, THY1, PROCR, THBD, F8, SERPINE1, ADAMTS13, COL4A1, EMCN, ERG, FLI1, SOX17, SOX18, EPAS1, HEY1 |
| T cells | 61 | CD3D, CD3E, CD3G, CD8A, CD4, IL2RB, GZMB, PRF1, CD2, CD5, TRAC, TRBC1, TRBC2, TRGC1, TRGC2, TRDC, CD247, LCK, ZAP70, IL2, IL4, IL5, IL13, IL17A, IL17F, IL21, IL22, IFNG, TNF, GZMA, GZMH, GZMK, GZMM, GNLY, FASLG, TNFSF10, CCL5, CD25, CD69, CD71, CD95, ICOS, CTLA4, PDCD1, LAG3, TIGIT, HAVCR2, CCR7, SELL, CD127, KLRG1, CX3CR1, CD57, CXCR3, CCR4, CCR6, CXCR5, BCL6, FOXP3, RORC, TBX21 |
| B cells | 59 | CD19, MS4A1, CD79A, CD79B, IGHM, IGHA1, IGHG1, CD20, PAX5, BLNK, IGHG2, IGHG3, IGHG4, IGHA2, IGHD, IGHE, IGKC, IGLC1, IGLC2, IGLC3, EBF1, TCF3, IKZF1, IKZF3, IRF4, IRF8, PRDM1, XBP1, BCL6, AICDA, CD21, CD23, CD27, CD38, CD138, TNFRSF17, TNFRSF13B, TNFRSF13C, SDC1, JCHAIN, MZB1, DERL3, FKBP11, CR2, FCER2, FCMR, FCRL1, FCRL2, FCRL3, FCRL4, FCRL5, CD22, SIGLEC2, BANK1, SPIB, MEF2C, BACH2, IL4R, CXCR4 |
| NK cells | 57 | KLRB1, NCR1, FCGR3A, GNLY, PRF1, GZMH, KLRD1, KLRF1, NKG7, NCAM1, KIR2DL1, KIR2DL2, KIR2DL3, KIR2DL4, KIR3DL1, KIR3DL2, KIR3DL3, KIR2DS1, KIR2DS2, KIR2DS3, KIR2DS4, KIR2DS5, KIR3DS1, NCR2, NCR3, KLRK1, CD226, TIGIT, DNAM1, GZMA, GZMB, GZMK, GZMM, FASLG, TNFSF10, CCL3, CCL4, CCL5, CD56, CD16, CD57, CD69, CD25, KLRC1, KLRC2, KLRC3, KLRC4, IL2RB, IL15R, IL12RB2, IFNG, TNF, CSF2, XCL1, XCL2, CXCR1, S1PR5 |
| Neutrophils | 58 | FCGR3B, CSF3R, CEACAM8, S100A12, ELANE, MPO, CAMP, LTF, DEFA1, DEFA3, DEFA4, AZU1, CTSG, PRTN3, BPI, LCN2, MMP8, MMP9, OLFM4, CXCR1, CXCR2, CXCR4, CCR1, CCR2, CCR3, CCR5, C5AR1, ITGAM, ITGB2, SELL, SELPLG, PADI4, HIST1H2A, HIST1H2B, HIST1H3A, HIST1H4A, G0S2, MNDA, ANXA3, S100A8, S100A9, FFAR2, ALPL, CREB5, GFI1, SPI1, CEBPE, LEF1, NAMPT, VNN1, RETN, TCN1, HP, CEACAM1, CEACAM6, ARG1, IL1R2, TNFAIP6 |
| Monocytes | 60 | CD14, FCGR1A, S100A8, S100A9, LYZ, VCAN, CCR2, CX3CR1, CSF1R, IRF8, FCGR3A, CCR5, CCR7, SELL, ITGAM, ITGAX, TLR2, TLR4, TLR8, CD163, MAFB, MAF, NR4A1, NR4A2, NR4A3, EGR1, EGR2, FOS, JUN, IL1B, IL6, TNF, CCL2, CCL3, CCL4, CXCL10, MNDA, SIGLEC1, MARCO, MSR1, SCARB1, SPI1, IRF4, KLF4, MAFB, CD86, HLA-DRA, HLA-DRB1, HLA-DQA1, HLA-DQB1, CD68, LILRB2, LILRB4, CLEC7A, CLEC4E, CLEC10A, EMR1, VSIG4, FOLR2, STAB1 |

^1^Total Markers: 528. Unique Markers: 458. Available in Dataset: 496 (93.9%) Average per Cell Type: 58.7 genes.

Table S2: The top canonical pathways enriched in ASH/HC by DEG

| Ingenuity Canonical Pathways | log(p-value) | Ratio | z-score | count |
| --- | --- | --- | --- | --- |
| Binding and Uptake of Ligands by Scavenger Receptors | 8.67 | 0.105 | -3.464 | 12 |
| Complement cascade | 5.85 | 0.0735 | -3.162 | 10 |
| Cell surface interactions at the vascular wall | 4.87 | 0.0514 | -3.317 | 11 |
| Immunoregulatory interactions between a Lymphoid and a non-Lymphoid cell | 4.85 | 0.0512 | -2.714 | 11 |
| Receptor-type tyrosine-protein phosphatases | 4.35 | 0.2 | 1 | 4 |
| Fcgamma receptor (FCGR) dependent phagocytosis | 4.31 | 0.0549 | -3 | 9 |
| Fc epsilon receptor (FCERI) signaling | 3.55 | 0.0437 | -3 | 9 |
| Signaling by the B Cell Receptor (BCR) | 3.43 | 0.0471 | -2.828 | 8 |
| LXR/RXR Activation | 3.42 | 0.0538 | -1.633 | 7 |
| Mevalonate Pathway I | 3.28 | 0.188 | - | 3 |
| DHCR24 Signaling Pathway | 3.17 | 0.049 | -1.89 | 7 |
| Intrinsic Prothrombin Activation Pathway | 3.02 | 0.093 | -2 | 4 |
| Superpathway of Geranylgeranyldiphosphate Biosynthesis I (via Mevalonate) | 2.98 | 0.15 | - | 3 |
| B Cell Development | 2.77 | 0.0265 | - | 13 |
| Pathogen Induced Cytokine Storm Signaling Pathway | 2.7 | 0.0288 | -1.508 | 11 |
| B Cell Receptor Signaling | 2.61 | 0.0235 | - | 15 |
| Cholesterol biosynthesis | 2.59 | 0.111 | - | 3 |
| IL-15 Signaling | 2.5 | 0.0247 | - | 13 |
| FcγRIIB Signaling in B Lymphocytes | 2.46 | 0.0244 | - | 13 |
| Systemic Lupus Erythematosus in B Cell Signaling Pathway | 2.46 | 0.022 | - | 16 |
| Hepatic Fibrosis / Hepatic Stellate Cell Activation | 2.44 | 0.0365 | - | 7 |
| Superpathway of Cholesterol Biosynthesis | 2.42 | 0.0968 | - | 3 |
| Potassium Channels | 2.39 | 0.0485 | 0.447 | 5 |
| Response of EIF2AK4 (GCN2) to amino acid deficiency | 2.39 | 0.0485 | -2.236 | 5 |
| POU5F1 (OCT4), SOX2, NANOG repress genes related to differentiation | 2.36 | 0.2 | - | 2 |
| NAD biosynthesis II (from tryptophan) | 2.27 | 0.182 | - | 2 |
| TRIM21 Intracellular Antibody Signaling Pathway | 2.24 | 0.0229 | -3.606 | 13 |
| Vitamin D (calciferol) metabolism | 2.2 | 0.167 | - | 2 |
| Specification of primordial germ cells | 2.2 | 0.167 | - | 2 |
| Hematopoiesis from Multipotent Stem Cells | 2.2 | 0.167 | - | 2 |
| Systemic Lupus Erythematosus Signaling | 2.17 | 0.0187 | - | 20 |
| p70S6K Signaling | 2.16 | 0.0224 | - | 13 |
| Nonsense-Mediated Decay (NMD) | 2.15 | 0.0427 | -2.236 | 5 |
| PI3K Signaling in B Lymphocytes | 2.09 | 0.022 | - | 13 |
| Eukaryotic Translation Elongation | 2.08 | 0.041 | -2.236 | 5 |
| Eukaryotic Translation Termination | 2.08 | 0.041 | -2.236 | 5 |
| Regulation of Insulin-like Growth Factor (IGF) transport and uptake by IGFBPs | 2.05 | 0.0403 | -2.236 | 5 |
| Activation of gene expression by SREBF (SREBP) | 2.05 | 0.0714 | - | 3 |
| GP6 Signaling Pathway | 1.99 | 0.0391 | -0.447 | 5 |
| Collagen chain trimerization | 1.99 | 0.0682 | - | 3 |
| Retinoid metabolism and transport | 1.99 | 0.0682 | - | 3 |
| Extrinsic Prothrombin Activation Pathway | 1.95 | 0.125 | - | 2 |
| Signaling by Retinoic Acid | 1.94 | 0.0652 | - | 3 |
| Interleukin-2 family signaling | 1.91 | 0.0638 | - | 3 |
| Cachexia Signaling Pathway | 1.87 | 0.0245 | -1 | 9 |
| Wound Healing Signaling Pathway | 1.85 | 0.0281 | -1.134 | 7 |
| SRP-dependent cotranslational protein targeting to membrane | 1.81 | 0.0352 | -2.236 | 5 |
| Communication between Innate and Adaptive Immune Cells | 1.78 | 0.0181 | -3.742 | 17 |
| Selenoamino acid metabolism | 1.77 | 0.0342 | -2.236 | 5 |
| Sensory processing of sound by outer hair cells of the cochlea | 1.73 | 0.0545 | - | 3 |
| Eukaryotic Translation Initiation | 1.72 | 0.0333 | -2.236 | 5 |
| Branched-chain amino acid catabolism | 1.72 | 0.0952 | - | 2 |
| Formation of the ureteric bud | 1.72 | 0.0952 | - | 2 |
| Interleukin-3, Interleukin-5 and GM-CSF signaling | 1.71 | 0.0536 | - | 3 |
| Regulation of TLR by endogenous ligand | 1.68 | 0.0909 | - | 2 |
| Extracellular matrix organization | 1.62 | 0.0374 | -2 | 4 |
| Transport of inorganic cations/anions and amino acids/oligopeptides | 1.61 | 0.037 | 0 | 4 |
| Coronavirus Replication Pathway | 1.55 | 0.0301 | - | 5 |
| Integrin signaling | 1.51 | 0.0741 | - | 2 |
| Collagen biosynthesis and modifying enzymes | 1.51 | 0.0448 | - | 3 |
| G-Protein Coupled Receptor Signaling | 1.5 | 0.0182 | -0.277 | 13 |
| Amyotrophic Lateral Sclerosis Signaling | 1.5 | 0.0342 | 0 | 4 |
| Regulation of lipid metabolism by PPARalpha | 1.47 | 0.0336 | 1 | 4 |
| EGR2 and SOX10-mediated initiation of Schwann cell myelination | 1.46 | 0.069 | - | 2 |
| Osteoarthritis Pathway | 1.45 | 0.0252 | -0.447 | 6 |
| Role of Osteoblasts in Rheumatoid Arthritis Signaling Pathway | 1.43 | 0.025 | -2.449 | 6 |
| cAMP-mediated signaling | 1.43 | 0.0249 | 0.816 | 6 |
| ID3 Signaling Pathway | 1.4 | 0.0165 | 2.5 | 16 |
| ERK5 Signaling | 1.4 | 0.0405 | - | 3 |
| NAD Biosynthesis III | 1.4 | 0.25 | - | 1 |
| Major pathway of rRNA processing in the nucleolus and cytosol | 1.37 | 0.0269 | -2.236 | 5 |
| Plasma lipoprotein assembly, remodeling, and clearance | 1.37 | 0.0395 | - | 3 |
| Airway Inflammation in Asthma | 1.35 | 0.0606 | - | 2 |
| TR/RXR Activation | 1.34 | 0.0305 | - | 4 |
| Maturity Onset Diabetes of Young (MODY) Signaling | 1.34 | 0.0385 | - | 3 |
| Response to elevated platelet cytosolic Ca2+ | 1.33 | 0.0303 | -2 | 4 |
| Acute Phase Response Signaling | 1.33 | 0.0262 | - | 5 |
| Cellular Effects of Sildenafil (Viagra) | 1.31 | 0.0175 | 0 | 12 |
| Coagulation System | 1.31 | 0.0571 | - | 2 |
| NAD Biosynthesis from 2-amino-3-carboxymuconate Semialdehyde | 1.3 | 0.2 | - | 1 |
| Protein Citrullination | 1.3 | 0.2 | - | 1 |
| Serotonin and Melatonin Biosynthesis | 1.3 | 0.2 | - | 1 |
| NAD Salvage Pathway III | 1.3 | 0.2 | - | 1 |
| Tyrosine Degradation I | 1.3 | 0.2 | - | 1 |

Table S3: The top canonical pathways enriched in ASH/HC by DSD

| Ingenuity Canonical Pathways | -log(p-value) | Ratio | z-score | COUNT |
| --- | --- | --- | --- | --- |
| Cell Cycle Checkpoints | 16 | 0.14 | 6.164 | 38 |
| Kinetochore Metaphase Signaling Pathway | 11.4 | 0.194 | 2 | 20 |
| Mitotic Prometaphase | 9.42 | 0.122 | 5 | 25 |
| RHO GTPases Activate Formins | 8.87 | 0.143 | 4.472 | 20 |
| Mitotic Metaphase and Anaphase | 6.87 | 0.0975 | 4.796 | 23 |
| Cohesin Chromatin Regulation Pathway | 6.61 | 0.0913 | 3.873 | 24 |
| Estrogen-mediated S-phase Entry | 6.55 | 0.308 | 2.828 | 8 |
| Mitotic G1 phase and G1/S transition | 6.22 | 0.121 | -1.807 | 16 |
| Nucleosome assembly | 6.2 | 0.204 | 3.162 | 10 |
| TR/RXR Activation | 5.56 | 0.115 | 3.207 | 15 |
| Glioblastoma Multiforme Signaling | 4.75 | 0.0936 | 3 | 16 |
| Ribonucleotide Reductase Signaling Pathway | 4.66 | 0.092 | 2.84 | 16 |
| Molecular Mechanisms of Cancer | 4.64 | 0.0523 | 4.323 | 45 |
| TP53 Regulates Transcription of Cell Cycle Genes | 4.33 | 0.163 | 2.828 | 8 |
| Replacement of protamines by nucleosomes in the male pronucleus | 4.28 | 0.189 | 2.646 | 7 |
| DNA methylation | 4.25 | 0.231 | 2.449 | 6 |
| HDR through Homologous Recombination (HRR) or Single Strand Annealing (SSA) | 4.2 | 0.105 | 3.464 | 12 |
| DNA Damage/Telomere Stress Induced Senescence | 3.9 | 0.143 | 2.828 | 8 |
| Erythropoietin Signaling Pathway | 3.89 | 0.0829 | -0.632 | 15 |
| Breast Cancer Regulation by Stathmin1 | 3.88 | 0.0545 | 4.596 | 33 |
| Oxidative Stress Induced Senescence | 3.87 | 0.114 | 3.162 | 10 |
| RUNX1 regulates megakaryocyte differentiation and platelet function | 3.79 | 0.138 | -0.707 | 8 |
| Role of CHK Proteins in Cell Cycle Checkpoint Control | 3.79 | 0.138 | -1.89 | 8 |
| Chromatin modifications during the maternal to zygotic transition (MZT) | 3.72 | 0.188 | 2.449 | 6 |
| Synaptic adhesion-like molecules | 3.69 | 0.238 | 2.236 | 5 |
| Kinesins | 3.59 | 0.129 | 2.828 | 8 |
| Prostate Cancer Signaling | 3.5 | 0.0948 | 2.53 | 11 |
| p53 Signaling | 3.49 | 0.102 | -0.333 | 10 |
| Small Cell Lung Cancer Signaling | 3.49 | 0.102 | - | 10 |
| Mitotic G2-G2/M phases | 3.42 | 0.075 | 3.873 | 15 |
| Granulocyte Adhesion and Diapedesis | 3.33 | 0.0735 | - | 15 |
| Activation of NMDA receptors and postsynaptic events | 3.31 | 0.106 | 3 | 9 |
| Cell Cycle Regulation by BTG Family Proteins | 3.29 | 0.158 | 1.633 | 6 |
| Docosahexaenoic Acid (DHA) Signaling | 3.28 | 0.0677 | 1.5 | 17 |
| Cyclins and Cell Cycle Regulation | 3.27 | 0.105 | 2.828 | 9 |
| Amyloid fiber formation | 3.2 | 0.102 | 3 | 9 |
| Neuroinflammation Signaling Pathway | 3.13 | 0.0606 | 4.123 | 20 |
| Activated PKN1 stimulates transcription of AR regulated genes KLK2 and KLK3 | 3.08 | 0.179 | 2.236 | 5 |
| WNT/SHH Axonal Guidance Signaling Pathway | 3.06 | 0.0795 | 2.887 | 12 |
| Interleukin-4 and Interleukin-13 signaling | 3.06 | 0.0901 | 1.265 | 10 |
| SIRT1 negatively regulates rRNA expression | 3 | 0.172 | 2.236 | 5 |
| TNFs bind their physiological receptors | 3 | 0.172 | 2.236 | 5 |
| Role of Pattern Recognition Receptors in Recognition of Bacteria and Viruses | 2.99 | 0.0779 | 2.828 | 12 |
| Pre-NOTCH Expression and Processing | 2.95 | 0.104 | 2.828 | 8 |
| Agranulocyte Adhesion and Diapedesis | 2.91 | 0.067 | - | 15 |
| Ovarian Cancer Signaling | 2.87 | 0.0755 | 2 | 12 |
| Reelin Signaling in Neurons | 2.87 | 0.0797 | 3.317 | 11 |
| Hereditary Breast Cancer Signaling | 2.87 | 0.0797 | 1.265 | 11 |
| DNA damage-induced 14-3-3σ Signaling | 2.84 | 0.13 | 0.447 | 6 |
| Aryl Hydrocarbon Receptor Signaling | 2.82 | 0.071 | 2.111 | 13 |
| Degradation of the extracellular matrix | 2.8 | 0.0988 | 2.828 | 8 |
| ATM Signaling | 2.8 | 0.09 | 0.378 | 9 |
| Chronic Myeloid Leukemia Signaling | 2.74 | 0.0605 | 3.638 | 17 |
| Potassium Channels | 2.71 | 0.0874 | 3 | 9 |
| Autism Signaling Pathway | 2.69 | 0.0583 | 3.771 | 18 |
| Sertoli Cell-Germ Cell Junction Signaling Pathway (Enhanced) | 2.69 | 0.0636 | -1.291 | 15 |
| DNA Replication Pre-Initiation | 2.68 | 0.0865 | 3 | 9 |
| Pancreatic Adenocarcinoma Signaling | 2.67 | 0.08 | 2 | 10 |
| Mitotic Roles of Polo-Like Kinase | 2.66 | 0.104 | 1.633 | 7 |
| PRC2 methylates histones and DNA | 2.62 | 0.143 | 2.236 | 5 |
| IL-9 Signaling | 2.62 | 0.143 | 1 | 5 |
| Transcriptional regulation of granulopoiesis | 2.61 | 0.118 | 1.633 | 6 |
| Cell Cycle: G2/M DNA Damage Checkpoint Regulation | 2.61 | 0.118 | -1 | 6 |
| Glycosaminoglycan metabolism | 2.59 | 0.0781 | 3.162 | 10 |
| Cell Cycle: G1/S Checkpoint Regulation | 2.58 | 0.101 | -2.646 | 7 |
| B-WICH complex positively regulates rRNA expression | 2.57 | 0.115 | 2.449 | 6 |
| NAP1L1 Transcription Regulation Signaling Pathway | 2.54 | 0.0899 | 2.121 | 8 |
| CDX Gastrointestinal Cancer Signaling Pathway | 2.54 | 0.066 | -2.887 | 13 |
| ERCC6 (CSB) and EHMT2 (G9a) positively regulate rRNA expression | 2.52 | 0.135 | 2.236 | 5 |
| Fatty acyl-CoA biosynthesis | 2.52 | 0.135 | 2.236 | 5 |
| Differential Regulation of Cytokine Production in Intestinal Epithelial Cells by IL-17A and IL-17F | 2.5 | 0.174 | 2 | 4 |
| Sertoli Cell-Sertoli Cell Junction Signaling | 2.5 | 0.0607 | 2.324 | 15 |
| Glycation Signaling Pathway | 2.46 | 0.0622 | 2.138 | 14 |
| Cell junction organization | 2.46 | 0.087 | 2.828 | 8 |
| ID1 Signaling Pathway | 2.45 | 0.0644 | 1.387 | 13 |
| Meiotic synapsis | 2.44 | 0.109 | 2.449 | 6 |
| Senescence Pathway | 2.43 | 0.0565 | -1.069 | 17 |
| Regulation of TP53 Activity through Phosphorylation | 2.43 | 0.086 | 2.828 | 8 |
| Senescence-Associated Secretory Phenotype (SASP) | 2.41 | 0.0946 | 2.646 | 7 |
| Sheddase Signaling Pathway | 2.41 | 0.0637 | 3.606 | 13 |
| Class A/1 (Rhodopsin-like receptors) | 2.36 | 0.0542 | 4.243 | 18 |
| Role of PKR in Interferon Induction and Antiviral Response | 2.36 | 0.0725 | 1.89 | 10 |
| Preeclampsia Signaling Pathway | 2.34 | 0.0683 | -0.905 | 11 |
| Lung Ionic Balance Signaling Pathway | 2.34 | 0.0479 | 4.6 | 25 |
| Triacylglycerol Biosynthesis | 2.33 | 0.103 | 2.236 | 6 |
| RET signaling | 2.32 | 0.122 | 2.236 | 5 |
| Hematoma Resolution Signaling Pathway | 2.31 | 0.0579 | -1.291 | 15 |
| Coronavirus Pathogenesis Pathway | 2.29 | 0.0616 | -0.832 | 13 |
| DNA Double Strand Break Response | 2.29 | 0.102 | 2.449 | 6 |
| Activation of gene expression by SREBF (SREBP) | 2.27 | 0.119 | 2.236 | 5 |
| GADD45 Signaling | 2.25 | 0.1 | -2.236 | 6 |
| Regulation of TP53 Activity through Association with Co-factors | 2.24 | 0.214 | - | 3 |
| Signaling by SCF-KIT | 2.23 | 0.116 | 2.236 | 5 |
| Role of BRCA1 in DNA Damage Response | 2.23 | 0.0875 | -0.378 | 7 |
| Mitotic Telophase/Cytokinesis | 2.15 | 0.2 | - | 3 |
| Stearate Biosynthesis I (Animals) | 2.15 | 0.0952 | 2.236 | 6 |
| Tuberculosis Active Signaling Pathway | 2.14 | 0.0571 | 0.277 | 14 |
| UFMylation Signaling Pathway | 2.12 | 0.0938 | 0 | 6 |
| Activation of anterior HOX genes in hindbrain during early embryogenesis | 2.11 | 0.0833 | 2.646 | 7 |
| Apelin Pancreas Signaling Pathway | 2.1 | 0.109 | - | 5 |
| IL-23 Signaling Pathway | 2.1 | 0.109 | 2 | 5 |
| Sleep REM Signaling Pathway | 2.09 | 0.0755 | -1.414 | 8 |
| Cyclophilin Signaling Pathway | 2.08 | 0.0562 | 2.673 | 14 |
| S100 Family Signaling Pathway | 2.07 | 0.0422 | 5.396 | 33 |
| Pathogen Induced Cytokine Storm Signaling Pathway | 2.07 | 0.0497 | 4.123 | 19 |
| Transcriptional regulation by RUNX1 | 2.07 | 0.0658 | 3.162 | 10 |
| Type II Diabetes Mellitus Signaling | 2.07 | 0.0658 | 2.236 | 10 |
| BBSome Signaling Pathway | 2.07 | 0.0467 | 3.838 | 23 |
| Sonic Hedgehog Signaling | 2.03 | 0.129 | - | 4 |
| NoRC negatively regulates rRNA expression | 2.02 | 0.0896 | 2.449 | 6 |
| IL-10 Signaling | 2.01 | 0.0645 | 0 | 10 |
| Trehalose Degradation II (Trehalase) | 1.99 | 0.333 | - | 2 |
| Nonhomologous End-Joining (NHEJ) | 1.99 | 0.102 | 2.236 | 5 |
| Telomere Maintenance | 1.98 | 0.0787 | 2.646 | 7 |
| Meiotic recombination | 1.95 | 0.1 | 2.236 | 5 |
| Atherosclerosis Signaling | 1.93 | 0.0662 | 2.828 | 9 |
| HMGB1 Signaling | 1.92 | 0.0625 | 2.449 | 10 |
| Transcriptional regulation by RUNX2 | 1.92 | 0.098 | 1.342 | 5 |
| G-Protein Coupled Receptor Signaling | 1.92 | 0.0421 | 4.017 | 30 |
| Basal Cell Carcinoma Signaling | 1.9 | 0.0845 | 1 | 6 |
| Synaptogenesis Signaling Pathway | 1.89 | 0.0506 | 3.5 | 16 |
| CGAS-STING Signaling Pathway | 1.89 | 0.0652 | 3 | 9 |
| Hepatitis B Chronic Liver Pathogenesis Signaling Pathway | 1.86 | 0.0585 | 1.265 | 11 |
| Mitochondrial Uncoupling | 1.85 | 0.286 | - | 2 |
| Salvage Pathways of Pyrimidine Deoxyribonucleotides | 1.85 | 0.286 | - | 2 |
| Role of Osteoblasts in Rheumatoid Arthritis Signaling Pathway | 1.85 | 0.0542 | 0.277 | 13 |
| Activation of the pre-replicative complex | 1.84 | 0.114 | 2 | 4 |
| HOTAIR Regulatory Pathway | 1.83 | 0.0606 | 2.828 | 10 |
| ESR-mediated signaling | 1.82 | 0.0678 | 2.828 | 8 |
| RNA Polymerase I Transcription | 1.82 | 0.0811 | 2.449 | 6 |
| Non-Small Cell Lung Cancer Signaling | 1.81 | 0.0729 | 2 | 7 |
| Colorectal Cancer Metastasis Signaling | 1.79 | 0.0517 | 3.051 | 14 |
| Autophagy | 1.78 | 0.0548 | 1.732 | 12 |
| Myelination Signaling Pathway | 1.75 | 0.0488 | 4 | 16 |
| Nuclear Cytoskeleton Signaling Pathway | 1.75 | 0.0543 | 3.317 | 12 |
| CREB Signaling in Neurons | 1.75 | 0.0422 | 4.6 | 26 |
| Synthesis of DNA | 1.74 | 0.0656 | 2.828 | 8 |
| Neurexins and neuroligins | 1.72 | 0.0877 | 2.236 | 5 |
| TCF dependent signaling in response to WNT | 1.71 | 0.0556 | 3.317 | 11 |
| Neuropathic Pain Signaling in Dorsal Horn Neurons | 1.7 | 0.0693 | 2.646 | 7 |
| Role of IL-17A in Arthritis | 1.69 | 0.0862 | - | 5 |
| Signaling by MET | 1.69 | 0.0759 | 2.449 | 6 |
| TREM1 Signaling | 1.69 | 0.0759 | 2.236 | 6 |
| Cellular hexose transport | 1.68 | 0.136 | - | 3 |
| Pulmonary Healing Signaling Pathway | 1.67 | 0.0547 | 2.53 | 11 |
| Role of Macrophages, Fibroblasts and Endothelial Cells in Rheumatoid Arthritis | 1.66 | 0.0475 | 3.5 | 16 |
| Mitotic Prophase | 1.64 | 0.0673 | 2.646 | 7 |
| GDP-glucose Biosynthesis | 1.63 | 0.222 | - | 2 |
| Glioma Signaling | 1.63 | 0.0625 | - | 8 |
| IL-6 Signaling | 1.61 | 0.062 | 2.236 | 8 |
| Estrogen-Dependent Breast Cancer Signaling | 1.6 | 0.0723 | 2.236 | 6 |
| Tumor Microenvironment Pathway | 1.6 | 0.0556 | 3 | 10 |
| Tumoricidal Function of Hepatic Natural Killer Cells | 1.58 | 0.125 | - | 3 |
| Extracellular matrix organization | 1.58 | 0.0654 | 2.646 | 7 |
| Transport of bile salts and organic acids, metal ions and amine compounds | 1.58 | 0.0714 | 2.449 | 6 |
| PEDF Signaling | 1.58 | 0.0714 | 2 | 6 |
| Irritable Bowel Syndrome Signaling Pathway | 1.57 | 0.0484 | 2.673 | 14 |
| Acetyl-CoA Biosynthesis III (from Citrate) | 1.57 | 1 | - | 1 |
| 14-3-3-mediated Signaling | 1.56 | 0.0606 | 0.447 | 8 |
| Transport of inorganic cations/anions and amino acids/oligopeptides | 1.56 | 0.0648 | 2.646 | 7 |
| Chondroitin Sulfate Biosynthesis (Late Stages) | 1.56 | 0.0794 | 2 | 5 |
| Neurotransmitter clearance | 1.55 | 0.2 | - | 2 |
| mRNA Editing | 1.55 | 0.2 | - | 2 |
| Glucose and Glucose-1-phosphate Degradation | 1.55 | 0.2 | - | 2 |
| Regulation of the Epithelial Mesenchymal Transition in Development Pathway | 1.53 | 0.0698 | 2.236 | 6 |
| TP53 Regulates Transcription of Cell Death Genes | 1.51 | 0.0909 | 2 | 4 |
| Assembly and cell surface presentation of NMDA receptors | 1.51 | 0.0909 | 2 | 4 |
| HEY1 Signaling Pathway | 1.5 | 0.0559 | 3 | 9 |
| Regulation of mitotic cell cycle | 1.49 | 0.0682 | 2.449 | 6 |
| Sirtuin Signaling Pathway | 1.49 | 0.0471 | -1.414 | 14 |
| Interleukin-10 signaling | 1.48 | 0.0889 | 2 | 4 |
| Pulmonary Fibrosis Idiopathic Signaling Pathway | 1.48 | 0.046 | 3.873 | 15 |
| Activation of IRF by Cytosolic Pattern Recognition Receptors | 1.48 | 0.0758 | 1 | 5 |
| Induction of Apoptosis by HIV1 | 1.48 | 0.0758 | 1 | 5 |
| IL-17 Signaling | 1.47 | 0.0529 | 3.162 | 10 |
| CD40 Signaling | 1.46 | 0.0746 | 1 | 5 |
| Tuberculosis Latent Signaling Pathway | 1.45 | 0.0667 | -1.633 | 6 |
| Role of IL-17F in Allergic Inflammatory Airway Diseases | 1.45 | 0.087 | - | 4 |
| Hepatic Fibrosis Signaling Pathway | 1.45 | 0.0433 | 2.324 | 18 |
| Pyroptosis | 1.45 | 0.111 | - | 3 |
| Macrophage Alternative Activation Signaling Pathway | 1.43 | 0.0521 | 1.667 | 10 |
| Ephrin A Signaling | 1.43 | 0.0571 | 2.121 | 8 |
| IL-17A Signaling in Airway Cells | 1.43 | 0.0735 | - | 5 |
| Ceramide Signaling | 1.43 | 0.0659 | 0 | 6 |
| Wound Healing Signaling Pathway | 1.4 | 0.0482 | 1.155 | 12 |
| IL-33 Signaling Pathway | 1.4 | 0.0513 | 2.121 | 10 |
| FXR/RXR Activation | 1.4 | 0.0513 | -1.667 | 10 |
| Peptide hormone biosynthesis | 1.39 | 0.167 | - | 2 |
| GP1b-IX-V activation signalling | 1.39 | 0.167 | - | 2 |
| HDR through MMEJ (alt-NHEJ) | 1.39 | 0.167 | - | 2 |
| UDP-N-acetyl-D-galactosamine Biosynthesis II | 1.39 | 0.167 | - | 2 |
| Role of p14/p19ARF in Tumor Suppression | 1.37 | 0.103 | - | 3 |
| PFKFB4 Signaling Pathway | 1.36 | 0.0816 | - | 4 |
| Glioma Invasiveness Signaling | 1.36 | 0.0704 | 2.236 | 5 |
| Chondroitin Sulfate Biosynthesis | 1.36 | 0.0704 | 2 | 5 |
| Hepatic Cholestasis | 1.36 | 0.0489 | 2.53 | 11 |
| Chromatin organization | 1.34 | 0.0471 | 3.464 | 12 |
| Melanoma Signaling | 1.34 | 0.08 | 2 | 4 |
| MYC Mediated Apoptosis Signaling | 1.34 | 0.08 | 2 | 4 |
| WNT/β-catenin Signaling | 1.33 | 0.052 | 0 | 9 |
| Advanced glycosylation endproduct receptor signaling | 1.33 | 0.154 | - | 2 |
| Endocannabinoid Cancer Inhibition Pathway | 1.33 | 0.0544 | -2.646 | 8 |
| IL-15 Production | 1.32 | 0.0574 | 2.449 | 7 |
| FAT10 Cancer Signaling Pathway | 1.31 | 0.0784 | - | 4 |
| RAR Activation | 1.31 | 0.0416 | 0.943 | 18 |
| IL-13 Signaling Pathway | 1.3 | 0.0569 | 1.633 | 7 |
| Interferon gamma signaling | 1.3 | 0.0612 | 2.449 | 6 |
| DNA Methylation and Transcriptional Repression Signaling | 1.3 | 0.0612 | 2.449 | 6 |
| Signaling by ALK | 1.3 | 0.0968 | - | 3 |

Table S4: The top common pathways enriched in ASH/HC by DEG and DSD

| Ingenuity Canonical Pathways | -log(p-value) | Ratio | z-score | Gene count |
| --- | --- | --- | --- | --- |
| Binding and Uptake of Ligands by Scavenger Receptors | 32.4 | 0.377 | -6.557 | 43 |
| Complement cascade | 27.6 | 0.309 | -6.481 | 42 |
| Fcgamma receptor (FCGR) dependent phagocytosis | 25 | 0.262 | -6.557 | 43 |
| Cell surface interactions at the vascular wall | 22.7 | 0.215 | -6.193 | 46 |
| Immunoregulatory interactions between a Lymphoid and a non-Lymphoid cell | 22.6 | 0.214 | -6.193 | 46 |
| Fc epsilon receptor (FCERI) signaling | 22.5 | 0.218 | -6.557 | 45 |
| Signaling by the B Cell Receptor (BCR) | 20.4 | 0.229 | -6.245 | 39 |
| B Cell Development | 17.7 | 0.126 | - | 62 |
| PI3K Signaling in B Lymphocytes | 17.2 | 0.115 | -2.646 | 68 |
| Systemic Lupus Erythematosus in B Cell Signaling Pathway | 15.7 | 0.102 | -2.183 | 74 |
| IL-15 Signaling | 15.6 | 0.116 | - | 61 |
| B Cell Receptor Signaling | 15 | 0.105 | -1.633 | 67 |
| FcγRIIB Signaling in B Lymphocytes | 14.8 | 0.113 | - | 60 |
| p70S6K Signaling | 14.8 | 0.108 | -2 | 63 |
| TRIM21 Intracellular Antibody Signaling Pathway | 14.7 | 0.109 | -7.62 | 62 |
| Metallothioneins bind metals | 9.49 | 0.727 | -2.828 | 8 |
| Communication between Innate and Adaptive Immune Cells | 8.6 | 0.0744 | -7.814 | 70 |
| Altered T Cell and B Cell Signaling in Rheumatoid Arthritis | 8.19 | 0.0732 | - | 69 |
| ID3 Signaling Pathway | 8.11 | 0.0724 | 6.693 | 70 |
| Role of NFAT in Regulation of the Immune Response | 7.42 | 0.0688 | -1.89 | 72 |
| Systemic Lupus Erythematosus Signaling | 7.08 | 0.0674 | - | 72 |
| Mitochondrial RNA degradation | 7.04 | 0.36 | 3 | 9 |
| Role of Macrophages, Fibroblasts and Endothelial Cells in Rheumatoid Arthritis | 6.94 | 0.0979 | -3.413 | 33 |
| Primary Immunodeficiency Signaling | 6.74 | 0.21 | - | 13 |
| Phospholipase C Signaling | 6.24 | 0.0641 | -3 | 72 |
| rRNA processing | 6 | 0.281 | 3 | 9 |
| Macrophage Alternative Activation Signaling Pathway | 5.96 | 0.115 | -3.578 | 22 |
| ABRA Signaling Pathway | 5.67 | 0.159 | -2.673 | 14 |
| Activin Inhibin Signaling Pathway | 5.25 | 0.104 | -3.411 | 22 |
| S100 Family Signaling Pathway | 5.1 | 0.0665 | -5.181 | 52 |
| tRNA processing in the mitochondrion | 5.02 | 0.22 | 3 | 9 |
| Phagosome Formation | 4.37 | 0.0652 | -5.466 | 46 |
| Neutrophil Extracellular Trap Signaling Pathway | 4.29 | 0.0758 | -1.095 | 31 |
| Cardiac conduction | 3.73 | 0.108 | -2.333 | 14 |
| Glycation Signaling Pathway | 3.43 | 0.0844 | -2.524 | 19 |
| VDR/RXR Activation | 3.42 | 0.128 | 1.633 | 10 |
| Hepatic Fibrosis / Hepatic Stellate Cell Activation | 3.36 | 0.0885 | - | 17 |
| Interleukin-4 and Interleukin-13 signaling | 3.29 | 0.108 | -2.887 | 12 |
| PXR/RXR Activation | 3.29 | 0.134 | 1.342 | 9 |
| SPINK1 General Cancer Pathway | 3.29 | 0.134 | 2.333 | 9 |
| FXR/RXR Activation | 3.29 | 0.0872 | 0.577 | 17 |
| Pulmonary Fibrosis Idiopathic Signaling Pathway | 3.28 | 0.0736 | -2.294 | 24 |
| p53 Signaling | 3.2 | 0.112 | 1.633 | 11 |
| Airway Pathology in Chronic Obstructive Pulmonary Disease | 3.19 | 0.105 | - | 12 |
| Interleukin-10 signaling | 3.05 | 0.156 | 0.378 | 7 |
| Agranulocyte Adhesion and Diapedesis | 3.03 | 0.0804 | - | 18 |
| Role of Chondrocytes in Rheumatoid Arthritis Signaling Pathway | 2.78 | 0.0903 | -2.496 | 13 |
| IL-17A Signaling in Gastric Cells | 2.74 | 0.192 | - | 5 |
| Transcriptional regulation by RUNX2 | 2.73 | 0.137 | 0.378 | 7 |
| Superpathway of Citrulline Metabolism | 2.72 | 0.25 | -2 | 4 |
| FLT3 Signaling | 2.66 | 0.154 | -2.449 | 6 |
| Cerebral Malformation Signaling Pathway | 2.57 | 0.094 | -0.905 | 11 |
| Potassium Channels | 2.49 | 0.0971 | -3.162 | 10 |
| Thyronamine and Iodothyronamine Metabolism | 2.45 | 0.667 | - | 2 |
| Thyroid Hormone Metabolism I (via Deiodination) | 2.45 | 0.667 | - | 2 |
| Smooth Muscle Contraction | 2.44 | 0.14 | -2.449 | 6 |
| Cellular Effects of Sildenafil (Viagra) | 2.44 | 0.0555 | 1.151 | 38 |
| FOXO-mediated transcription | 2.42 | 0.211 | -2 | 4 |
| Aspirin ADME | 2.39 | 0.136 | -2.449 | 6 |
| Sirtuin Signaling Pathway | 2.38 | 0.0673 | -3 | 20 |
| Post-translational protein phosphorylation | 2.37 | 0.0935 | -3.162 | 10 |
| HEY1 Signaling Pathway | 2.36 | 0.0807 | -1.155 | 13 |
| Multiple Sclerosis Signaling Pathway | 2.34 | 0.0731 | -1.604 | 16 |
| Ion channel transport | 2.29 | 0.0765 | -3.207 | 14 |
| RAR Activation | 2.26 | 0.06 | -1.961 | 26 |
| Phenylalanine and tyrosine metabolism | 2.25 | 0.273 | - | 3 |
| Thyroid Cancer Signaling | 2.23 | 0.103 | -1.414 | 8 |
| Hepatic Cholestasis | 2.23 | 0.0711 | -1.941 | 16 |
| IL-8 Signaling | 2.18 | 0.0721 | -2.496 | 15 |
| IL-17A Signaling in Fibroblasts | 2.17 | 0.1 | -1.414 | 8 |
| Glucocorticoid Receptor Signaling | 2.14 | 0.055 | - | 33 |
| Role of JAK2 in Hormone-like Cytokine Signaling | 2.13 | 0.108 | -1.342 | 7 |
| Pathogen Induced Cytokine Storm Signaling Pathway | 2.07 | 0.0602 | -2.683 | 23 |
| Leukocyte Extravasation Signaling | 2.07 | 0.0722 | -2.496 | 14 |
| Adipogenesis pathway | 2.04 | 0.0797 | 0.302 | 11 |
| Transcriptional regulation of testis differentiation | 2.03 | 0.231 | - | 3 |
| Hematoma Resolution Signaling Pathway | 2 | 0.0656 | 1.698 | 17 |
| CREB Signaling in Neurons | 1.98 | 0.0536 | -4.95 | 33 |
| Tumor Microenvironment Pathway | 1.97 | 0.0722 | -1.387 | 13 |
| Erythropoietin Signaling Pathway | 1.95 | 0.0718 | 0.277 | 13 |
| Pulmonary Healing Signaling Pathway | 1.95 | 0.0697 | -3.207 | 14 |
| IL-12 Signaling and Production in Macrophages | 1.94 | 0.0661 | 2.84 | 16 |
| Tyrosine Degradation I | 1.94 | 0.4 | - | 2 |
| Role of IL-17A in Psoriasis | 1.94 | 0.214 | - | 3 |
| Regulation of Insulin-like Growth Factor (IGF) transport and uptake by IGFBPs | 1.93 | 0.0806 | -3.162 | 10 |
| Bupropion Degradation | 1.92 | 0.154 | - | 4 |
| Neurotransmitter release cycle | 1.91 | 0.125 | -2.236 | 5 |
| Bone Mineralization Signaling Pathway | 1.91 | 0.0764 | 0.302 | 11 |
| Extracellular matrix organization | 1.9 | 0.0841 | -3 | 9 |
| Granulocyte Adhesion and Diapedesis | 1.89 | 0.0686 | - | 14 |
| G alpha (s) signalling events | 1.89 | 0.0759 | -2.714 | 11 |
| Irritable Bowel Syndrome Signaling Pathway | 1.87 | 0.0623 | -1.886 | 18 |
| Oxidative Phosphorylation | 1.87 | 0.0833 | 1.414 | 9 |
| Cell Cycle Control of Chromosomal Replication | 1.85 | 0.105 | 0 | 6 |
| Wound Healing Signaling Pathway | 1.84 | 0.0643 | -2.5 | 16 |
| Parkinson's Signaling Pathway | 1.8 | 0.0601 | -1.606 | 19 |
| Post-translational modification: synthesis of GPI-anchored proteins | 1.79 | 0.086 | -2.828 | 8 |
| Arginine Biosynthesis IV | 1.78 | 0.333 | - | 2 |
| FOXO-mediated transcription of cell death genes | 1.77 | 0.188 | - | 3 |
| Dilated Cardiomyopathy Signaling Pathway | 1.77 | 0.0728 | -0.333 | 11 |
| GADD45 Signaling | 1.74 | 0.1 | -0.816 | 6 |
| Netrin-1 signaling | 1.74 | 0.114 | -2.236 | 5 |
| Molecular Mechanisms of Cancer | 1.72 | 0.0488 | -4.629 | 42 |
| FOXO-mediated transcription of oxidative stress, metabolic and neuronal genes | 1.71 | 0.133 | -2 | 4 |
| FOXO-mediated transcription of cell cycle genes | 1.7 | 0.176 | - | 3 |
| IL-10 Signaling | 1.69 | 0.071 | 0 | 11 |
| IL-7 Signaling Pathway | 1.69 | 0.0886 | -2.236 | 7 |
| Atherosclerosis Signaling | 1.67 | 0.0735 | -1.134 | 10 |
| CDX Gastrointestinal Cancer Signaling Pathway | 1.67 | 0.066 | 0.277 | 13 |
| Role of Osteoblasts in Rheumatoid Arthritis Signaling Pathway | 1.66 | 0.0625 | -1.291 | 15 |
| Transcriptional Regulation by MECP2 | 1.65 | 0.0952 | -2.449 | 6 |
| Urea Cycle | 1.64 | 0.286 | - | 2 |
| Metabolism of amine-derived hormones | 1.63 | 0.167 | - | 3 |
| Estrogen Biosynthesis | 1.63 | 0.106 | - | 5 |
| Cardiac Hypertrophy Signaling (Enhanced) | 1.62 | 0.0521 | -3.138 | 28 |
| Bladder Cancer Signaling | 1.62 | 0.0756 | - | 9 |
| STAT3 Pathway | 1.62 | 0.0719 | -1.633 | 10 |
| Class A/1 (Rhodopsin-like receptors) | 1.6 | 0.0572 | -0.229 | 19 |
| Neuropathic Pain Signaling in Dorsal Horn Neurons | 1.6 | 0.0792 | -2.121 | 8 |
| NAFLD Signaling Pathway | 1.58 | 0.0625 | -1.604 | 14 |
| Gap Junction Signaling | 1.57 | 0.0567 | 0.243 | 19 |
| RHO GTPases activate CIT | 1.57 | 0.158 | - | 3 |
| RHO GTPases Activate ROCKs | 1.57 | 0.158 | - | 3 |
| IL-15 Production | 1.56 | 0.0738 | -2.333 | 9 |
| Sheddase Signaling Pathway | 1.56 | 0.0637 | 1.387 | 13 |
| Aryl Hydrocarbon Receptor Signaling | 1.56 | 0.0656 | - | 12 |
| NRF2-mediated Oxidative Stress Response | 1.55 | 0.0593 | 0.447 | 16 |
| Nicotine Degradation II | 1.54 | 0.0824 | - | 7 |
| rRNA modification in the mitochondrion | 1.53 | 0.25 | - | 2 |
| Histidine catabolism | 1.53 | 0.25 | - | 2 |
| Signaling by NOTCH2 | 1.52 | 0.118 | -2 | 4 |
| Role of Osteoclasts in Rheumatoid Arthritis Signaling Pathway | 1.52 | 0.0568 | -1.5 | 18 |
| Role of Osteoblasts, Osteoclasts and Chondrocytes in Rheumatoid Arthritis | 1.51 | 0.0611 | - | 14 |
| Receptor-type tyrosine-protein phosphatases | 1.51 | 0.15 | - | 3 |
| PIP3 activates AKT signaling | 1.51 | 0.069 | -2.53 | 10 |
| Sleep REM Signaling Pathway | 1.49 | 0.0755 | 0 | 8 |
| IL-17 Signaling | 1.47 | 0.0635 | -1.732 | 12 |
| Respiratory electron transport | 1.46 | 0.0795 | 1.134 | 7 |
| Melatonin Degradation III | 1.46 | 1 | - | 1 |
| Histamine Biosynthesis | 1.46 | 1 | - | 1 |
| 4-hydroxybenzoate Biosynthesis | 1.46 | 1 | - | 1 |
| 4-hydroxyphenylpyruvate Biosynthesis | 1.46 | 1 | - | 1 |
| ROBO SLIT Signaling Pathway | 1.45 | 0.0703 | 1 | 9 |
| GP6 Signaling Pathway | 1.45 | 0.0703 | -2.828 | 9 |
| Striated Muscle Contraction | 1.44 | 0.111 | -2 | 4 |
| Sperm Motility | 1.43 | 0.0584 | -1.342 | 15 |
| Pathogenesis of Multiple Sclerosis | 1.43 | 0.222 | - | 2 |
| G alpha (i) signalling events | 1.42 | 0.0581 | -0.258 | 15 |
| WNT/SHH Axonal Guidance Signaling Pathway | 1.4 | 0.0662 | -1.667 | 10 |
| Superpathway of Melatonin Degradation | 1.4 | 0.0769 | - | 7 |
| Regulation of the Epithelial Mesenchymal Transition by Growth Factors Pathway | 1.39 | 0.0619 | -2.333 | 12 |
| TR/RXR Activation | 1.39 | 0.0687 | -2.121 | 9 |
| Mitotic G1 phase and G1/S transition | 1.38 | 0.0682 | -1 | 9 |
| Ribonucleotide Reductase Signaling Pathway | 1.37 | 0.0632 | -1 | 11 |
| G-Protein Coupled Receptor Signaling | 1.37 | 0.0477 | -3.087 | 34 |
| Lymphotoxin β Receptor Signaling | 1.37 | 0.0909 | -0.447 | 5 |
| Role of Cytokines in Mediating Communication between Immune Cells | 1.37 | 0.0909 | - | 5 |
| Regulation of TP53 Expression and Degradation | 1.37 | 0.105 | -1 | 4 |
| Phase I - Functionalization of compounds | 1.35 | 0.0708 | -2.121 | 8 |
| RAF-independent MAPK1/3 activation | 1.35 | 0.13 | - | 3 |
| Apelin Cardiac Fibroblast Signaling Pathway | 1.35 | 0.13 | - | 3 |
| Urea cycle | 1.34 | 0.2 | - | 2 |
| RUNX1 and FOXP3 control the development of regulatory T lymphocytes (Tregs) | 1.34 | 0.2 | - | 2 |
| Prednisone ADME | 1.34 | 0.2 | - | 2 |
| Citrulline Biosynthesis | 1.34 | 0.2 | - | 2 |
| Lung Ionic Balance Signaling Pathway | 1.34 | 0.0498 | -3.922 | 26 |
| NGF-stimulated transcription | 1.33 | 0.103 | -2 | 4 |
| Nicotine Degradation III | 1.33 | 0.08 | - | 6 |
| Xenobiotic Metabolism PXR Signaling Pathway | 1.33 | 0.0588 | -0.816 | 13 |
| Class B/2 (Secretin family receptors) | 1.31 | 0.0737 | -0.378 | 7 |
| Human Embryonic Stem Cell Pluripotency | 1.31 | 0.06 | -2.887 | 12 |

Table S5: The top canonical pathways enriched in AH/HC by DEG analysis

| Ingenuity Canonical Pathways | -log(p-value) | Ratio | z-score | Gene count |
| --- | --- | --- | --- | --- |
| Heme signaling | 2.79 | 0.0851 | 1 | 4 |
| Circadian Clock | 2.45 | 0.069 | -2 | 4 |
| Triacylglycerol Biosynthesis | 2.45 | 0.069 | - | 4 |
| Interferon gamma signaling | 2.38 | 0.051 | -1.342 | 5 |
| cAMP-mediated signaling | 2.32 | 0.0332 | -1.134 | 8 |
| Stearate Biosynthesis I (Animals) | 2.32 | 0.0635 | - | 4 |
| Sumoylation Pathway | 2.32 | 0.0495 | - | 5 |
| Macrophage Alternative Activation Signaling Pathway | 2.31 | 0.0365 | -2.236 | 7 |
| Complement System | 2.07 | 0.0769 | - | 3 |
| Circadian Rhythm Signaling | 2.04 | 0.0296 | - | 8 |
| Cardiac β-adrenergic Signaling | 1.98 | 0.0353 | -1 | 6 |
| Vitamin E | 1.97 | 1 | - | 1 |
| Melatonin Degradation III | 1.97 | 1 | - | 1 |
| Adenine and Adenosine Salvage VI | 1.97 | 1 | - | 1 |
| Extrinsic Prothrombin Activation Pathway | 1.9 | 0.125 | - | 2 |
| PFKFB4 Signaling Pathway | 1.8 | 0.0612 | - | 3 |
| Complement cascade | 1.8 | 0.0368 | -0.447 | 5 |
| GABA synthesis, release, reuptake and degradation | 1.76 | 0.105 | - | 2 |
| Apelin Adipocyte Signaling Pathway | 1.73 | 0.0426 | - | 4 |
| UVB-Induced MAPK Signaling | 1.69 | 0.0556 | - | 3 |
| Nucleotide salvage | 1.6 | 0.087 | - | 2 |
| IL-10 Signaling | 1.58 | 0.0323 | - | 5 |
| GABA receptor activation | 1.57 | 0.05 | - | 3 |
| Relaxin Signaling | 1.57 | 0.0321 | - | 5 |
| Serotonin Receptor Signaling | 1.5 | 0.0213 | -2.53 | 10 |
| Xanthine and Xanthosine Salvage | 1.5 | 0.333 | - | 1 |
| 4-aminobutyrate Degradation I | 1.5 | 0.333 | - | 1 |
| Xenobiotic Metabolism PXR Signaling Pathway | 1.48 | 0.0271 | -1.342 | 6 |
| Phase I - Functionalization of compounds | 1.47 | 0.0354 | -1 | 4 |
| Ribavirin ADME | 1.45 | 0.0448 | - | 3 |
| PXR/RXR Activation | 1.45 | 0.0448 | - | 3 |
| CDP-diacylglycerol Biosynthesis I | 1.44 | 0.0714 | - | 2 |
| Sleep NREM Signaling Pathway | 1.43 | 0.0345 | -1 | 4 |
| Glycolysis I | 1.41 | 0.069 | - | 2 |
| Sphingosine-1-phosphate Signaling | 1.4 | 0.0336 | - | 4 |
| FOXO-mediated transcription of oxidative stress, metabolic and neuronal genes | 1.39 | 0.0667 | - | 2 |
| Phosphatidylglycerol Biosynthesis II (Non-plastidic) | 1.39 | 0.0667 | - | 2 |
| Gluconeogenesis I | 1.39 | 0.0667 | - | 2 |
| Threonine catabolism | 1.37 | 0.25 | - | 1 |
| Guanine and Guanosine Salvage I | 1.37 | 0.25 | - | 1 |
| Adenine and Adenosine Salvage I | 1.37 | 0.25 | - | 1 |
| Molybdenum Cofactor Biosynthesis | 1.37 | 0.25 | - | 1 |
| Ephrin B Signaling | 1.36 | 0.0411 | - | 3 |
| RAR Activation | 1.34 | 0.0208 | -2.333 | 9 |
| RHO GTPases activate IQGAPs | 1.33 | 0.0625 | - | 2 |
| MHC class II antigen presentation | 1.3 | 0.0312 | -2 | 4 |

Table S6: The top canonical pathways enriched in AH/HC by DSD analysis

| Ingenuity Canonical Pathways | -log(p-value) | Ratio | z-score | Gene count |
| --- | --- | --- | --- | --- |
| RHO GTPases Activate Formins | 7.54 | 0.136 | 4.243 | 19 |
| Cohesin Chromatin Regulation Pathway | 7.14 | 0.0989 | 3.051 | 26 |
| Pulmonary Fibrosis Idiopathic Signaling Pathway | 6.89 | 0.089 | 4.707 | 29 |
| Wound Healing Signaling Pathway | 6.44 | 0.0964 | 4.082 | 24 |
| Hepatic Fibrosis / Hepatic Stellate Cell Activation | 5.4 | 0.099 | - | 19 |
| Interleukin-4 and Interleukin-13 signaling | 5.34 | 0.126 | 2.138 | 14 |
| Neutrophil degranulation | 4.87 | 0.0671 | 5.657 | 32 |
| Collagen biosynthesis and modifying enzymes | 4.62 | 0.149 | 3.162 | 10 |
| Sensory processing of sound by inner hair cells of the cochlea | 4.51 | 0.145 | 3.162 | 10 |
| Collagen chain trimerization | 4.44 | 0.182 | 2.828 | 8 |
| Mitotic Prometaphase | 4.44 | 0.0878 | 4.123 | 18 |
| Agrin Interactions at Neuromuscular Junction | 3.8 | 0.132 | 2.646 | 9 |
| Germ Cell-Sertoli Cell Junction Signaling | 3.8 | 0.0877 | - | 15 |
| PAK Signaling | 3.74 | 0.102 | 3.162 | 12 |
| Reelin Signaling in Neurons | 3.67 | 0.0942 | 2.714 | 13 |
| ABRA Signaling Pathway | 3.6 | 0.114 | 2.53 | 10 |
| Molecular Mechanisms of Cancer | 3.57 | 0.0511 | 5.947 | 44 |
| L1CAM interactions | 3.54 | 0.0968 | 3.317 | 12 |
| Semaphorin Signaling in Neurons | 3.51 | 0.136 | - | 8 |
| Arachidonic acid metabolism | 3.51 | 0.156 | 2.646 | 7 |
| Axonal Guidance Signaling | 3.48 | 0.058 | - | 30 |
| Ion channel transport | 3.48 | 0.082 | 3.873 | 15 |
| Synaptogenesis Signaling Pathway | 3.35 | 0.0665 | 4.025 | 21 |
| Synthesis of Lipoxins (LX) | 3.33 | 0.5 | - | 3 |
| Collagen degradation | 3.27 | 0.125 | 2.828 | 8 |
| Semaphorin interactions | 3.27 | 0.125 | 2.828 | 8 |
| G alpha (12/13) signalling events | 3.27 | 0.113 | 3 | 9 |
| Interferon gamma signaling | 3.22 | 0.102 | 3.162 | 10 |
| Crosstalk between Dendritic Cells and Natural Killer Cells | 3.22 | 0.102 | 2 | 10 |
| Interferon Signaling | 3.18 | 0.162 | 2 | 6 |
| Gap junction trafficking and regulation | 3.17 | 0.137 | 2.449 | 7 |
| Multiple Sclerosis Signaling Pathway | 3.12 | 0.0731 | 3.873 | 16 |
| p75 NTR receptor-mediated signalling | 3.1 | 0.0823 | 3.606 | 13 |
| Epithelial Adherens Junction Signaling | 3.1 | 0.0823 | 0.577 | 13 |
| GPER1 signaling | 3.05 | 0.0971 | 3.162 | 10 |
| ID1 Signaling Pathway | 3.03 | 0.0743 | 2.138 | 15 |
| Ephrin Receptor Signaling | 3.03 | 0.0743 | 3.162 | 15 |
| Signaling by Rho Family GTPases | 3.02 | 0.0674 | 3.742 | 18 |
| Natural Killer Cell Signaling | 3.01 | 0.0739 | 1.732 | 15 |
| Leukotriene Biosynthesis | 3 | 0.25 | - | 4 |
| Sensory processing of sound by outer hair cells of the cochlea | 2.97 | 0.127 | 2.646 | 7 |
| Parkinson's Signaling Pathway | 2.96 | 0.0633 | 0.943 | 20 |
| Cell Cycle Checkpoints | 2.93 | 0.0662 | 4.243 | 18 |
| Cell junction organization | 2.83 | 0.0978 | 3 | 9 |
| Cardiac conduction | 2.81 | 0.0846 | 2.828 | 11 |
| Serotonin Receptor Signaling | 2.8 | 0.0553 | 4.082 | 26 |
| Phagosome Maturation | 2.79 | 0.076 | - | 13 |
| Mitotic Metaphase and Anaphase | 2.78 | 0.0678 | 3.873 | 16 |
| Assembly and cell surface presentation of NMDA receptors | 2.77 | 0.136 | 2.236 | 6 |
| Aggrephagy | 2.77 | 0.136 | 2.236 | 6 |
| Interferon alpha/beta signaling | 2.77 | 0.105 | 2.828 | 8 |
| Regulation of the Epithelial Mesenchymal Transition by Growth Factors Pathway | 2.75 | 0.0722 | 3.606 | 14 |
| Class B/2 (Secretin family receptors) | 2.73 | 0.0947 | 3 | 9 |
| Role of MAPK Signaling in Promoting the Pathogenesis of Influenza | 2.72 | 0.0877 | 3.162 | 10 |
| RAF/MAP kinase cascade | 2.71 | 0.0649 | 4 | 17 |
| Assembly of collagen fibrils and other multimeric structures | 2.71 | 0.115 | 2.646 | 7 |
| Autism Signaling Pathway | 2.69 | 0.0615 | 4.123 | 19 |
| ILK Signaling | 2.69 | 0.0711 | 2.309 | 14 |
| Caveolar-mediated Endocytosis Signaling | 2.66 | 0.101 | 2.236 | 8 |
| Kinesins | 2.66 | 0.113 | 2.449 | 7 |
| RHOGDI Signaling | 2.66 | 0.0682 | -2.887 | 15 |
| RHO GTPases activate IQGAPs | 2.66 | 0.156 | 2 | 5 |
| Nuclear Cytoskeleton Signaling Pathway | 2.64 | 0.0679 | 3.742 | 15 |
| Human Embryonic Stem Cell Pluripotency | 2.62 | 0.07 | 3.742 | 14 |
| CGAS-STING Signaling Pathway | 2.6 | 0.0797 | 3.317 | 11 |
| Tumor Microenvironment Pathway | 2.59 | 0.0722 | 2.309 | 13 |
| mRNA Editing | 2.59 | 0.3 | - | 3 |
| CREB Signaling in Neurons | 2.58 | 0.0503 | 4.707 | 31 |
| Role of Osteoclasts in Rheumatoid Arthritis Signaling Pathway | 2.57 | 0.0599 | 2.982 | 19 |
| Virus Entry via Endocytic Pathways | 2.55 | 0.0833 | 2.828 | 10 |
| Role of MAPK Signaling in the Pathogenesis of Influenza | 2.5 | 0.0952 | - | 8 |
| Kinetochore Metaphase Signaling Pathway | 2.49 | 0.0874 | 1.414 | 9 |
| Abacavir ADME | 2.47 | 0.104 | 2.646 | 7 |
| Mitotic Roles of Polo-Like Kinase | 2.47 | 0.104 | 1 | 7 |
| Prostanoid Biosynthesis | 2.46 | 0.273 | - | 3 |
| Neurovascular Coupling Signaling Pathway | 2.44 | 0.0647 | 3.357 | 15 |
| RHO GTPase cycle | 2.43 | 0.0533 | 4.899 | 24 |
| Myelination Signaling Pathway | 2.41 | 0.0579 | 3.441 | 19 |
| Hematoma Resolution Signaling Pathway | 2.38 | 0.0618 | -1.5 | 16 |
| PPAR Signaling | 2.35 | 0.0833 | -3 | 9 |
| Pathogen Induced Cytokine Storm Signaling Pathway | 2.34 | 0.055 | 4.472 | 21 |
| Glioblastoma Multiforme Signaling | 2.34 | 0.0702 | 3.162 | 12 |
| Glioma Invasiveness Signaling | 2.33 | 0.0986 | 1.89 | 7 |
| Remodeling of Epithelial Adherens Junctions | 2.33 | 0.0986 | - | 7 |
| PTEN Signaling | 2.3 | 0.0728 | -2.53 | 11 |
| Translocation of SLC2A4 (GLUT4) to the plasma membrane | 2.3 | 0.0972 | 2.449 | 7 |
| TR/RXR Activation | 2.28 | 0.0763 | 1.414 | 10 |
| Regulation of Actin-based Motility by Rho | 2.27 | 0.0811 | 2.646 | 9 |
| Transcriptional regulation of testis differentiation | 2.24 | 0.231 | - | 3 |
| DNA Damage/Telomere Stress Induced Senescence | 2.24 | 0.107 | 2.449 | 6 |
| O-linked glycosylation | 2.22 | 0.0796 | 3 | 9 |
| HSP90 chaperone cycle for steroid hormone receptors in the presence of ligand | 2.2 | 0.105 | 2.236 | 6 |
| Sertoli Cell-Sertoli Cell Junction Signaling | 2.19 | 0.0607 | 3.207 | 15 |
| Colorectal Cancer Metastasis Signaling | 2.19 | 0.059 | 3.742 | 16 |
| Oxytocin in Brain Signaling Pathway | 2.19 | 0.0647 | 2.496 | 13 |
| Macropinocytosis Signaling | 2.17 | 0.0921 | 2 | 7 |
| Signaling by PDGF | 2.17 | 0.103 | 2.449 | 6 |
| Cancer Drug Resistance by Drug Efflux | 2.17 | 0.103 | 2 | 6 |
| Breast Cancer Regulation by Stathmin1 | 2.16 | 0.0479 | 4.6 | 29 |
| RAC Signaling | 2.15 | 0.073 | 3 | 10 |
| Intra-Golgi and retrograde Golgi-to-ER traffic | 2.14 | 0.0637 | 3.464 | 13 |
| HMGB1 Signaling | 2.12 | 0.0688 | 3.162 | 11 |
| STAT3 Pathway | 2.11 | 0.0719 | 3 | 10 |
| GABAergic Receptor Signaling Pathway (Enhanced) | 2.08 | 0.0714 | 0.333 | 10 |
| Sphingosine-1-phosphate Signaling | 2.08 | 0.0756 | 1.667 | 9 |
| Signaling by MET | 2.08 | 0.0886 | 2.646 | 7 |
| Eicosanoid Signaling | 2.03 | 0.0567 | 4 | 16 |
| Interconversion of nucleotide di- and triphosphates | 2.02 | 0.138 | 2 | 4 |
| EGR2 and SOX10-mediated initiation of Schwann cell myelination | 2.02 | 0.138 | 2 | 4 |
| Renin-Angiotensin Signaling | 2.01 | 0.0738 | 2.449 | 9 |
| G alpha (s) signalling events | 1.98 | 0.069 | 3.162 | 10 |
| Osteoarthritis Pathway | 1.97 | 0.0588 | 1.897 | 14 |
| Potassium Channels | 1.97 | 0.0777 | 2.828 | 8 |
| Semaphorin Neuronal Repulsive Signaling Pathway | 1.94 | 0.068 | -1.265 | 10 |
| Role of Osteoblasts in Rheumatoid Arthritis Signaling Pathway | 1.94 | 0.0583 | 1.604 | 14 |
| Cellular Effects of Sildenafil (Viagra) | 1.94 | 0.0453 | -1.461 | 31 |
| Activation of anterior HOX genes in hindbrain during early embryogenesis | 1.94 | 0.0833 | 2.449 | 7 |
| Integrin cell surface interactions | 1.91 | 0.0824 | 2.646 | 7 |
| Calcium Signaling | 1.91 | 0.0596 | 3.162 | 13 |
| Paxillin Signaling | 1.9 | 0.0755 | 2.646 | 8 |
| MHC class II antigen presentation | 1.88 | 0.0703 | 2.828 | 9 |
| GP6 Signaling Pathway | 1.88 | 0.0703 | 3 | 9 |
| ERK/MAPK Signaling | 1.88 | 0.0591 | 1.897 | 13 |
| Protein Kinase A Signaling | 1.88 | 0.0505 | 0.775 | 20 |
| Extracellular matrix organization | 1.88 | 0.0748 | 2.828 | 8 |
| Ribavirin ADME | 1.87 | 0.0896 | 2.449 | 6 |
| PDGF Signaling | 1.86 | 0.0805 | 2.646 | 7 |
| Nucleosome assembly | 1.86 | 0.102 | 2.236 | 5 |
| Amyloid fiber formation | 1.83 | 0.0795 | 2.646 | 7 |
| S100 Family Signaling Pathway | 1.83 | 0.0435 | 4.382 | 34 |
| PI3K/AKT Signaling | 1.82 | 0.06 | 2.121 | 12 |
| Role of Pattern Recognition Receptors in Recognition of Bacteria and Viruses | 1.81 | 0.0649 | 2 | 10 |
| Role of JAK1 and JAK3 in γc Cytokine Signaling | 1.81 | 0.087 | 2.449 | 6 |
| NAP1L1 Transcription Regulation Signaling Pathway | 1.81 | 0.0787 | 2.449 | 7 |
| Regulation of Cellular Mechanics by Calpain Protease | 1.81 | 0.0787 | 1.342 | 7 |
| Pulmonary Healing Signaling Pathway | 1.81 | 0.0597 | 3.464 | 12 |
| 14-3-3-mediated Signaling | 1.8 | 0.0682 | 2.646 | 9 |
| Transcriptional regulation of granulopoiesis | 1.79 | 0.098 | 2.236 | 5 |
| Cell Cycle: G2/M DNA Damage Checkpoint Regulation | 1.79 | 0.098 | 0 | 5 |
| FOXO-mediated transcription | 1.77 | 0.158 | - | 3 |
| IL-27 Signaling Pathway | 1.77 | 0.0672 | 1.414 | 9 |
| Sheddase Signaling Pathway | 1.76 | 0.0588 | 1.155 | 12 |
| Ceramide Signaling | 1.76 | 0.0769 | 1.633 | 7 |
| Actin Nucleation by ARP-WASP Complex | 1.76 | 0.0769 | 2.449 | 7 |
| B-WICH complex positively regulates rRNA expression | 1.75 | 0.0962 | 2.236 | 5 |
| Neuroinflammation Signaling Pathway | 1.75 | 0.0515 | 1.604 | 17 |
| Xenobiotic Metabolism Signaling | 1.73 | 0.0514 | - | 17 |
| IL-9 Signaling | 1.73 | 0.114 | 2 | 4 |
| CCR3 Signaling in Eosinophils | 1.73 | 0.0662 | 2.449 | 9 |
| PD-1, PD-L1 cancer immunotherapy pathway | 1.72 | 0.0702 | 0.378 | 8 |
| G-Protein Coupled Receptor Signaling | 1.72 | 0.0435 | 4.382 | 31 |
| EPH-Ephrin signaling | 1.71 | 0.0753 | 2.646 | 7 |
| Role of Tissue Factor in Cancer | 1.7 | 0.0577 | 2.714 | 12 |
| Role of PKR in Interferon Induction and Antiviral Response | 1.69 | 0.0652 | 1.89 | 9 |
| Striated Muscle Contraction | 1.69 | 0.111 | 2 | 4 |
| Sleep NREM Signaling Pathway | 1.68 | 0.069 | -0.378 | 8 |
| Integrin Signaling | 1.67 | 0.0571 | 3.464 | 12 |
| CLEAR Signaling Pathway | 1.67 | 0.0526 | -2.84 | 15 |
| Immunogenic Cell Death Signaling Pathway | 1.67 | 0.0737 | 2.449 | 7 |
| Sertoli Cell-Germ Cell Junction Signaling Pathway (Enhanced) | 1.65 | 0.0551 | -0.277 | 13 |
| Formation of the ureteric bud | 1.65 | 0.143 | - | 3 |
| HOTAIR Regulatory Pathway | 1.63 | 0.0606 | 3 | 10 |
| Irritable Bowel Syndrome Signaling Pathway | 1.62 | 0.0519 | 2.84 | 15 |
| TGF-β Signaling | 1.62 | 0.0722 | 1.633 | 7 |
| Cachexia Signaling Pathway | 1.62 | 0.0489 | 3.153 | 18 |
| Inhibition of Matrix Metalloproteases | 1.61 | 0.105 | - | 4 |
| Immunoregulatory interactions between a Lymphoid and a non-Lymphoid cell | 1.6 | 0.0558 | 3.464 | 12 |
| Neurexins and neuroligins | 1.6 | 0.0877 | 2 | 5 |
| Hedgehog 'on' state | 1.59 | 0.0625 | 3 | 9 |
| Protein folding | 1.58 | 0.0707 | 2.449 | 7 |
| Actin Cytoskeleton Signaling | 1.58 | 0.0537 | 2.53 | 13 |
| Orexin Signaling Pathway | 1.58 | 0.0537 | 3.051 | 13 |
| NGF-stimulated transcription | 1.58 | 0.103 | 2 | 4 |
| VDR/RXR Activation | 1.57 | 0.0769 | - | 6 |
| PKR-mediated signaling | 1.57 | 0.0769 | 2.236 | 6 |
| RHO GTPases Activate Rhotekin and Rhophilins | 1.57 | 0.222 | - | 2 |
| RUNX1 regulates megakaryocyte differentiation and platelet function | 1.57 | 0.0862 | -0.447 | 5 |
| NOD1/2 Signaling Pathway | 1.57 | 0.057 | 3.162 | 11 |
| Cardiac Hypertrophy Signaling (Enhanced) | 1.56 | 0.0447 | 4.359 | 24 |
| Phagosome Formation | 1.56 | 0.0425 | 5.014 | 30 |
| Regulation of the Epithelial-Mesenchymal Transition Pathway | 1.55 | 0.0567 | - | 11 |
| AMPK Signaling | 1.55 | 0.0533 | 0.816 | 13 |
| NRF2-mediated Oxidative Stress Response | 1.55 | 0.0519 | 2.828 | 14 |
| Tuberculosis Active Signaling Pathway | 1.54 | 0.0531 | -1.732 | 13 |
| FXR/RXR Activation | 1.54 | 0.0564 | -0.302 | 11 |
| Gαi Signaling | 1.54 | 0.0612 | 1.134 | 9 |
| Asparagine Biosynthesis I | 1.53 | 1 | - | 1 |
| COPI-mediated anterograde transport | 1.52 | 0.0686 | 2.449 | 7 |
| Role of JAK family kinases in IL-6-type Cytokine Signaling | 1.5 | 0.0741 | 2.449 | 6 |
| Thrombin Signaling | 1.5 | 0.0538 | 2.121 | 12 |
| Adrenergic Receptor Signaling Pathway (Enhanced) | 1.5 | 0.0556 | 0 | 11 |
| Other interleukin signaling | 1.5 | 0.125 | - | 3 |
| Interleukin-6 family signaling | 1.5 | 0.125 | - | 3 |
| Agranulocyte Adhesion and Diapedesis | 1.48 | 0.0536 | - | 12 |
| Interleukin-9 signaling | 1.48 | 0.2 | - | 2 |
| RHOBTB3 ATPase cycle | 1.48 | 0.2 | - | 2 |
| Gαs Signaling | 1.48 | 0.063 | 2.646 | 8 |
| Netrin Signaling | 1.47 | 0.0571 | 0.632 | 10 |
| Mitotic G2-G2/M phases | 1.47 | 0.055 | 3.162 | 11 |
| Oncostatin M Signaling | 1.44 | 0.093 | 2 | 4 |
| Antigen Presentation Pathway | 1.44 | 0.093 | - | 4 |
| Granulocyte Adhesion and Diapedesis | 1.42 | 0.0539 | - | 11 |
| Activation of NMDA receptors and postsynaptic events | 1.42 | 0.0706 | 2 | 6 |
| Retinoid metabolism and transport | 1.41 | 0.0909 | 2 | 4 |
| DNA methylation | 1.41 | 0.115 | - | 3 |
| Integration of energy metabolism | 1.4 | 0.0648 | 2.646 | 7 |
| Phenylalanine and tyrosine metabolism | 1.4 | 0.182 | - | 2 |
| GABA Receptor Signaling | 1.38 | 0.0602 | - | 8 |
| Carboxyterminal post-translational modifications of tubulin | 1.38 | 0.0889 | - | 4 |
| Syndecan interactions | 1.36 | 0.111 | - | 3 |
| Microautophagy Signaling Pathway | 1.35 | 0.0566 | - | 9 |
| DNA damage-induced 14-3-3σ Signaling | 1.35 | 0.087 | -1 | 4 |
| ERBB4 Signaling | 1.34 | 0.0746 | 2 | 5 |
| Telomere Maintenance | 1.33 | 0.0674 | 2.449 | 6 |
| Atherosclerosis Signaling | 1.33 | 0.0588 | 2.828 | 8 |
| RHO GTPases activate PKNs | 1.32 | 0.107 | - | 3 |
| Activated PKN1 stimulates transcription of AR regulated genes KLK2 and KLK3 | 1.32 | 0.107 | - | 3 |
| Preeclampsia Signaling Pathway | 1.32 | 0.0559 | -0.333 | 9 |
| Interleukin-2 family signaling | 1.32 | 0.0851 | 2 | 4 |
| Glutamate Receptor Signaling | 1.31 | 0.0735 | - | 5 |
| Hepatitis B Chronic Liver Pathogenesis Signaling Pathway | 1.3 | 0.0532 | 1.414 | 10 |

Table S7: The top common pathways enriched in AH/HC by DEG and DSD analysis

| Ingenuity Canonical Pathways | -log(p-value) | Ratio | z-score | Gene count |
| --- | --- | --- | --- | --- |
| Extracellular matrix organization | 17.2 | 0.467 | 4.243 | 50 |
| Molecular Mechanisms of Cancer | 15.8 | 0.226 | 2.691 | 195 |
| Sheddase Signaling Pathway | 14.9 | 0.343 | 3.586 | 70 |
| LXR/RXR Activation | 13.7 | 0.392 | -4.422 | 51 |
| DHCR24 Signaling Pathway | 13.6 | 0.378 | -3.781 | 54 |
| S100 Family Signaling Pathway | 13.1 | 0.221 | 3.086 | 173 |
| Regulation of Insulin-like Growth Factor (IGF) transport and uptake by IGFBPs | 12.7 | 0.387 | 0.289 | 48 |
| Tumor Microenvironment Pathway | 12.3 | 0.333 | 4.106 | 60 |
| Atherosclerosis Signaling | 12.2 | 0.368 | 0 | 50 |
| Acute Phase Response Signaling | 12.1 | 0.325 | 0.949 | 62 |
| Phase I - Functionalization of compounds | 11.8 | 0.389 | -4.529 | 44 |
| Integrin cell surface interactions | 11.7 | 0.435 | 4.11 | 37 |
| Wound Healing Signaling Pathway | 11.1 | 0.289 | 2.151 | 72 |
| Post-translational protein phosphorylation | 10.8 | 0.383 | 0.469 | 41 |
| Coagulation System | 10.3 | 0.6 | -1.606 | 21 |
| Interleukin-4 and Interleukin-13 signaling | 10.2 | 0.369 | 1.093 | 41 |
| Role of Osteoclasts in Rheumatoid Arthritis Signaling Pathway | 9.85 | 0.259 | 4.129 | 82 |
| Hepatic Cholestasis | 9.65 | 0.284 | 2.84 | 64 |
| Hematoma Resolution Signaling Pathway | 9.4 | 0.27 | -2.183 | 70 |
| Axonal Guidance Signaling | 9.36 | 0.224 | - | 116 |
| Breast Cancer Regulation by Stathmin1 | 9.36 | 0.216 | 1.782 | 131 |
| LPS/IL-1 Mediated Inhibition of RXR Function | 9.27 | 0.258 | 1.091 | 77 |
| Pulmonary Fibrosis Idiopathic Signaling Pathway | 9.21 | 0.252 | 5.143 | 82 |
| Response to elevated platelet cytosolic Ca2+ | 9.18 | 0.333 | -0.302 | 44 |
| Phagosome Formation | 8.94 | 0.207 | 1.623 | 146 |
| Granulocyte Adhesion and Diapedesis | 8.8 | 0.284 | - | 58 |
| Aspirin ADME | 8.65 | 0.5 | -3.411 | 22 |
| Hepatic Fibrosis / Hepatic Stellate Cell Activation | 8.51 | 0.286 | - | 55 |
| IL-6 Signaling | 8.45 | 0.326 | 1.947 | 42 |
| RAR Activation | 8.22 | 0.226 | -0.308 | 98 |
| Role of IL-17A in Arthritis | 8.05 | 0.431 | - | 25 |
| Sertoli Cell-Sertoli Cell Junction Signaling | 7.88 | 0.259 | 2.898 | 64 |
| G-Protein Coupled Receptor Signaling | 7.87 | 0.201 | 0.256 | 143 |
| FXR/RXR Activation | 7.81 | 0.277 | -0.146 | 54 |
| Collagen degradation | 7.69 | 0.406 | 3.53 | 26 |
| HEY1 Signaling Pathway | 7.66 | 0.292 | 2.771 | 47 |
| Irritable Bowel Syndrome Signaling Pathway | 7.62 | 0.246 | 2.492 | 71 |
| Docosahexaenoic Acid (DHA) Signaling | 7.59 | 0.255 | 0.128 | 64 |
| Cellular Effects of Sildenafil (Viagra) | 7.49 | 0.2 | -0.26 | 137 |
| Xenobiotic Metabolism Signaling | 7.46 | 0.236 | - | 78 |
| Lung Ionic Balance Signaling Pathway | 7.34 | 0.211 | 0.191 | 110 |
| Formation of Fibrin Clot (Clotting Cascade) | 7.32 | 0.487 | -2.065 | 19 |
| CREB Signaling in Neurons | 7.26 | 0.203 | 0.093 | 125 |
| Hepatic Fibrosis Signaling Pathway | 7.25 | 0.221 | 2.491 | 92 |
| IL-8 Signaling | 7.19 | 0.264 | 3.429 | 55 |
| Bupropion Degradation | 7.17 | 0.577 | - | 15 |
| Sertoli Cell-Germ Cell Junction Signaling Pathway (Enhanced) | 7.11 | 0.254 | 0 | 60 |
| HIF1α Signaling | 7.04 | 0.262 | 1.236 | 55 |
| Glycosaminoglycan metabolism | 7.02 | 0.305 | 3.244 | 39 |
| Cachexia Signaling Pathway | 7 | 0.226 | 2.111 | 83 |
| Serotonin Receptor Signaling | 6.95 | 0.213 | 1.91 | 100 |
| Activin Inhibin Signaling Pathway | 6.89 | 0.259 | 2.292 | 55 |
| Senescence Pathway | 6.87 | 0.236 | 2.047 | 71 |
| ID1 Signaling Pathway | 6.84 | 0.262 | 0.832 | 53 |
| Assembly of collagen fibrils and other multimeric structures | 6.84 | 0.393 | 4.082 | 24 |
| Agranulocyte Adhesion and Diapedesis | 6.81 | 0.254 | - | 57 |
| Dissolution of Fibrin Clot | 6.67 | 0.769 | 1.265 | 10 |
| Nicotine Degradation III | 6.67 | 0.36 | -0.333 | 27 |
| p53 Signaling | 6.64 | 0.327 | 0.655 | 32 |
| Leukocyte Extravasation Signaling | 6.63 | 0.263 | 2.592 | 51 |
| IL-17A Signaling in Fibroblasts | 6.59 | 0.35 | 2.887 | 28 |
| Nicotine Degradation II | 6.53 | 0.341 | -0.632 | 29 |
| Airway Pathology in Chronic Obstructive Pulmonary Disease | 6.46 | 0.307 | -0.378 | 35 |
| Estrogen Biosynthesis | 6.46 | 0.426 | 0 | 20 |
| IL-17A Signaling in Airway Cells | 6.42 | 0.368 | 1.147 | 25 |
| Class A/1 (Rhodopsin-like receptors) | 6.42 | 0.226 | 0.577 | 75 |
| HMGB1 Signaling | 6.4 | 0.275 | 2.611 | 44 |
| Production of Nitric Oxide and Reactive Oxygen Species in Macrophages | 6.38 | 0.26 | -0.603 | 50 |
| Cardiac conduction | 6.34 | 0.292 | -1.3 | 38 |
| Clathrin-mediated Endocytosis Signaling | 6.33 | 0.254 | - | 53 |
| WNT/SHH Axonal Guidance Signaling Pathway | 6.29 | 0.278 | 1.718 | 42 |
| Colorectal Cancer Metastasis Signaling | 6.28 | 0.236 | 2.994 | 64 |
| Retinoid metabolism and transport | 6.28 | 0.432 | 0.229 | 19 |
| Aryl Hydrocarbon Receptor Signaling | 6.25 | 0.262 | 0.209 | 48 |
| IL-33 Signaling Pathway | 6.16 | 0.256 | 3.202 | 50 |
| Bile acid and bile salt metabolism | 6.1 | 0.422 | -2.524 | 19 |
| Transport of bile salts and organic acids, metal ions and amine compounds | 6.1 | 0.333 | -0.378 | 28 |
| Glycation Signaling Pathway | 5.99 | 0.244 | 3.641 | 55 |
| Bladder Cancer Signaling | 5.97 | 0.294 | 1.897 | 35 |
| PXR/RXR Activation | 5.96 | 0.358 | 0.832 | 24 |
| CD40 Signaling | 5.96 | 0.358 | 1.706 | 24 |
| Preeclampsia Signaling Pathway | 5.9 | 0.267 | 1.677 | 43 |
| Regulation of the Epithelial Mesenchymal Transition by Growth Factors Pathway | 5.85 | 0.253 | 2.846 | 49 |
| Semaphorin Neuronal Repulsive Signaling Pathway | 5.76 | 0.272 | 0.174 | 40 |
| Pathogen Induced Cytokine Storm Signaling Pathway | 5.7 | 0.212 | 2.692 | 81 |
| CDX Gastrointestinal Cancer Signaling Pathway | 5.64 | 0.249 | -1.896 | 49 |
| Elastic fibre formation | 5.57 | 0.409 | 3.3 | 18 |
| Germ Cell-Sertoli Cell Junction Signaling | 5.55 | 0.257 | - | 44 |
| Glutaminergic Receptor Signaling Pathway (Enhanced) | 5.51 | 0.218 | 0.361 | 71 |
| Melatonin Degradation I | 5.45 | 0.318 | -0.333 | 27 |
| Activation of Matrix Metalloproteinases | 5.42 | 0.455 | 0.775 | 15 |
| Arachidonic acid metabolism | 5.4 | 0.4 | -1.213 | 18 |
| Acetone Degradation I (to Methylglyoxal) | 5.39 | 0.415 | 1 | 17 |
| Collagen biosynthesis and modifying enzymes | 5.37 | 0.343 | 2.294 | 23 |
| Degradation of the extracellular matrix | 5.36 | 0.321 | 3.922 | 26 |
| NAFLD Signaling Pathway | 5.35 | 0.237 | 1.786 | 53 |
| Superpathway of Melatonin Degradation | 5.32 | 0.308 | -0.632 | 28 |
| Paxillin Signaling | 5.31 | 0.292 | 1.964 | 31 |
| IL-12 Signaling and Production in Macrophages | 5.3 | 0.231 | 2.496 | 56 |
| Role of Tissue Factor in Cancer | 5.29 | 0.24 | 2.263 | 50 |
| Inhibition of Matrix Metalloproteases | 5.21 | 0.421 | -2.138 | 16 |
| HOTAIR Regulatory Pathway | 5.2 | 0.255 | 3.363 | 42 |
| GP6 Signaling Pathway | 5.17 | 0.273 | 2.535 | 35 |
| Pancreatic Adenocarcinoma Signaling | 4.99 | 0.272 | 1.789 | 34 |
| Actin Cytoskeleton Signaling | 4.97 | 0.227 | 2.335 | 55 |
| Type II Diabetes Mellitus Signaling | 4.97 | 0.257 | 1.46 | 39 |
| Collagen chain trimerization | 4.9 | 0.386 | 1.698 | 17 |
| Autism Signaling Pathway | 4.88 | 0.214 | 1.664 | 66 |
| Osteoarthritis Pathway | 4.87 | 0.227 | 1.859 | 54 |
| Plasma lipoprotein assembly, remodeling, and clearance | 4.87 | 0.316 | -1.225 | 24 |
| Role of JAK family kinases in IL-6-type Cytokine Signaling | 4.85 | 0.309 | 1.043 | 25 |
| Xenobiotic Metabolism CAR Signaling Pathway | 4.83 | 0.227 | -0.186 | 53 |
| IL-10 Signaling | 4.75 | 0.252 | 0.973 | 39 |
| G alpha (i) signalling events | 4.75 | 0.221 | -0.132 | 57 |
| NRF2-mediated Oxidative Stress Response | 4.74 | 0.219 | 1.091 | 59 |
| Eicosanoid Signaling | 4.73 | 0.216 | 2.474 | 61 |
| BBSome Signaling Pathway | 4.72 | 0.193 | -0.314 | 95 |
| GP1b-IX-V activation signalling | 4.69 | 0.667 | 1.414 | 8 |
| L1CAM interactions | 4.65 | 0.266 | 1.219 | 33 |
| Neutrophil degranulation | 4.61 | 0.193 | 3.753 | 92 |
| ERK/MAPK Signaling | 4.58 | 0.227 | 2.197 | 50 |
| Myelination Signaling Pathway | 4.56 | 0.207 | 2.81 | 68 |
| Signaling by MET | 4.55 | 0.304 | 2.449 | 24 |
| Role of MAPK Signaling in Inhibiting the Pathogenesis of Influenza | 4.55 | 0.304 | 0.894 | 24 |
| Regulation of Cellular Mechanics by Calpain Protease | 4.55 | 0.292 | 1.508 | 26 |
| Cardiac Hypertrophy Signaling (Enhanced) | 4.53 | 0.188 | 2.429 | 101 |
| Human Embryonic Stem Cell Pluripotency | 4.41 | 0.23 | 0.295 | 46 |
| PI3K/AKT Signaling | 4.41 | 0.23 | 1.706 | 46 |
| IL-17 Signaling | 4.38 | 0.233 | 3.395 | 44 |
| Role of Chondrocytes in Rheumatoid Arthritis Signaling Pathway | 4.38 | 0.25 | 2.535 | 36 |
| Glioma Invasiveness Signaling | 4.37 | 0.31 | 1.528 | 22 |
| Cerebral Malformation Signaling Pathway | 4.37 | 0.265 | 0.539 | 31 |
| Noradrenaline and Adrenaline Degradation | 4.36 | 0.385 | -2.53 | 15 |
| Syndecan interactions | 4.33 | 0.444 | 2.887 | 12 |
| IL-13 Signaling Pathway | 4.32 | 0.26 | 1.095 | 32 |
| Pancreatic Secretion Signaling Pathway | 4.3 | 0.218 | 1.664 | 53 |
| Signaling by Rho Family GTPases | 4.3 | 0.213 | 2.53 | 57 |
| Cell surface interactions at the vascular wall | 4.27 | 0.224 | 3.464 | 48 |
| Cell junction organization | 4.27 | 0.283 | 2.746 | 26 |
| Ephrin A Signaling | 4.27 | 0.25 | 1.372 | 35 |
| ILK Signaling | 4.25 | 0.228 | 0.649 | 45 |
| Triacylglycerol Biosynthesis | 4.22 | 0.328 | -0.258 | 19 |
| Xenobiotic Metabolism PXR Signaling Pathway | 4.22 | 0.222 | -1.512 | 49 |
| Neuroinflammation Signaling Pathway | 4.21 | 0.203 | 3 | 67 |
| Integrin Signaling | 4.18 | 0.224 | 2.03 | 47 |
| Erythropoietin Signaling Pathway | 4.17 | 0.232 | -0.16 | 42 |
| Ethanol Degradation II | 4.17 | 0.389 | -2.333 | 14 |
| Estrogen-Dependent Breast Cancer Signaling | 4.17 | 0.289 | 2 | 24 |
| Phase II - Conjugation of compounds | 4.16 | 0.266 | -3.157 | 29 |
| Regulation of the Epithelial-Mesenchymal Transition Pathway | 4.09 | 0.227 | - | 44 |
| MSP-RON Signaling in Cancer Cells Pathway | 4.07 | 0.245 | 2.611 | 35 |
| Ion channel transport | 4.06 | 0.23 | 0.617 | 42 |
| CGAS-STING Signaling Pathway | 4.03 | 0.246 | 4.352 | 34 |
| Superoxide Radicals Degradation | 4.03 | 0.75 | 0.816 | 6 |
| Parkinson's Signaling Pathway | 4.02 | 0.203 | -0.128 | 64 |
| Serotonin Degradation | 4.02 | 0.274 | -2.324 | 26 |
| Gap Junction Signaling | 4.01 | 0.2 | 1 | 67 |
| GADD45 Signaling | 4 | 0.317 | -0.229 | 19 |
| Role of IL-17F in Allergic Inflammatory Airway Diseases | 4 | 0.348 | 2.333 | 16 |
| RAF/MAP kinase cascade | 3.97 | 0.21 | 2.1 | 55 |
| Mitochondrial RNA degradation | 3.96 | 0.44 | 3.317 | 11 |
| Metallothioneins bind metals | 3.96 | 0.636 | -2.646 | 7 |
| Paracetamol ADME | 3.96 | 0.414 | 0 | 12 |
| FGF Signaling | 3.9 | 0.279 | 1.877 | 24 |
| PAK Signaling | 3.89 | 0.254 | 1.414 | 30 |
| Adrenergic Receptor Signaling Pathway (Enhanced) | 3.88 | 0.222 | -2.412 | 44 |
| Role of Osteoblasts in Rheumatoid Arthritis Signaling Pathway | 3.86 | 0.212 | 1.82 | 51 |
| O-linked glycosylation | 3.86 | 0.257 | 2.414 | 29 |
| 4-1BB Signaling in T Lymphocytes | 3.83 | 0.382 | 1.897 | 13 |
| Intrinsic Prothrombin Activation Pathway | 3.8 | 0.349 | -0.832 | 15 |
| SPINK1 General Cancer Pathway | 3.78 | 0.299 | 2.524 | 20 |
| IL-1 Signaling | 3.78 | 0.265 | 1.5 | 26 |
| TR/RXR Activation | 3.76 | 0.244 | -0.186 | 32 |
| HGF Signaling | 3.76 | 0.244 | 1.414 | 32 |
| Glioblastoma Multiforme Signaling | 3.75 | 0.228 | 1.616 | 39 |
| ERBB Signaling | 3.75 | 0.269 | 2.449 | 25 |
| Pulmonary Healing Signaling Pathway | 3.72 | 0.219 | 2.714 | 44 |
| Maturity Onset Diabetes of Young (MODY) Signaling | 3.7 | 0.282 | - | 22 |
| Neurotrophin/TRK Signaling | 3.7 | 0.282 | 0.243 | 22 |
| Reelin Signaling in Neurons | 3.67 | 0.239 | 1.257 | 33 |
| RHOGDI Signaling | 3.67 | 0.214 | -2.043 | 47 |
| eNOS Signaling | 3.66 | 0.23 | 0.2 | 37 |
| IGF-1 Signaling | 3.65 | 0.257 | 0 | 27 |
| Prostate Cancer Signaling | 3.65 | 0.25 | 1.347 | 29 |
| Atorvastatin ADME | 3.61 | 0.667 | -2.449 | 6 |
| Ribonucleotide Reductase Signaling Pathway | 3.59 | 0.224 | 3 | 39 |
| CDK5 Signaling | 3.58 | 0.255 | 0 | 27 |
| PPARα/RXRα Activation | 3.57 | 0.217 | -0.507 | 43 |
| Gα12/13 Signaling | 3.57 | 0.239 | 2.117 | 32 |
| Activation of NMDA receptors and postsynaptic events | 3.55 | 0.271 | 0.853 | 23 |
| LPS-stimulated MAPK Signaling | 3.55 | 0.271 | 1.789 | 23 |
| Gαq Signaling | 3.54 | 0.225 | 1.3 | 38 |
| Lymphotoxin β Receptor Signaling | 3.5 | 0.309 | 2.138 | 17 |
| Apelin Pancreas Signaling Pathway | 3.43 | 0.326 | 0.535 | 15 |
| IL-23 Signaling Pathway | 3.43 | 0.326 | 0.535 | 15 |
| GDNF Family Ligand-Receptor Interactions | 3.42 | 0.276 | 0.243 | 21 |
| RAC Signaling | 3.39 | 0.234 | 2.4 | 32 |
| Platelet Aggregation (Plug Formation) | 3.35 | 0.538 | 0.378 | 7 |
| Toll-like Receptor Signaling | 3.34 | 0.273 | 2 | 21 |
| Synaptogenesis Signaling Pathway | 3.28 | 0.193 | 2.364 | 61 |
| STAT3 Pathway | 3.27 | 0.23 | 0.218 | 32 |
| NF-κB Activation by Viruses | 3.26 | 0.269 | 2.524 | 21 |
| Fructose metabolism | 3.26 | 0.714 | -2.236 | 5 |
| Neurotransmitter clearance | 3.26 | 0.6 | -1.633 | 6 |
| Creatine metabolism | 3.26 | 0.6 | 0 | 6 |
| Prednisone ADME | 3.26 | 0.6 | -2.449 | 6 |
| Role of Osteoblasts, Osteoclasts and Chondrocytes in Rheumatoid Arthritis | 3.26 | 0.205 | - | 47 |
| Thrombin Signaling | 3.25 | 0.206 | 1.667 | 46 |
| PTEN Signaling | 3.25 | 0.225 | -0.447 | 34 |
| NAP1L1 Transcription Regulation Signaling Pathway | 3.24 | 0.258 | 2.711 | 23 |
| Role of Macrophages, Fibroblasts and Endothelial Cells in Rheumatoid Arthritis | 3.22 | 0.19 | 2.433 | 64 |
| NCAM signaling for neurite out-growth | 3.21 | 0.286 | 1.886 | 18 |
| Sleep REM Signaling Pathway | 3.2 | 0.245 | -1.961 | 26 |
| Signaling by PDGF | 3.2 | 0.293 | 2.668 | 17 |
| Hepatitis B Chronic Liver Pathogenesis Signaling Pathway | 3.18 | 0.213 | 2.53 | 40 |
| Neurovascular Coupling Signaling Pathway | 3.13 | 0.203 | 0.457 | 47 |
| Non-Small Cell Lung Cancer Signaling | 3.13 | 0.25 | 1.807 | 24 |
| Tryptophan catabolism | 3.1 | 0.5 | -1.89 | 7 |
| Role of IL-17A in Psoriasis | 3.1 | 0.5 | 1.342 | 7 |
| Ephrin Receptor Signaling | 3.09 | 0.208 | 1.706 | 42 |
| IL-17A Signaling in Gastric Cells | 3.09 | 0.385 | 1.134 | 10 |
| Nucleotide catabolism | 3.07 | 0.343 | -1.732 | 12 |
| IL-9 Signaling | 3.07 | 0.343 | 0.632 | 12 |
| Transport of inorganic cations/anions and amino acids/oligopeptides | 3.07 | 0.241 | 1.177 | 26 |
| Neurotransmitter release cycle | 3.03 | 0.325 | -1.387 | 13 |
| cAMP-mediated signaling | 3.01 | 0.199 | -1.508 | 48 |
| Interleukin-10 signaling | 3.01 | 0.311 | 2.673 | 14 |
| Hereditary Breast Cancer Signaling | 3 | 0.225 | 0.898 | 31 |
| Macrophage Alternative Activation Signaling Pathway | 2.99 | 0.208 | 0.48 | 40 |
| Bone Mineralization Signaling Pathway | 2.99 | 0.222 | 2.828 | 32 |
| Small Cell Lung Cancer Signaling | 2.99 | 0.245 | 1.941 | 24 |
| CXCR4 Signaling | 2.98 | 0.214 | 0.928 | 36 |
| Chondroitin Sulfate Biosynthesis | 2.97 | 0.268 | 1.941 | 19 |
| RANK Signaling in Osteoclasts | 2.95 | 0.247 | 2.065 | 23 |
| NOD1/2 Signaling Pathway | 2.95 | 0.207 | 3.124 | 40 |
| PIP3 activates AKT signaling | 2.94 | 0.221 | 0.707 | 32 |
| Renin-Angiotensin Signaling | 2.91 | 0.23 | 1.091 | 28 |
| Apelin Cardiac Fibroblast Signaling Pathway | 2.89 | 0.391 | -1.667 | 9 |
| Sucrose Degradation V (Mammalian) | 2.88 | 0.625 | -1.342 | 5 |
| Glyoxylate metabolism and glycine degradation | 2.86 | 0.421 | -2.828 | 8 |
| GABA synthesis, release, reuptake and degradation | 2.86 | 0.421 | 0 | 8 |
| Thyroid Cancer Signaling | 2.85 | 0.256 | 0.894 | 20 |
| Aldosterone Signaling in Epithelial Cells | 2.84 | 0.211 | 0.775 | 36 |
| Ovarian Cancer Signaling | 2.83 | 0.214 | 1.5 | 34 |
| April Mediated Signaling | 2.81 | 0.31 | 1.732 | 13 |
| PEDF Signaling | 2.81 | 0.25 | 1.213 | 21 |
| p38 MAPK Signaling | 2.8 | 0.226 | 2.041 | 28 |
| Glucocorticoid Receptor Signaling | 2.75 | 0.168 | - | 101 |
| Interleukin-6 family signaling | 2.74 | 0.375 | 1 | 9 |
| Androgen Biosynthesis | 2.74 | 0.375 | -2.236 | 9 |
| Cardiac Hypertrophy Signaling | 2.73 | 0.192 | 1.067 | 50 |
| Triglyceride metabolism | 2.72 | 0.316 | -0.577 | 12 |
| Actin Nucleation by ARP-WASP Complex | 2.71 | 0.242 | 1.732 | 22 |
| Lysine catabolism | 2.71 | 0.5 | -2.449 | 6 |
| B Cell Activating Factor Signaling | 2.71 | 0.302 | 1.897 | 13 |
| Uracil Degradation II (Reductive) | 2.69 | 1 | - | 3 |
| Thymine Degradation | 2.69 | 1 | - | 3 |
| Potassium Channels | 2.67 | 0.233 | 0 | 24 |
| Extrinsic Prothrombin Activation Pathway | 2.67 | 0.438 | - | 7 |
| Glycerophospholipid biosynthesis | 2.59 | 0.219 | 0 | 28 |
| Glioma Signaling | 2.59 | 0.219 | 0.5 | 28 |
| Endocannabinoid Developing Neuron Pathway | 2.59 | 0.219 | 0.816 | 28 |
| HER-2 Signaling in Breast Cancer | 2.58 | 0.194 | 3.904 | 44 |
| Regulation of Actin-based Motility by Rho | 2.55 | 0.225 | 1.069 | 25 |
| Amyotrophic Lateral Sclerosis Signaling | 2.54 | 0.222 | 1.279 | 26 |
| Role of NANOG in Mammalian Embryonic Stem Cell Pluripotency | 2.54 | 0.22 | 0.258 | 27 |
| Choline catabolism | 2.5 | 0.667 | -2 | 4 |
| Lipid particle organization | 2.5 | 0.667 | -1 | 4 |
| JAK/STAT Signaling | 2.5 | 0.241 | 1.414 | 20 |
| Metabolism of Angiotensinogen to Angiotensins | 2.49 | 0.412 | -1.89 | 7 |
| Neuregulin Signaling | 2.49 | 0.22 | 2.138 | 26 |
| EGF Signaling | 2.46 | 0.268 | 1.291 | 15 |
| Estrogen-mediated S-phase Entry | 2.46 | 0.346 | 1.667 | 9 |
| VDR/RXR Activation | 2.46 | 0.244 | 0.277 | 19 |
| Transcriptional regulation by RUNX2 | 2.44 | 0.275 | 1.941 | 14 |
| FAT10 Cancer Signaling Pathway | 2.44 | 0.275 | 1.732 | 14 |
| AMPK Signaling | 2.42 | 0.189 | 0.557 | 46 |
| Signaling by ALK | 2.42 | 0.323 | 1.897 | 10 |
| Sperm Motility | 2.42 | 0.187 | 0.258 | 48 |
| Tuberculosis Latent Signaling Pathway | 2.42 | 0.233 | -0.655 | 21 |
| Dermatan Sulfate Biosynthesis | 2.42 | 0.247 | 1.732 | 18 |
| Kinesins | 2.42 | 0.258 | 2.5 | 16 |
| IL-7 Signaling Pathway | 2.39 | 0.241 | 0.5 | 19 |
| Role of PKR in Interferon Induction and Antiviral Response | 2.39 | 0.21 | 1.964 | 29 |
| Estrogen Receptor Signaling | 2.38 | 0.173 | 3.048 | 71 |
| Dilated Cardiomyopathy Signaling Pathway | 2.35 | 0.205 | -0.2 | 31 |
| TGF-β Signaling | 2.35 | 0.227 | 1.5 | 22 |
| Chondroitin Sulfate Biosynthesis (Late Stages) | 2.34 | 0.254 | 1.897 | 16 |
| Stearate Biosynthesis I (Animals) | 2.34 | 0.254 | -1.265 | 16 |
| Xenobiotic Metabolism AHR Signaling Pathway | 2.34 | 0.22 | 0 | 24 |
| Gαs Signaling | 2.33 | 0.213 | -0.688 | 27 |
| Transcriptional Regulatory Network in Embryonic Stem Cells | 2.33 | 0.201 | 0.87 | 33 |
| Nuclear Cytoskeleton Signaling Pathway | 2.33 | 0.19 | 3.363 | 42 |
| Ketone body metabolism | 2.32 | 0.5 | -0.447 | 5 |
| Epithelial Adherens Junction Signaling | 2.32 | 0.203 | 1.976 | 32 |
| rRNA processing | 2.31 | 0.312 | 3.162 | 10 |
| Extra-nuclear estrogen signaling | 2.29 | 0.24 | 2.828 | 18 |
| Endocannabinoid Cancer Inhibition Pathway | 2.26 | 0.204 | -1.3 | 30 |
| Azathioprine ADME | 2.25 | 0.348 | 0 | 8 |
| NFE2L2 regulating anti-oxidant/detoxification enzymes | 2.25 | 0.348 | 1.414 | 8 |
| Post-translational modification: synthesis of GPI-anchored proteins | 2.24 | 0.226 | -0.218 | 21 |
| Role of Pattern Recognition Receptors in Recognition of Bacteria and Viruses | 2.22 | 0.201 | 2.324 | 31 |
| Macropinocytosis Signaling | 2.22 | 0.237 | 1.508 | 18 |
| Metabolism of steroid hormones | 2.21 | 0.289 | -0.302 | 11 |
| TNFR2 Signaling | 2.21 | 0.303 | 1.414 | 10 |
| BEX2 Signaling Pathway | 2.21 | 0.232 | -0.471 | 19 |
| Role of JAK2 in Hormone-like Cytokine Signaling | 2.2 | 0.246 | -1.732 | 16 |
| CCR3 Signaling in Eosinophils | 2.2 | 0.206 | 1 | 28 |
| Apelin Adipocyte Signaling Pathway | 2.19 | 0.223 | -0.447 | 21 |
| Apoptosis Signaling | 2.18 | 0.217 | 0.688 | 23 |
| γ-linolenate Biosynthesis II (Animals) | 2.18 | 0.368 | -1.134 | 7 |
| Bile Acid Biosynthesis, Neutral Pathway | 2.18 | 0.368 | -2 | 7 |
| G alpha (q) signalling events | 2.18 | 0.195 | 0.686 | 34 |
| Tuberculosis Active Signaling Pathway | 2.17 | 0.184 | 0 | 45 |
| Multiple Sclerosis Signaling Pathway | 2.17 | 0.187 | 1.718 | 41 |
| TP53 Regulates Transcription of Cell Cycle Genes | 2.17 | 0.265 | 3.606 | 13 |
| PFKFB4 Signaling Pathway | 2.17 | 0.265 | 0.905 | 13 |
| Angiopoietin Signaling | 2.16 | 0.234 | 0.632 | 18 |
| Relaxin Signaling | 2.14 | 0.199 | 0.447 | 31 |
| Netrin-1 signaling | 2.14 | 0.273 | 0 | 12 |
| Catecholamine Biosynthesis | 2.13 | 0.75 | - | 3 |
| Interleukin-7 signaling | 2.13 | 0.333 | 0 | 8 |
| Fatty Acid Activation | 2.13 | 0.4 | -0.816 | 6 |
| Sensory processing of sound by outer hair cells of the cochlea | 2.12 | 0.255 | -0.535 | 14 |
| SUMOylation of intracellular receptors | 2.11 | 0.31 | -2.333 | 9 |
| Role of p14/p19ARF in Tumor Suppression | 2.11 | 0.31 | -1.134 | 9 |
| Phenylalanine and tyrosine metabolism | 2.11 | 0.455 | -2.236 | 5 |
| Aspartate and asparagine metabolism | 2.11 | 0.455 | -1.342 | 5 |
| NFE2L2 regulating tumorigenic genes | 2.11 | 0.455 | 2.236 | 5 |
| Glycine Betaine Degradation | 2.11 | 0.455 | -2.236 | 5 |
| 14-3-3-mediated Signaling | 2.1 | 0.205 | 1.414 | 27 |
| Melanoma Signaling | 2.09 | 0.26 | 0.905 | 13 |
| Role of MAPK Signaling in the Pathogenesis of Influenza | 2.09 | 0.226 | - | 19 |
| NAD Signaling Pathway | 2.09 | 0.199 | 0.756 | 30 |
| MSP-RON Signaling in Macrophages Pathway | 2.09 | 0.208 | 1.877 | 25 |
| Neutrophil Extracellular Trap Signaling Pathway | 2.09 | 0.169 | 0.985 | 69 |
| Role of MAPK Signaling in Promoting the Pathogenesis of Influenza | 2.08 | 0.211 | 1.279 | 24 |
| Prolactin Signaling | 2.08 | 0.219 | 0.535 | 21 |
| Apelin Endothelial Signaling Pathway | 2.08 | 0.2 | 0.853 | 29 |
| White Adipose Tissue Browning Pathway | 2.07 | 0.201 | 0.2 | 28 |
| CD27 Signaling in Lymphocytes | 2.05 | 0.25 | 1.265 | 14 |
| Cyclophilin Signaling Pathway | 2.05 | 0.181 | 3.13 | 45 |
| Ceramide Signaling | 2.03 | 0.22 | 1 | 20 |
| Mouse Embryonic Stem Cell Pluripotency | 2.03 | 0.214 | 0.655 | 22 |
| GABAergic Receptor Signaling Pathway (Enhanced) | 2.03 | 0.2 | -0.192 | 28 |
| Gap junction trafficking and regulation | 2.01 | 0.255 | 1.387 | 13 |
| Gustation Pathway | 2 | 0.185 | -0.333 | 39 |
| Gαi Signaling | 2 | 0.197 | -1.279 | 29 |
| Sleep NREM Signaling Pathway | 1.99 | 0.207 | -0.408 | 24 |
| Signaling by Retinoic Acid | 1.98 | 0.261 | -2.309 | 12 |
| WNT/β-catenin Signaling | 1.97 | 0.191 | 0.392 | 33 |
| Platelet Adhesion to exposed collagen | 1.97 | 0.375 | 0.816 | 6 |
| Superpathway of Citrulline Metabolism | 1.97 | 0.375 | -1.633 | 6 |
| Oxytocin Signaling Pathway | 1.97 | 0.175 | 2.48 | 50 |
| DAG and IP3 signaling | 1.95 | 0.268 | 0.632 | 11 |
| tRNA processing in the mitochondrion | 1.95 | 0.268 | 3.317 | 11 |
| RET signaling | 1.95 | 0.268 | 0.302 | 11 |
| Cholecystokinin/Gastrin-mediated Signaling | 1.94 | 0.205 | 1.46 | 24 |
| α-Adrenergic Signaling | 1.94 | 0.207 | 0 | 23 |
| Melanocyte Development and Pigmentation Signaling | 1.93 | 0.212 | 1.213 | 21 |
| Formation of intermediate mesoderm | 1.93 | 0.5 | 0 | 4 |
| NFE2L2 regulating MDR associated enzymes | 1.93 | 0.5 | 1 | 4 |
| Vitamin D (calciferol) metabolism | 1.92 | 0.417 | -0.447 | 5 |
| Oleate Biosynthesis II (Animals) | 1.92 | 0.417 | -1.342 | 5 |
| Tight Junction Signaling | 1.92 | 0.188 | - | 34 |
| Netrin Signaling | 1.9 | 0.189 | 0.898 | 33 |
| Metabolism of porphyrins | 1.9 | 0.308 | -1.414 | 8 |
| ERB2-ERBB3 Signaling | 1.9 | 0.234 | 1.387 | 15 |
| Chronic Myeloid Leukemia Signaling | 1.89 | 0.174 | 4.907 | 49 |
| ABRA Signaling Pathway | 1.87 | 0.216 | 2.065 | 19 |
| Signaling by EGFR | 1.87 | 0.245 | 1.387 | 13 |
| Xenobiotic Metabolism General Signaling Pathway | 1.87 | 0.19 | 0.816 | 31 |
| Adrenomedullin signaling pathway | 1.87 | 0.183 | 0.343 | 37 |
| Th1 and Th2 Activation Pathway | 1.87 | 0.188 | - | 33 |
| Orexin Signaling Pathway | 1.86 | 0.178 | 0.926 | 43 |
| Role of NFAT in Cardiac Hypertrophy | 1.85 | 0.179 | 1 | 41 |
| Detoxification of Reactive Oxygen Species | 1.84 | 0.27 | 1.265 | 10 |
| Neuropathic Pain Signaling in Dorsal Horn Neurons | 1.83 | 0.208 | -0.218 | 21 |
| Mitochondrial Dysfunction | 1.82 | 0.168 | -0.687 | 58 |
| Surfactant metabolism | 1.81 | 0.281 | 0.333 | 9 |
| Signaling by TGF-beta Receptor Complex | 1.8 | 0.241 | 2.496 | 13 |
| UVB-Induced MAPK Signaling | 1.8 | 0.241 | 1.155 | 13 |
| Effects of PIP2 hydrolysis | 1.8 | 0.296 | 0.707 | 8 |
| BMAL1:CLOCK,NPAS2 activates circadian gene expression | 1.8 | 0.296 | 0.707 | 8 |
| Integrin signaling | 1.8 | 0.296 | 0.707 | 8 |
| GABA Receptor Signaling | 1.8 | 0.195 | - | 26 |
| Glycine Degradation (Creatine Biosynthesis) | 1.79 | 1 | - | 2 |
| Glutamine Degradation I | 1.79 | 1 | - | 2 |
| Integration of energy metabolism | 1.79 | 0.204 | 1.706 | 22 |
| PPAR Signaling | 1.79 | 0.204 | -1.342 | 22 |
| Transport of vitamins, nucleosides, and related molecules | 1.79 | 0.256 | -0.302 | 11 |
| Melanin biosynthesis | 1.78 | 0.6 | - | 3 |
| Intestinal absorption | 1.78 | 0.6 | - | 3 |
| Lysine Degradation II | 1.78 | 0.6 | - | 3 |
| Lysine Degradation V | 1.78 | 0.6 | - | 3 |
| Tyrosine Degradation I | 1.78 | 0.6 | - | 3 |
| Renal Cell Carcinoma Signaling | 1.77 | 0.218 | 0.905 | 17 |
| iNOS Signaling | 1.76 | 0.245 | 2.53 | 12 |
| SUMOylation of immune response proteins | 1.76 | 0.385 | 0.447 | 5 |
| Autophagy | 1.75 | 0.178 | 0.324 | 39 |
| Death Receptor Signaling | 1.73 | 0.206 | 0.243 | 20 |
| IL-15 Production | 1.73 | 0.197 | 1.46 | 24 |
| PI3K Cascade | 1.72 | 0.25 | 0.302 | 11 |
| TAK1-dependent IKK and NF-kappa-B activation | 1.72 | 0.25 | 0.905 | 11 |
| Assembly and cell surface presentation of NMDA receptors | 1.72 | 0.25 | 0.905 | 11 |
| Interleukin-1 processing | 1.72 | 0.444 | 2 | 4 |
| Sphingosine and Sphingosine-1-phosphate Metabolism | 1.72 | 0.444 | -1 | 4 |
| Caveolar-mediated Endocytosis Signaling | 1.72 | 0.215 | 3.207 | 17 |
| Sphingolipid metabolism | 1.71 | 0.2 | 1.667 | 22 |
| Interleukin-1 family signaling | 1.7 | 0.194 | 1.043 | 25 |
| Clathrin-mediated endocytosis | 1.7 | 0.194 | 0.6 | 25 |
| Endothelin-1 Signaling | 1.7 | 0.18 | 0.557 | 35 |
| MYC Mediated Apoptosis Signaling | 1.69 | 0.24 | -1.508 | 12 |
| Retinoate Biosynthesis I | 1.69 | 0.24 | -2 | 12 |
| Nephrin family interactions | 1.69 | 0.304 | 1.134 | 7 |
| MAP kinase activation | 1.68 | 0.256 | 0.632 | 10 |
| Acute Myeloid Leukemia Signaling | 1.68 | 0.207 | 0.775 | 19 |
| Antioxidant Action of Vitamin C | 1.68 | 0.197 | -0.577 | 23 |
| Cyclins and Cell Cycle Regulation | 1.67 | 0.209 | 0 | 18 |
| Cargo recognition for clathrin-mediated endocytosis | 1.65 | 0.2 | 0.218 | 21 |
| Neurexins and neuroligins | 1.62 | 0.228 | -1.387 | 13 |
| Carnitine metabolism | 1.61 | 0.357 | -0.447 | 5 |
| Glutamate and glutamine metabolism | 1.61 | 0.357 | -0.447 | 5 |
| Glycolysis I | 1.61 | 0.276 | -0.378 | 8 |
| Transcriptional Regulation by MECP2 | 1.61 | 0.222 | -1.604 | 14 |
| Sensory processing of sound by inner hair cells of the cochlea | 1.61 | 0.217 | 0.258 | 15 |
| Cell Cycle: G1/S Checkpoint Regulation | 1.61 | 0.217 | -2.111 | 15 |
| Protein Kinase A Signaling | 1.6 | 0.162 | 1.508 | 64 |
| Sphingosine-1-phosphate Signaling | 1.6 | 0.193 | 1.342 | 23 |
| Tryptophan Degradation III (Eukaryotic) | 1.59 | 0.292 | -2.449 | 7 |
| G alpha (s) signalling events | 1.59 | 0.186 | -0.192 | 27 |
| Calcium Signaling | 1.59 | 0.174 | 0.2 | 38 |
| Mitochondrial L-carnitine Shuttle Pathway | 1.58 | 0.316 | -0.816 | 6 |
| Acetylcholine Receptor Signaling Pathway | 1.58 | 0.177 | 0.169 | 35 |
| GPVI-mediated activation cascade | 1.56 | 0.257 | 1 | 9 |
| Semaphorin interactions | 1.55 | 0.219 | 2.138 | 14 |
| Peroxisomal protein import | 1.55 | 0.219 | -3.742 | 14 |
| Caspase activation via Dependence Receptors in the absence of ligand | 1.54 | 0.4 | 1 | 4 |
| Glycogen Degradation II | 1.54 | 0.4 | - | 4 |
| Citrulline Biosynthesis | 1.54 | 0.4 | -1 | 4 |
| FOXO-mediated transcription of oxidative stress, metabolic and neuronal genes | 1.53 | 0.267 | -1.414 | 8 |
| NFE2L2 regulating inflammation associated genes | 1.52 | 0.5 | - | 3 |
| Arginine Biosynthesis IV | 1.52 | 0.5 | - | 3 |
| Chondroitin and Dermatan Biosynthesis | 1.52 | 0.5 | - | 3 |
| Tryptophan Degradation to 2-amino-3-carboxymuconate Semialdehyde | 1.52 | 0.5 | - | 3 |
| Basal Cell Carcinoma Signaling | 1.51 | 0.211 | 0.333 | 15 |
| Semaphorin Signaling in Neurons | 1.5 | 0.22 | - | 13 |
| Oxytocin in Brain Signaling Pathway | 1.49 | 0.174 | 0.174 | 35 |
| BMP signaling pathway | 1.49 | 0.2 | 0.258 | 18 |
| Macrophage Classical Activation Signaling Pathway | 1.48 | 0.176 | 3.889 | 33 |
| Receptor-type tyrosine-protein phosphatases | 1.48 | 0.3 | -2.449 | 6 |
| Synthesis, secretion, and deacylation of Ghrelin | 1.48 | 0.3 | -0.816 | 6 |
| Valine Degradation I | 1.48 | 0.3 | -1.633 | 6 |
| Glucocorticoid Biosynthesis | 1.48 | 0.3 | - | 6 |
| Apelin Liver Signaling Pathway | 1.48 | 0.3 | 0.816 | 6 |
| Circadian Rhythm Signaling | 1.47 | 0.167 | - | 45 |
| Insulin Receptor Signaling | 1.46 | 0.186 | 0.853 | 24 |
| Induction of Apoptosis by HIV1 | 1.45 | 0.212 | 1.155 | 14 |
| Oxidative Ethanol Degradation III | 1.45 | 0.212 | -1 | 14 |
| Dermatan Sulfate Biosynthesis (Late Stages) | 1.45 | 0.217 | 1.134 | 13 |
| Natural Killer Cell Signaling | 1.44 | 0.172 | 2.263 | 35 |
| Neuroprotective Role of THOP1 in Alzheimer's Disease | 1.43 | 0.185 | 0 | 24 |
| Growth Hormone Signaling | 1.41 | 0.205 | -0.277 | 15 |
| Pregnenolone Biosynthesis | 1.41 | 0.269 | - | 7 |
| Ethanol Degradation IV | 1.41 | 0.269 | -0.816 | 7 |
| Abacavir ADME | 1.4 | 0.209 | -0.535 | 14 |
| Intrinsic Pathway for Apoptosis | 1.39 | 0.218 | 2.887 | 12 |
| Signaling by PTK6 | 1.39 | 0.218 | 1.732 | 12 |
| Regulation of the Epithelial Mesenchymal Transition in Development Pathway | 1.39 | 0.198 | 2.324 | 17 |
| Branched-chain amino acid catabolism | 1.38 | 0.286 | -1.633 | 6 |
| RIPK1-mediated regulated necrosis | 1.37 | 0.25 | 0.707 | 8 |
| ERK5 Signaling | 1.36 | 0.203 | 3.051 | 15 |
| RHO GTPases regulate CFTR trafficking | 1.35 | 0.667 | - | 2 |
| Interconversion of 2-oxoglutarate and 2-hydroxyglutarate | 1.35 | 0.667 | - | 2 |
| Serine Biosynthesis | 1.35 | 0.667 | - | 2 |
| Tyrosine Biosynthesis IV | 1.35 | 0.667 | - | 2 |
| Cell Cycle Regulation by BTG Family Proteins | 1.35 | 0.237 | 1.89 | 9 |
| TWEAK Signaling | 1.35 | 0.237 | 1.134 | 9 |
| MSP-RON Signaling Pathway | 1.34 | 0.21 | - | 13 |
| Thyroid Hormone Metabolism II (via Conjugation and/or Degradation) | 1.34 | 0.21 | -1.134 | 13 |
| Signaling by ERBB2 | 1.34 | 0.22 | 1.508 | 11 |
| Signaling by TGFBR3 | 1.34 | 0.22 | 2.111 | 11 |
| Chemokine Signaling | 1.33 | 0.198 | 0.277 | 16 |
| fMLP Signaling in Neutrophils | 1.33 | 0.18 | 1.698 | 24 |
| Histidine Degradation VI | 1.33 | 0.259 | -1 | 7 |
| Erythrocytes take up oxygen and release carbon dioxide | 1.32 | 0.429 | - | 3 |
| Ceramide Degradation | 1.32 | 0.429 | - | 3 |
| Heparan Sulfate Biosynthesis | 1.3 | 0.188 | 0.905 | 19 |
| Sumoylation Pathway | 1.3 | 0.188 | -0.258 | 19 |
| Airway Inflammation in Asthma | 1.3 | 0.242 | 2.236 | 8 |
| Inhibition of Angiogenesis by TSP1 | 1.3 | 0.242 | 0.816 | 8 |
| IL-27 Signaling Pathway | 1.3 | 0.179 | 2.041 | 24 |

Table S8: The top canonical pathways enriched in AH/ASH by DEG analysis

| Ingenuity Canonical Pathways | -log(p-value) | Ratio | z-score | Gene count |
| --- | --- | --- | --- | --- |
| Fcgamma receptor (FCGR) dependent phagocytosis | 10.4 | 0.11 | 4.243 | 18 |
| Complement cascade | 9.77 | 0.118 | 3 | 16 |
| Binding and Uptake of Ligands by Scavenger Receptors | 8.87 | 0.123 | 3.207 | 14 |
| Immunoregulatory interactions between a Lymphoid and a non-Lymphoid cell | 7.67 | 0.0791 | 3.638 | 17 |
| Fc epsilon receptor (FCERI) signaling | 5.63 | 0.068 | 3.742 | 14 |
| Cell surface interactions at the vascular wall | 5.44 | 0.0654 | 3.742 | 14 |
| Mitochondrial Fatty Acid Beta-Oxidation | 4.83 | 0.162 | -1.633 | 6 |
| Stearate Biosynthesis I (Animals) | 4.43 | 0.111 | - | 7 |
| Signaling by the B Cell Receptor (BCR) | 4.36 | 0.0647 | 3.317 | 11 |
| Remodeling of Epithelial Adherens Junctions | 4.09 | 0.0986 | - | 7 |
| RHO GTPases activate IQGAPs | 4.02 | 0.156 | 2.236 | 5 |
| Fatty Acid β-oxidation I | 3.89 | 0.147 | - | 5 |
| eNOS Signaling | 3.86 | 0.0621 | 1.342 | 10 |
| Response to elevated platelet cytosolic Ca2+ | 3.84 | 0.0682 | 1 | 9 |
| Transport of glycerol from adipocytes to the liver by Aquaporins | 3.67 | 1 | - | 2 |
| Inhibition of Matrix Metalloproteases | 3.66 | 0.132 | -0.447 | 5 |
| IL-15 Signaling | 3.5 | 0.0361 | - | 19 |
| Leukocyte Extravasation Signaling | 3.22 | 0.0515 | 1.414 | 10 |
| Translocation of SLC2A4 (GLUT4) to the plasma membrane | 3.19 | 0.0833 | 1.633 | 6 |
| LXR/RXR Activation | 3.17 | 0.0615 | -2.449 | 8 |
| Paracetamol ADME | 3.1 | 0.138 | -1 | 4 |
| Gap junction trafficking and regulation | 3.05 | 0.098 | 2.236 | 5 |
| FcγRIIB Signaling in B Lymphocytes | 3.02 | 0.0338 | - | 18 |
| p70S6K Signaling | 2.99 | 0.0327 | - | 19 |
| HIF1α Signaling | 2.95 | 0.0476 | 1.667 | 10 |
| Phase II - Conjugation of compounds | 2.95 | 0.0642 | -1.134 | 7 |
| DHCR24 Signaling Pathway | 2.9 | 0.0559 | -1.134 | 8 |
| NAD Biosynthesis III | 2.9 | 0.5 | - | 2 |
| PI3K Signaling in B Lymphocytes | 2.9 | 0.0321 | - | 19 |
| HSP90 chaperone cycle for steroid hormone receptors in the presence of ligand | 2.83 | 0.0877 | 2.236 | 5 |
| Xenobiotic Metabolism PXR Signaling Pathway | 2.79 | 0.0452 | -1.134 | 10 |
| Intestinal absorption | 2.68 | 0.4 | - | 2 |
| Chondroitin Sulfate Biosynthesis (Late Stages) | 2.64 | 0.0794 | - | 5 |
| Xenobiotic Metabolism CAR Signaling Pathway | 2.62 | 0.0429 | 0.378 | 10 |
| Semaphorin interactions | 2.61 | 0.0781 | 2.236 | 5 |
| B Cell Development | 2.59 | 0.0326 | - | 16 |
| Systemic Lupus Erythematosus in B Cell Signaling Pathway | 2.58 | 0.0288 | 2.236 | 21 |
| Human Embryonic Stem Cell Pluripotency | 2.55 | 0.045 | 1.667 | 9 |
| B Cell Receptor Signaling | 2.53 | 0.0298 | - | 19 |
| Pulmonary Healing Signaling Pathway | 2.53 | 0.0448 | 2.333 | 9 |
| Germ Cell-Sertoli Cell Junction Signaling | 2.42 | 0.0468 | - | 8 |
| Chondroitin Sulfate Biosynthesis | 2.41 | 0.0704 | - | 5 |
| Aggrephagy | 2.41 | 0.0909 | 2 | 4 |
| Chaperone Mediated Autophagy | 2.41 | 0.136 | - | 3 |
| Dermatan Sulfate Biosynthesis | 2.36 | 0.0685 | - | 5 |
| RAF-independent MAPK1/3 activation | 2.35 | 0.13 | - | 3 |
| Neutrophil degranulation | 2.32 | 0.0314 | 2.84 | 15 |
| Tryptophan Degradation III (Eukaryotic) | 2.3 | 0.125 | - | 3 |
| Tumor Microenvironment Pathway | 2.29 | 0.0444 | 1.89 | 8 |
| NF-κB Activation by Viruses | 2.24 | 0.0641 | 2 | 5 |
| Maturity Onset Diabetes of Young (MODY) Signaling | 2.24 | 0.0641 | - | 5 |
| RHO GTPase cycle | 2.16 | 0.0311 | 3.742 | 14 |
| NAD Signaling Pathway | 2.16 | 0.0464 | -1.134 | 7 |
| Lipophagy | 2.14 | 0.222 | - | 2 |
| Neuregulin Signaling | 2.1 | 0.0508 | - | 6 |
| Integrin cell surface interactions | 2.08 | 0.0588 | 0.447 | 5 |
| Activation of NMDA receptors and postsynaptic events | 2.08 | 0.0588 | 1.342 | 5 |
| Phospholipase C Signaling | 2.08 | 0.024 | 2.828 | 27 |
| Osteoarthritis Pathway | 2.06 | 0.0378 | 0.816 | 9 |
| TRIM21 Intracellular Antibody Signaling Pathway | 2.01 | 0.0282 | 4 | 16 |
| L1CAM interactions | 2 | 0.0484 | 2.449 | 6 |
| NAP1L1 Transcription Regulation Signaling Pathway | 2 | 0.0562 | 2.236 | 5 |
| Regulation of Cellular Mechanics by Calpain Protease | 2 | 0.0562 | - | 5 |
| Xenobiotic Metabolism Signaling | 2 | 0.0332 | - | 11 |
| Triacylglycerol Biosynthesis | 1.99 | 0.069 | - | 4 |
| Th1 Pathway | 1.99 | 0.048 | 2.236 | 6 |
| Granulocyte Adhesion and Diapedesis | 1.97 | 0.0392 | - | 8 |
| TCR signaling | 1.97 | 0.0476 | 2.449 | 6 |
| Aspartate and asparagine metabolism | 1.97 | 0.182 | - | 2 |
| NAD biosynthesis II (from tryptophan) | 1.97 | 0.182 | - | 2 |
| Cell junction organization | 1.94 | 0.0543 | 2.236 | 5 |
| Dermatan Sulfate Biosynthesis (Late Stages) | 1.94 | 0.0667 | - | 4 |
| Reversible hydration of carbon dioxide | 1.89 | 0.167 | - | 2 |
| Lysine catabolism | 1.89 | 0.167 | - | 2 |
| Oleate Biosynthesis II (Animals) | 1.89 | 0.167 | - | 2 |
| LPS/IL-1 Mediated Inhibition of RXR Function | 1.89 | 0.0336 | - | 10 |
| Kinesins | 1.89 | 0.0645 | 2 | 4 |
| Primary Immunodeficiency Signaling | 1.89 | 0.0645 | - | 4 |
| Aldosterone Signaling in Epithelial Cells | 1.88 | 0.0409 | - | 7 |
| Hematoma Resolution Signaling Pathway | 1.84 | 0.0347 | 0.707 | 9 |
| Methylthiopropionate Biosynthesis | 1.83 | 1 | - | 1 |
| Methylglyoxal Degradation VI | 1.83 | 1 | - | 1 |
| Adenine and Adenosine Salvage VI | 1.83 | 1 | - | 1 |
| Passive transport by Aquaporins | 1.83 | 0.154 | - | 2 |
| Nuclear Cytoskeleton Signaling Pathway | 1.78 | 0.0362 | 2.828 | 8 |
| Acetylcholine binding and downstream events | 1.76 | 0.143 | - | 2 |
| Agranulocyte Adhesion and Diapedesis | 1.75 | 0.0357 | - | 8 |
| FLT3 Signaling | 1.72 | 0.0769 | - | 3 |
| Fatty Acid Activation | 1.7 | 0.133 | - | 2 |
| Paxillin Signaling | 1.7 | 0.0472 | 2 | 5 |
| Role of NFAT in Regulation of the Immune Response | 1.68 | 0.0229 | 2.646 | 24 |
| Axonal Guidance Signaling | 1.68 | 0.0271 | - | 14 |
| Mechanisms of Viral Exit from Host Cells | 1.66 | 0.0732 | - | 3 |
| Hepatic Fibrosis / Hepatic Stellate Cell Activation | 1.63 | 0.0365 | - | 7 |
| DAP12 interactions | 1.6 | 0.0698 | - | 3 |
| Glutaryl-CoA Degradation | 1.6 | 0.118 | - | 2 |
| Interferon alpha/beta signaling | 1.6 | 0.0526 | 1 | 4 |
| Pathogen Induced Cytokine Storm Signaling Pathway | 1.59 | 0.0288 | 0.707 | 11 |
| Assembly and cell surface presentation of NMDA receptors | 1.58 | 0.0682 | - | 3 |
| Aspirin ADME | 1.58 | 0.0682 | - | 3 |
| IL-12 Signaling and Production in Macrophages | 1.57 | 0.0331 | 1.633 | 8 |
| Carboxyterminal post-translational modifications of tubulin | 1.55 | 0.0667 | - | 3 |
| Mitotic G2-G2/M phases | 1.55 | 0.035 | 2.646 | 7 |
| Asparagine Degradation I | 1.54 | 0.5 | - | 1 |
| β-alanine Degradation I | 1.54 | 0.5 | - | 1 |
| Glutamate Biosynthesis II | 1.54 | 0.5 | - | 1 |
| Glutamate Degradation X | 1.54 | 0.5 | - | 1 |
| Epithelial Adherens Junction Signaling | 1.54 | 0.038 | -0.816 | 6 |
| Sertoli Cell-Sertoli Cell Junction Signaling | 1.53 | 0.0324 | 2.828 | 8 |
| Natural Killer Cell Signaling | 1.52 | 0.0345 | 2.449 | 7 |
| γ-linolenate Biosynthesis II (Animals) | 1.51 | 0.105 | - | 2 |
| Mitochondrial L-carnitine Shuttle Pathway | 1.51 | 0.105 | - | 2 |
| Sheddase Signaling Pathway | 1.51 | 0.0343 | 0.378 | 7 |
| Role of JAK family kinases in IL-6-type Cytokine Signaling | 1.51 | 0.0494 | 1 | 4 |
| Interleukin-2 family signaling | 1.5 | 0.0638 | - | 3 |
| IL-15 Production | 1.47 | 0.041 | 2.236 | 5 |
| Integrin Signaling | 1.45 | 0.0333 | 2.449 | 7 |
| Sperm Motility | 1.44 | 0.0311 | - | 8 |
| RAF/MAP kinase cascade | 1.4 | 0.0305 | 2.121 | 8 |
| Glycosaminoglycan metabolism | 1.4 | 0.0391 | 2.236 | 5 |
| MHC class II antigen presentation | 1.4 | 0.0391 | 2.236 | 5 |
| GP6 Signaling Pathway | 1.4 | 0.0391 | 1 | 5 |
| Cellular hexose transport | 1.39 | 0.0909 | - | 2 |
| Regulation of TLR by endogenous ligand | 1.39 | 0.0909 | - | 2 |
| Clathrin-mediated endocytosis | 1.38 | 0.0388 | 0.447 | 5 |
| RHO GTPases regulate CFTR trafficking | 1.36 | 0.333 | - | 1 |
| Pyrophosphate hydrolysis | 1.36 | 0.333 | - | 1 |
| Uracil Degradation II (Reductive) | 1.36 | 0.333 | - | 1 |
| Thymine Degradation | 1.36 | 0.333 | - | 1 |
| Anandamide Degradation | 1.36 | 0.333 | - | 1 |
| Nephrin family interactions | 1.36 | 0.087 | - | 2 |
| Nucleotide salvage | 1.36 | 0.087 | - | 2 |
| Th1 and Th2 Activation Pathway | 1.34 | 0.0341 | - | 6 |
| NLR signaling pathways | 1.31 | 0.0536 | - | 3 |
| Heparan Sulfate Biosynthesis (Late Stages) | 1.31 | 0.0426 | - | 4 |
| Molecular Mechanisms of Cancer | 1.3 | 0.0221 | 3.771 | 19 |

Table S9: The top canonical pathways enriched in AH/ASH by DSD analysis

| Ingenuity Canonical Pathways | -log(p-value) | Ratio | z-score | Gene count |
| --- | --- | --- | --- | --- |
| CREB Signaling in Neurons | 6.15 | 0.0325 | -3.9 | 20 |
| BBSome Signaling Pathway | 5.65 | 0.0345 | -3.5 | 17 |
| Lung Ionic Balance Signaling Pathway | 5.32 | 0.0326 | -3.638 | 17 |
| S100 Family Signaling Pathway | 5.13 | 0.0269 | -4.025 | 21 |
| Breast Cancer Regulation by Stathmin1 | 5.05 | 0.0297 | -3.638 | 18 |
| Cellular Effects of Sildenafil (Viagra) | 4.35 | 0.0263 | 2.828 | 18 |
| Phagosome Formation | 4.18 | 0.0255 | -3.638 | 18 |
| G-Protein Coupled Receptor Signaling | 4.13 | 0.0252 | -3.638 | 18 |
| Molecular Mechanisms of Cancer | 4.03 | 0.0232 | -3.638 | 20 |
| Melanin biosynthesis | 3.11 | 0.4 | - | 2 |
| Class A/1 (Rhodopsin-like receptors) | 3.09 | 0.0301 | -3.162 | 10 |
| Neurotransmitter clearance | 2.47 | 0.2 | - | 2 |
| Oleate Biosynthesis II (Animals) | 2.31 | 0.167 | - | 2 |
| FAK Signaling | 2.27 | 0.0174 | -3.638 | 18 |
| Cardiac conduction | 2.21 | 0.0385 | -2.236 | 5 |
| Neuroprotective Role of THOP1 in Alzheimer's Disease | 2.21 | 0.0385 | -1 | 5 |
| TR/RXR Activation | 2.2 | 0.0382 | -0.447 | 5 |
| Platelet homeostasis | 2.14 | 0.0465 | -1 | 4 |
| G alpha (i) signalling events | 2.07 | 0.0271 | -1.89 | 7 |
| Lanosterol Biosynthesis | 2.05 | 1 | - | 1 |
| Gαi Signaling | 1.99 | 0.034 | -1 | 5 |
| Metabolism of amine-derived hormones | 1.96 | 0.111 | - | 2 |
| γ-linolenate Biosynthesis II (Animals) | 1.92 | 0.105 | - | 2 |
| Potassium Channels | 1.87 | 0.0388 | -1 | 4 |
| Cellular hexose transport | 1.79 | 0.0909 | - | 2 |
| L-dopachrome Biosynthesis | 1.75 | 0.5 | - | 1 |
| G alpha (q) signalling events | 1.7 | 0.0287 | -2.236 | 5 |
| cAMP-mediated signaling | 1.68 | 0.0249 | -1.342 | 6 |
| Cholesterol biosynthesis | 1.62 | 0.0741 | - | 2 |
| Serotonin Receptor Signaling | 1.6 | 0.0191 | -1.667 | 9 |
| NFE2L2 regulating TCA cycle genes | 1.58 | 0.333 | - | 1 |
| Thyronamine and Iodothyronamine Metabolism | 1.58 | 0.333 | - | 1 |
| Thyroid Hormone Metabolism I (via Deiodination) | 1.58 | 0.333 | - | 1 |
| Superpathway of Cholesterol Biosynthesis | 1.51 | 0.0645 | - | 2 |
| Circadian Rhythm Signaling | 1.47 | 0.0222 | - | 6 |
| Eumelanin Biosynthesis | 1.46 | 0.25 | - | 1 |
| Degradation of the extracellular matrix | 1.45 | 0.037 | - | 3 |
| Serotonin and Melatonin Biosynthesis | 1.36 | 0.2 | - | 1 |
| Citrulline-Nitric Oxide Cycle | 1.36 | 0.2 | - | 1 |
| Tuberculosis Latent Signaling Pathway | 1.34 | 0.0333 | - | 3 |
| Neurotransmitter release cycle | 1.31 | 0.05 | - | 2 |

Table S10: The top common pathways enriched in AH/ASH by DEG and DSD analysis

| Ingenuity Canonical Pathways | -log(p-value) | Ratio | z-score | Gene count |
| --- | --- | --- | --- | --- |
| Extracellular matrix organization | 19.4 | 0.421 | 6.112 | 45 |
| Phase I - Functionalization of compounds | 16.6 | 0.381 | -4.529 | 43 |
| Hepatic Fibrosis / Hepatic Stellate Cell Activation | 16 | 0.297 | - | 57 |
| Collagen degradation | 15.7 | 0.484 | 4.49 | 31 |
| Assembly of collagen fibrils and other multimeric structures | 15.5 | 0.492 | 5.112 | 30 |
| Sheddase Signaling Pathway | 15.3 | 0.284 | 2.364 | 58 |
| Pulmonary Fibrosis Idiopathic Signaling Pathway | 15 | 0.236 | 6.437 | 77 |
| Wound Healing Signaling Pathway | 15 | 0.261 | 4.5 | 65 |
| Integrin cell surface interactions | 14.1 | 0.4 | 5.488 | 34 |
| S100 Family Signaling Pathway | 12.8 | 0.17 | 5.945 | 133 |
| Interleukin-4 and Interleukin-13 signaling | 11.6 | 0.324 | 4 | 36 |
| Role of Osteoclasts in Rheumatoid Arthritis Signaling Pathway | 10.8 | 0.211 | 4.202 | 67 |
| Post-translational protein phosphorylation | 10.7 | 0.318 | 3.43 | 34 |
| Elastic fibre formation | 10.7 | 0.477 | 4.146 | 21 |
| Regulation of Insulin-like Growth Factor (IGF) transport and uptake by IGFBPs | 10.7 | 0.298 | 3.452 | 37 |
| Molecular Mechanisms of Cancer | 10.1 | 0.156 | 5.371 | 134 |
| Tumor Microenvironment Pathway | 9.94 | 0.25 | 4.841 | 45 |
| Collagen chain trimerization | 9.74 | 0.455 | 3.578 | 20 |
| Collagen biosynthesis and modifying enzymes | 8.96 | 0.358 | 3.674 | 24 |
| LPS/IL-1 Mediated Inhibition of RXR Function | 8.88 | 0.201 | 1.387 | 60 |
| Atherosclerosis Signaling | 8.82 | 0.265 | 2.191 | 36 |
| Degradation of the extracellular matrix | 8.46 | 0.321 | 4.315 | 26 |
| Hepatic Fibrosis Signaling Pathway | 8.26 | 0.178 | 4.131 | 74 |
| Osteoarthritis Pathway | 8.15 | 0.21 | 1.183 | 50 |
| Coagulation System | 7.96 | 0.457 | -0.258 | 16 |
| Syndecan interactions | 7.93 | 0.519 | 3.207 | 14 |
| IL-17A Signaling in Fibroblasts | 7.9 | 0.312 | 3.545 | 25 |
| LXR/RXR Activation | 7.66 | 0.254 | -2.353 | 33 |
| Cellular Effects of Sildenafil (Viagra) | 7.42 | 0.152 | -1.8 | 104 |
| Estrogen Biosynthesis | 7.4 | 0.383 | 0 | 18 |
| Axonal Guidance Signaling | 7.4 | 0.162 | - | 84 |
| FXR/RXR Activation | 7.28 | 0.215 | -0.324 | 42 |
| Signaling by PDGF | 7.25 | 0.345 | 4.025 | 20 |
| RAR Activation | 7.17 | 0.169 | -0.12 | 73 |
| Bupropion Degradation | 7.16 | 0.5 | 0.447 | 13 |
| ABRA Signaling Pathway | 6.98 | 0.284 | 4.2 | 25 |
| Bile acid and bile salt metabolism | 6.92 | 0.378 | -2.183 | 17 |
| Arachidonic acid metabolism | 6.92 | 0.378 | -1.5 | 17 |
| Binding and Uptake of Ligands by Scavenger Receptors | 6.83 | 0.254 | 2.785 | 29 |
| ILK Signaling | 6.7 | 0.208 | 4.004 | 41 |
| Nicotine Degradation II | 6.68 | 0.282 | -0.333 | 24 |
| Irritable Bowel Syndrome Signaling Pathway | 6.54 | 0.183 | 2.885 | 53 |
| G-Protein Coupled Receptor Signaling | 6.54 | 0.146 | 2.312 | 104 |
| Phagosome Formation | 6.49 | 0.146 | 4.724 | 103 |
| Inhibition of Matrix Metalloproteases | 6.47 | 0.395 | -1.155 | 15 |
| Xenobiotic Metabolism Signaling | 6.4 | 0.175 | - | 58 |
| Agranulocyte Adhesion and Diapedesis | 6.39 | 0.196 | - | 44 |
| Glycation Signaling Pathway | 6.33 | 0.196 | 3.92 | 44 |
| Formation of Fibrin Clot (Clotting Cascade) | 6.3 | 0.385 | 0.775 | 15 |
| Airway Pathology in Chronic Obstructive Pulmonary Disease | 6.28 | 0.246 | 1.89 | 28 |
| Transport of bile salts and organic acids, metal ions and amine compounds | 6.16 | 0.274 | -0.209 | 23 |
| Sertoli Cell-Germ Cell Junction Signaling Pathway (Enhanced) | 6.14 | 0.191 | -0.447 | 45 |
| Neuroinflammation Signaling Pathway | 6.09 | 0.173 | 3.244 | 57 |
| Cardiac Hypertrophy Signaling (Enhanced) | 6.09 | 0.153 | 3.75 | 82 |
| Role of Chondrocytes in Rheumatoid Arthritis Signaling Pathway | 6.03 | 0.222 | 3.651 | 32 |
| Ethanol Degradation II | 5.99 | 0.389 | -1.897 | 14 |
| Pathogen Induced Cytokine Storm Signaling Pathway | 5.94 | 0.165 | 4.557 | 63 |
| Xenobiotic Metabolism CAR Signaling Pathway | 5.9 | 0.189 | 0.2 | 44 |
| O-linked glycosylation | 5.83 | 0.239 | 4.426 | 27 |
| Serotonin Receptor Signaling | 5.77 | 0.155 | 2.926 | 73 |
| Hematoma Resolution Signaling Pathway | 5.75 | 0.181 | -2.714 | 47 |
| Class A/1 (Rhodopsin-like receptors) | 5.67 | 0.169 | 3.742 | 56 |
| Leukocyte Extravasation Signaling | 5.58 | 0.196 | 3.157 | 38 |
| Sertoli Cell-Sertoli Cell Junction Signaling | 5.58 | 0.182 | 3.13 | 45 |
| Aspirin ADME | 5.52 | 0.341 | -3.357 | 15 |
| Retinoid metabolism and transport | 5.52 | 0.341 | 0.775 | 15 |
| Dissolution of Fibrin Clot | 5.49 | 0.615 | 2.828 | 8 |
| Granulocyte Adhesion and Diapedesis | 5.44 | 0.191 | - | 39 |
| Formation of the ureteric bud | 5.38 | 0.476 | 0.707 | 10 |
| Preeclampsia Signaling Pathway | 5.37 | 0.205 | 2.611 | 33 |
| Nicotine Degradation III | 5.25 | 0.267 | 0 | 20 |
| NCAM signaling for neurite out-growth | 5.25 | 0.286 | 2.828 | 18 |
| GP6 Signaling Pathway | 5.22 | 0.219 | 4.536 | 28 |
| Acetone Degradation I (to Methylglyoxal) | 5.2 | 0.341 | 0.447 | 14 |
| Lung Ionic Balance Signaling Pathway | 5.19 | 0.148 | 2.393 | 77 |
| Regulation of the Epithelial Mesenchymal Transition by Growth Factors Pathway | 5.17 | 0.191 | 4.382 | 37 |
| DHCR24 Signaling Pathway | 5.16 | 0.21 | -1.671 | 30 |
| HIF1α Signaling | 5.13 | 0.186 | 2 | 39 |
| Regulation of Actin-based Motility by Rho | 4.96 | 0.225 | 2.668 | 25 |
| Superpathway of Melatonin Degradation | 4.96 | 0.242 | 0.707 | 22 |
| Response to elevated platelet cytosolic Ca2+ | 4.95 | 0.212 | 2.646 | 28 |
| Smooth Muscle Contraction | 4.93 | 0.326 | 3.207 | 14 |
| Melatonin Degradation I | 4.92 | 0.247 | 0.378 | 21 |
| Activation of Matrix Metalloproteinases | 4.86 | 0.364 | 2.309 | 12 |
| Colorectal Cancer Metastasis Signaling | 4.85 | 0.17 | 3.124 | 46 |
| Neurovascular Coupling Signaling Pathway | 4.81 | 0.177 | 1.298 | 41 |
| Regulation of the Epithelial-Mesenchymal Transition Pathway | 4.78 | 0.186 | - | 36 |
| Hepatic Cholestasis | 4.78 | 0.178 | 3.042 | 40 |
| Glycosaminoglycan metabolism | 4.75 | 0.211 | 2.2 | 27 |
| Noradrenaline and Adrenaline Degradation | 4.74 | 0.333 | -1.667 | 13 |
| Role of JAK family kinases in IL-6-type Cytokine Signaling | 4.71 | 0.247 | 1.886 | 20 |
| Bone Mineralization Signaling Pathway | 4.65 | 0.201 | 2.785 | 29 |
| RHOGDI Signaling | 4.64 | 0.177 | -3.4 | 39 |
| CDX Gastrointestinal Cancer Signaling Pathway | 4.63 | 0.183 | -1.715 | 36 |
| Xenobiotic Metabolism PXR Signaling Pathway | 4.6 | 0.176 | 0.209 | 39 |
| Breast Cancer Regulation by Stathmin1 | 4.57 | 0.139 | 4.472 | 84 |
| Aryl Hydrocarbon Receptor Signaling | 4.56 | 0.186 | 0 | 34 |
| IL-10 Signaling | 4.44 | 0.194 | 0.186 | 30 |
| Pulmonary Healing Signaling Pathway | 4.43 | 0.179 | 3.773 | 36 |
| CREB Signaling in Neurons | 4.31 | 0.136 | 3.85 | 84 |
| HOTAIR Regulatory Pathway | 4.31 | 0.188 | 3.402 | 31 |
| Role of Macrophages, Fibroblasts and Endothelial Cells in Rheumatoid Arthritis | 4.22 | 0.154 | 2.188 | 52 |
| Phase II - Conjugation of compounds | 4.15 | 0.211 | -1.46 | 23 |
| Serotonin Degradation | 4.15 | 0.221 | -1.508 | 21 |
| Role of Osteoblasts in Rheumatoid Arthritis Signaling Pathway | 4.12 | 0.167 | 3.042 | 40 |
| PI3K/AKT Signaling | 4.12 | 0.175 | 2.183 | 35 |
| Role of Tissue Factor in Cancer | 4.11 | 0.173 | 4.564 | 36 |
| Chondroitin Sulfate Biosynthesis (Late Stages) | 4.05 | 0.254 | 2.111 | 16 |
| Ephrin Receptor Signaling | 4.03 | 0.173 | 1.213 | 35 |
| Signaling by Rho Family GTPases | 4.03 | 0.161 | 4.49 | 43 |
| Integrin Signaling | 4.03 | 0.171 | 4.017 | 36 |
| Fructose metabolism | 4.01 | 0.714 | -1.342 | 5 |
| Activin Inhibin Signaling Pathway | 3.94 | 0.17 | 4 | 36 |
| Glioma Invasiveness Signaling | 3.92 | 0.239 | 1.291 | 17 |
| Chondroitin Sulfate Biosynthesis | 3.92 | 0.239 | 1.732 | 17 |
| Interleukin-6 family signaling | 3.9 | 0.375 | 1.667 | 9 |
| Cell junction organization | 3.87 | 0.217 | 3.9 | 20 |
| Dilated Cardiomyopathy Signaling Pathway | 3.86 | 0.185 | -1.342 | 28 |
| WNT/SHH Axonal Guidance Signaling Pathway | 3.86 | 0.185 | 1.89 | 28 |
| Adrenergic Receptor Signaling Pathway (Enhanced) | 3.85 | 0.172 | -1.715 | 34 |
| Signaling by MET | 3.82 | 0.228 | 3.771 | 18 |
| HMGB1 Signaling | 3.79 | 0.181 | 2.982 | 29 |
| Dermatan Sulfate Biosynthesis | 3.76 | 0.233 | 1.732 | 17 |
| Actin Cytoskeleton Signaling | 3.72 | 0.161 | 3.13 | 39 |
| IL-12 Signaling and Production in Macrophages | 3.72 | 0.161 | 0.164 | 39 |
| Signaling by ALK | 3.64 | 0.323 | 3.162 | 10 |
| Platelet Adhesion to exposed collagen | 3.62 | 0.438 | 1.89 | 7 |
| Sucrose Degradation V (Mammalian) | 3.61 | 0.625 | -1.342 | 5 |
| Superoxide Radicals Degradation | 3.61 | 0.625 | 1.342 | 5 |
| PAK Signaling | 3.6 | 0.195 | 3.873 | 23 |
| Regulation of Cellular Mechanics by Calpain Protease | 3.6 | 0.213 | 2.333 | 19 |
| GP1b-IX-V activation signalling | 3.56 | 0.5 | 2.449 | 6 |
| Bladder Cancer Signaling | 3.55 | 0.193 | 2.449 | 23 |
| Cell surface interactions at the vascular wall | 3.53 | 0.164 | 5.24 | 35 |
| Retinoate Biosynthesis I | 3.5 | 0.26 | 0 | 13 |
| Immunoregulatory interactions between a Lymphoid and a non-Lymphoid cell | 3.49 | 0.163 | 4.802 | 35 |
| MSP-RON Signaling in Cancer Cells Pathway | 3.49 | 0.182 | 3.266 | 26 |
| Clathrin-mediated Endocytosis Signaling | 3.4 | 0.163 | - | 34 |
| Maturity Onset Diabetes of Young (MODY) Signaling | 3.39 | 0.218 | - | 17 |
| HEY1 Signaling Pathway | 3.38 | 0.174 | 2.117 | 28 |
| ID1 Signaling Pathway | 3.35 | 0.163 | 2.335 | 33 |
| Transport of inorganic cations/anions and amino acids/oligopeptides | 3.33 | 0.194 | 1.091 | 21 |
| Caveolar-mediated Endocytosis Signaling | 3.33 | 0.215 | 3.742 | 17 |
| NRF2-mediated Oxidative Stress Response | 3.32 | 0.152 | 0 | 41 |
| Platelet Aggregation (Plug Formation) | 3.32 | 0.462 | 1.633 | 6 |
| Atorvastatin ADME | 3.3 | 0.556 | -2.236 | 5 |
| Apelin Adipocyte Signaling Pathway | 3.28 | 0.202 | 0.775 | 19 |
| Apelin Cardiac Fibroblast Signaling Pathway | 3.26 | 0.348 | -1.89 | 8 |
| Metabolism of amine-derived hormones | 3.25 | 0.389 | -1.134 | 7 |
| IL-17 Signaling | 3.22 | 0.164 | 4.158 | 31 |
| Dermatan Sulfate Biosynthesis (Late Stages) | 3.21 | 0.233 | 1.897 | 14 |
| Paracetamol ADME | 3.19 | 0.31 | -0.333 | 9 |
| Acute Phase Response Signaling | 3.14 | 0.162 | 2.065 | 31 |
| Ion channel transport | 3.13 | 0.164 | 2.921 | 30 |
| IL-8 Signaling | 3.12 | 0.159 | 4.271 | 33 |
| Agrin Interactions at Neuromuscular Junction | 3.12 | 0.221 | 2.53 | 15 |
| Role of IL-17A in Psoriasis | 3.11 | 0.429 | 1.342 | 6 |
| Docosahexaenoic Acid (DHA) Signaling | 3.1 | 0.151 | -0.333 | 38 |
| Production of Nitric Oxide and Reactive Oxygen Species in Macrophages | 3.1 | 0.161 | 1.134 | 31 |
| Choline catabolism | 3.1 | 0.667 | -2 | 4 |
| Trehalose Degradation II (Trehalase) | 3.1 | 0.667 | - | 4 |
| GABA synthesis, release, reuptake and degradation | 3.08 | 0.368 | 0.378 | 7 |
| Bile Acid Biosynthesis, Neutral Pathway | 3.08 | 0.368 | -2.236 | 7 |
| Complement cascade | 3.07 | 0.176 | 2.041 | 24 |
| Prednisone ADME | 3.03 | 0.5 | -2.236 | 5 |
| Actin Nucleation by ARP-WASP Complex | 3.03 | 0.198 | 1.667 | 18 |
| G alpha (s) signalling events | 3.02 | 0.172 | 1.8 | 25 |
| Cachexia Signaling Pathway | 3.02 | 0.139 | 3.286 | 51 |
| Role of Osteoblasts, Osteoclasts and Chondrocytes in Rheumatoid Arthritis | 2.98 | 0.153 | - | 35 |
| Detoxification of Reactive Oxygen Species | 2.96 | 0.27 | 2.53 | 10 |
| Glioblastoma Multiforme Signaling | 2.95 | 0.164 | 2.294 | 28 |
| EPH-Ephrin signaling | 2.91 | 0.194 | 3.3 | 18 |
| GABAergic Receptor Signaling Pathway (Enhanced) | 2.89 | 0.171 | 0.209 | 24 |
| Regulation of the Epithelial Mesenchymal Transition in Development Pathway | 2.89 | 0.198 | 2.138 | 17 |
| G alpha (i) signalling events | 2.87 | 0.147 | 3.569 | 38 |
| WNT/β-catenin Signaling | 2.87 | 0.162 | -0.392 | 28 |
| Xenobiotic Metabolism AHR Signaling Pathway | 2.87 | 0.183 | 0 | 20 |
| Ethanol Degradation IV | 2.86 | 0.308 | -0.707 | 8 |
| GABA Receptor Signaling | 2.85 | 0.173 | - | 23 |
| NFE2L2 regulating tumorigenic genes | 2.8 | 0.455 | 2.236 | 5 |
| Oxidative Ethanol Degradation III | 2.78 | 0.212 | -0.816 | 14 |
| Calcium Signaling | 2.77 | 0.151 | 1.807 | 33 |
| Role of IL-17F in Allergic Inflammatory Airway Diseases | 2.73 | 0.239 | 2.449 | 11 |
| PXR/RXR Activation | 2.71 | 0.209 | -0.816 | 14 |
| Glutaminergic Receptor Signaling Pathway (Enhanced) | 2.7 | 0.138 | 2.111 | 45 |
| IL-6 Signaling | 2.67 | 0.171 | 1.606 | 22 |
| Extra-nuclear estrogen signaling | 2.66 | 0.2 | 3.357 | 15 |
| Signaling by TGF-beta Receptor Complex | 2.63 | 0.222 | 3.464 | 12 |
| Cardiac conduction | 2.63 | 0.169 | 1.291 | 22 |
| STAT3 Pathway | 2.59 | 0.165 | 2.496 | 23 |
| Neurotransmitter uptake and metabolism In glial cells | 2.59 | 0.75 | - | 3 |
| Catecholamine Biosynthesis | 2.59 | 0.75 | - | 3 |
| NAFLD Signaling Pathway | 2.58 | 0.147 | 2.959 | 33 |
| Sensory processing of sound by outer hair cells of the cochlea | 2.56 | 0.218 | 0.577 | 12 |
| FAK Signaling | 2.56 | 0.114 | 5.199 | 118 |
| Glycolysis I | 2.53 | 0.276 | 0 | 8 |
| Heparan Sulfate Biosynthesis | 2.5 | 0.178 | 1.732 | 18 |
| VDR/RXR Activation | 2.49 | 0.192 | 0.333 | 15 |
| Heparan Sulfate Biosynthesis (Late Stages) | 2.46 | 0.181 | 2.111 | 17 |
| Formation of the nephric duct | 2.45 | 0.333 | 0.816 | 6 |
| Transport of vitamins, nucleosides, and related molecules | 2.43 | 0.233 | 0 | 10 |
| Intrinsic Prothrombin Activation Pathway | 2.43 | 0.233 | 1 | 10 |
| Androgen Biosynthesis | 2.42 | 0.292 | -2 | 7 |
| Glycerophospholipid biosynthesis | 2.37 | 0.164 | 0.218 | 21 |
| Role of IL-17A in Arthritis | 2.36 | 0.207 | - | 12 |
| Oxytocin Signaling Pathway | 2.36 | 0.137 | 4.542 | 39 |
| Netrin-1 signaling | 2.36 | 0.227 | 1.897 | 10 |
| Transcriptional regulation by RUNX2 | 2.35 | 0.216 | 0.333 | 11 |
| RAC Signaling | 2.34 | 0.161 | 3.606 | 22 |
| Germ Cell-Sertoli Cell Junction Signaling | 2.33 | 0.152 | - | 26 |
| Glyoxylate metabolism and glycine degradation | 2.32 | 0.316 | -2.449 | 6 |
| BBSome Signaling Pathway | 2.3 | 0.124 | 2.994 | 61 |
| Semaphorin Signaling in Neurons | 2.3 | 0.203 | - | 12 |
| Interleukin-10 signaling | 2.28 | 0.222 | 3.162 | 10 |
| Cardiac Hypertrophy Signaling | 2.28 | 0.138 | 1.732 | 36 |
| Parkinson's Signaling Pathway | 2.27 | 0.133 | 1.093 | 42 |
| Paxillin Signaling | 2.27 | 0.17 | 2.714 | 18 |
| RAF/MAP kinase cascade | 2.25 | 0.137 | 4 | 36 |
| PCP (Planar Cell Polarity) Pathway | 2.24 | 0.2 | 1.732 | 12 |
| Taurine Biosynthesis | 2.22 | 0.6 | - | 3 |
| IL-17A Signaling in Airway Cells | 2.21 | 0.191 | 1.89 | 13 |
| IL-17A Signaling in Gastric Cells | 2.21 | 0.269 | 1.633 | 7 |
| Eicosanoid Signaling | 2.2 | 0.135 | 3.244 | 38 |
| Plasma lipoprotein assembly, remodeling, and clearance | 2.19 | 0.184 | -1.604 | 14 |
| ERK/MAPK Signaling | 2.17 | 0.141 | 1.706 | 31 |
| Airway Inflammation in Asthma | 2.16 | 0.242 | 2 | 8 |
| CXCR4 Signaling | 2.15 | 0.149 | 2.828 | 25 |
| Neurotransmitter release cycle | 2.14 | 0.225 | 1 | 9 |
| Factors Promoting Cardiogenesis in Vertebrates | 2.14 | 0.152 | 1.706 | 23 |
| Neuregulin Signaling | 2.11 | 0.161 | 3 | 19 |
| Glutamine Degradation I | 2.1 | 1 | - | 2 |
| eNOS Signaling | 2.09 | 0.149 | 0.775 | 24 |
| Creatine metabolism | 2.08 | 0.4 | 0 | 4 |
| Glucose and Glucose-1-phosphate Degradation | 2.08 | 0.4 | - | 4 |
| PIP3 activates AKT signaling | 2.05 | 0.152 | 2.558 | 22 |
| Gap Junction Signaling | 2.05 | 0.128 | 0.156 | 43 |
| Class B/2 (Secretin family receptors) | 2.04 | 0.168 | 1 | 16 |
| Role of MAPK Signaling in Inhibiting the Pathogenesis of Influenza | 2.04 | 0.177 | 1.387 | 14 |
| Autism Signaling Pathway | 2 | 0.129 | 2.271 | 40 |
| CGAS-STING Signaling Pathway | 1.99 | 0.152 | 3.13 | 21 |
| Extrinsic Prothrombin Activation Pathway | 1.98 | 0.312 | - | 5 |
| G alpha (q) signalling events | 1.96 | 0.144 | 3 | 25 |
| Lipid particle organization | 1.95 | 0.5 | - | 3 |
| HGF Signaling | 1.94 | 0.153 | 2.53 | 20 |
| CDK5 Signaling | 1.93 | 0.16 | -0.333 | 17 |
| Striated Muscle Contraction | 1.92 | 0.222 | 2.121 | 8 |
| IL-13 Signaling Pathway | 1.92 | 0.154 | 0.728 | 19 |
| BEX2 Signaling Pathway | 1.9 | 0.171 | 1.155 | 14 |
| ERK5 Signaling | 1.9 | 0.176 | 3.317 | 13 |
| Nuclear Cytoskeleton Signaling Pathway | 1.9 | 0.136 | 3.651 | 30 |
| Th1 and Th2 Activation Pathway | 1.9 | 0.142 | - | 25 |
| Neutrophil degranulation | 1.89 | 0.119 | 4.371 | 57 |
| FAT10 Cancer Signaling Pathway | 1.89 | 0.196 | 2.333 | 10 |
| Azathioprine ADME | 1.89 | 0.261 | 0 | 6 |
| NFE2L2 regulating anti-oxidant/detoxification enzymes | 1.89 | 0.261 | 0.816 | 6 |
| L1CAM interactions | 1.89 | 0.153 | 3.441 | 19 |
| Microautophagy Signaling Pathway | 1.88 | 0.145 | 1 | 23 |
| Sleep NREM Signaling Pathway | 1.87 | 0.155 | 0.5 | 18 |
| Metabolism of folate and pterines | 1.87 | 0.294 | -0.447 | 5 |
| cAMP-mediated signaling | 1.86 | 0.133 | -0.577 | 32 |
| Estrogen-Dependent Breast Cancer Signaling | 1.86 | 0.169 | 1.633 | 14 |
| CD40 Signaling | 1.86 | 0.179 | 2.53 | 12 |
| Tryptophan Degradation X (Mammalian, via Tryptamine) | 1.85 | 0.233 | -0.816 | 7 |
| Pancreatic Adenocarcinoma Signaling | 1.85 | 0.152 | 2.887 | 19 |
| Gα12/13 Signaling | 1.84 | 0.149 | 3.153 | 20 |
| Amyotrophic Lateral Sclerosis Signaling | 1.83 | 0.154 | 1.155 | 18 |
| Cerebral Malformation Signaling Pathway | 1.83 | 0.154 | 2.828 | 18 |
| Cholecystokinin/Gastrin-mediated Signaling | 1.83 | 0.154 | 2.5 | 18 |
| Hepatitis B Chronic Liver Pathogenesis Signaling Pathway | 1.81 | 0.138 | 3.922 | 26 |
| PPARα/RXRα Activation | 1.79 | 0.136 | -1.342 | 27 |
| ERBB Signaling | 1.78 | 0.161 | 3.207 | 15 |
| Metabolism of steroid hormones | 1.78 | 0.211 | 0 | 8 |
| Reversible hydration of carbon dioxide | 1.77 | 0.333 | 0 | 4 |
| Specification of primordial germ cells | 1.77 | 0.333 | 2 | 4 |
| Apelin Endothelial Signaling Pathway | 1.76 | 0.145 | 2.84 | 21 |
| Sensory processing of sound by inner hair cells of the cochlea | 1.76 | 0.174 | 0.577 | 12 |
| ROBO SLIT Signaling Pathway | 1.75 | 0.148 | -0.471 | 19 |
| Retinol Biosynthesis | 1.72 | 0.185 | -1.633 | 10 |
| Potassium Channels | 1.72 | 0.155 | 0.5 | 16 |
| Multiple Sclerosis Signaling Pathway | 1.72 | 0.132 | 3.157 | 29 |
| Role of PKR in Interferon Induction and Antiviral Response | 1.71 | 0.145 | 2.324 | 20 |
| Semaphorin Neuronal Repulsive Signaling Pathway | 1.7 | 0.143 | 0.471 | 21 |
| Surfactant metabolism | 1.7 | 0.219 | 0.378 | 7 |
| Gustation Pathway | 1.69 | 0.133 | 2.502 | 28 |
| Neuroprotective Role of THOP1 in Alzheimer's Disease | 1.68 | 0.146 | 1.155 | 19 |
| Glucocorticoid Receptor Signaling | 1.67 | 0.113 | - | 68 |
| Uracil Degradation II (Reductive) | 1.65 | 0.667 | - | 2 |
| Serine Biosynthesis | 1.65 | 0.667 | - | 2 |
| Thyronamine and Iodothyronamine Metabolism | 1.65 | 0.667 | - | 2 |
| Thymine Degradation | 1.65 | 0.667 | - | 2 |
| Thyroid Hormone Metabolism I (via Deiodination) | 1.65 | 0.667 | - | 2 |
| Dopamine Degradation | 1.65 | 0.2 | -0.378 | 8 |
| Cholesterol Biosynthesis I | 1.64 | 0.308 | -2 | 4 |
| Cholesterol Biosynthesis II (via 24,25-dihydrolanosterol) | 1.64 | 0.308 | -2 | 4 |
| Cholesterol Biosynthesis III (via Desmosterol) | 1.64 | 0.308 | -2 | 4 |
| TNFR2 Signaling | 1.63 | 0.212 | 2 | 7 |
| IL-33 Signaling Pathway | 1.63 | 0.133 | 3.838 | 26 |
| Metabolism of porphyrins | 1.63 | 0.231 | -2.449 | 6 |
| Thrombin Signaling | 1.62 | 0.13 | 2.065 | 29 |
| CD27 Signaling in Lymphocytes | 1.62 | 0.179 | 2.121 | 10 |
| Ovarian Cancer Signaling | 1.62 | 0.138 | 1.89 | 22 |
| Sleep REM Signaling Pathway | 1.61 | 0.151 | -1.5 | 16 |
| Myelination Signaling Pathway | 1.61 | 0.122 | 4.427 | 40 |
| DAG and IP3 signaling | 1.59 | 0.195 | 1.134 | 8 |
| Chronic Myeloid Leukemia Signaling | 1.59 | 0.125 | 3.43 | 35 |
| Opioid Signaling Pathway | 1.59 | 0.125 | 0.756 | 35 |
| IL-1 Signaling | 1.59 | 0.153 | 1.414 | 15 |
| PTEN Signaling | 1.59 | 0.139 | -2.496 | 21 |
| PFKFB4 Signaling Pathway | 1.57 | 0.184 | 0.378 | 9 |
| Role of JAK2 in Hormone-like Cytokine Signaling | 1.57 | 0.169 | 0.333 | 11 |
| Tuberculosis Latent Signaling Pathway | 1.57 | 0.156 | 0 | 14 |
| Folate Signaling Pathway | 1.57 | 0.175 | 1.667 | 10 |
| Apelin Liver Signaling Pathway | 1.56 | 0.25 | 2.236 | 5 |
| Type II Diabetes Mellitus Signaling | 1.56 | 0.138 | 1 | 21 |
| Formation of intermediate mesoderm | 1.56 | 0.375 | - | 3 |
| PPAR Signaling | 1.55 | 0.148 | -2.673 | 16 |
| April Mediated Signaling | 1.53 | 0.19 | 2.646 | 8 |
| Glutamate and glutamine metabolism | 1.52 | 0.286 | 0 | 4 |
| Triacylglycerol Biosynthesis | 1.52 | 0.172 | -0.707 | 10 |
| Tight Junction Signaling | 1.52 | 0.133 | - | 24 |
| Erythropoietin Signaling Pathway | 1.52 | 0.133 | -0.426 | 24 |
| Human Embryonic Stem Cell Pluripotency | 1.51 | 0.13 | 2.746 | 26 |
| B Cell Activating Factor Signaling | 1.48 | 0.186 | 2.449 | 8 |
| Sulfur amino acid metabolism | 1.48 | 0.214 | -1.633 | 6 |
| RHO GTPases activate PKNs | 1.48 | 0.214 | 2.449 | 6 |
| CDP-diacylglycerol Biosynthesis I | 1.48 | 0.214 | 1 | 6 |
| Cyclophilin Signaling Pathway | 1.47 | 0.124 | 3.053 | 31 |
| GPCR-Mediated Integration of Enteroendocrine Signaling Exemplified by an L Cell | 1.46 | 0.158 | -0.577 | 12 |
| NOD1/2 Signaling Pathway | 1.45 | 0.13 | 3.838 | 25 |
| GNRH Signaling | 1.45 | 0.13 | 1.886 | 25 |
| SNARE Signaling Pathway | 1.44 | 0.138 | 2.065 | 19 |
| Virus Entry via Endocytic Pathways | 1.44 | 0.142 | 3.5 | 17 |
| MSP-RON Signaling in Macrophages Pathway | 1.44 | 0.142 | 1.291 | 17 |
| Activation of NMDA receptors and postsynaptic events | 1.44 | 0.153 | 0.302 | 13 |
| Netrin Signaling | 1.44 | 0.131 | 2.837 | 23 |
| GADD45 Signaling | 1.43 | 0.167 | -0.632 | 10 |
| YAP1- and WWTR1 (TAZ)-stimulated gene expression | 1.42 | 0.267 | 2 | 4 |
| Peroxisomal lipid metabolism | 1.41 | 0.207 | -2.449 | 6 |
| TNFs bind their physiological receptors | 1.41 | 0.207 | 1.633 | 6 |
| EGR2 and SOX10-mediated initiation of Schwann cell myelination | 1.41 | 0.207 | 0.816 | 6 |
| Serine biosynthesis | 1.41 | 0.333 | - | 3 |
| Histidine Degradation III | 1.41 | 0.333 | - | 3 |
| GDP-glucose Biosynthesis | 1.41 | 0.333 | - | 3 |
| Epithelial Adherens Junction Signaling | 1.4 | 0.133 | 0.894 | 21 |
| Fatty Acid α-oxidation | 1.4 | 0.227 | -0.447 | 5 |
| Senescence Pathway | 1.39 | 0.12 | 3.286 | 36 |
| Thyroid Cancer Signaling | 1.38 | 0.154 | 2.309 | 12 |
| Retinoate Biosynthesis II | 1.38 | 0.5 | - | 2 |
| D-glucuronate Degradation I | 1.38 | 0.5 | - | 2 |
| L-cysteine Degradation I | 1.38 | 0.5 | - | 2 |
| Signaling by EGFR | 1.38 | 0.17 | 2.333 | 9 |
| Estrogen Receptor Signaling | 1.37 | 0.114 | 3.479 | 47 |
| Regulation of CDH11 Expression and Function | 1.35 | 0.2 | 2.449 | 6 |
| Glutathione Redox Reactions I | 1.35 | 0.2 | 1 | 6 |
| Phosphatidylglycerol Biosynthesis II (Non-plastidic) | 1.35 | 0.2 | 1 | 6 |
| Thyroid Hormone Metabolism II (via Conjugation and/or Degradation) | 1.35 | 0.161 | 0 | 10 |
| Acetylcholine Receptor Signaling Pathway | 1.34 | 0.126 | 1.225 | 25 |
| Amyloid fiber formation | 1.34 | 0.148 | 1.941 | 13 |
| Death Receptor Signaling | 1.33 | 0.144 | 1.155 | 14 |
| TGF-β Signaling | 1.33 | 0.144 | 0.632 | 14 |
| Triglyceride metabolism | 1.33 | 0.184 | -2.646 | 7 |
| MIF Regulation of Innate Immunity | 1.33 | 0.174 | 1.89 | 8 |
| IL-23 Signaling Pathway | 1.33 | 0.174 | 0.378 | 8 |
| Histamine Degradation | 1.32 | 0.217 | -1 | 5 |
| Th1 Pathway | 1.3 | 0.136 | 1.698 | 17 |

Supplemental Figures

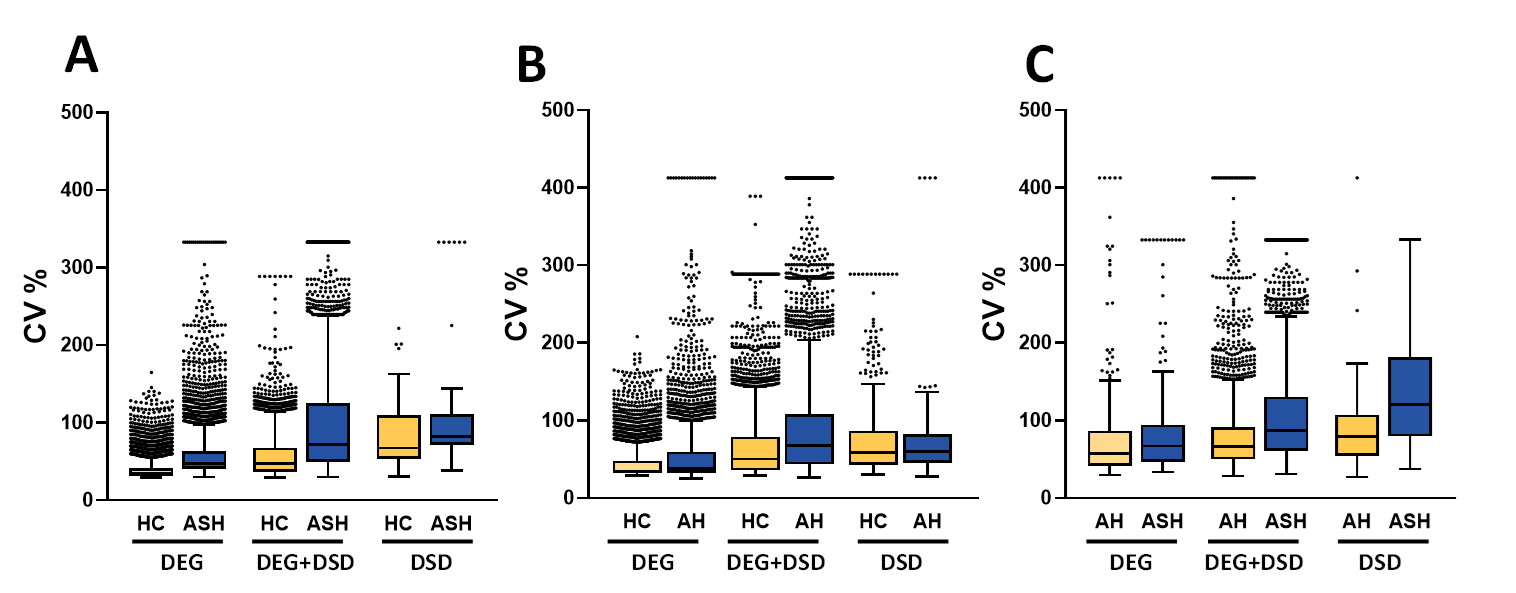

Figure S1

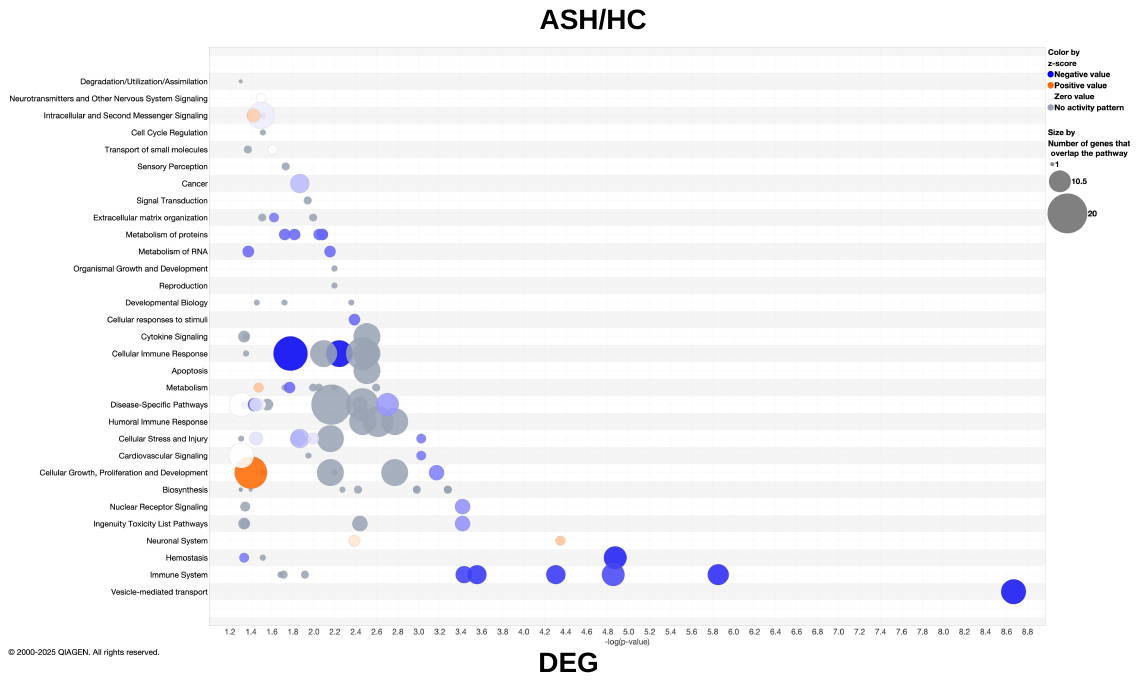

Figure S2

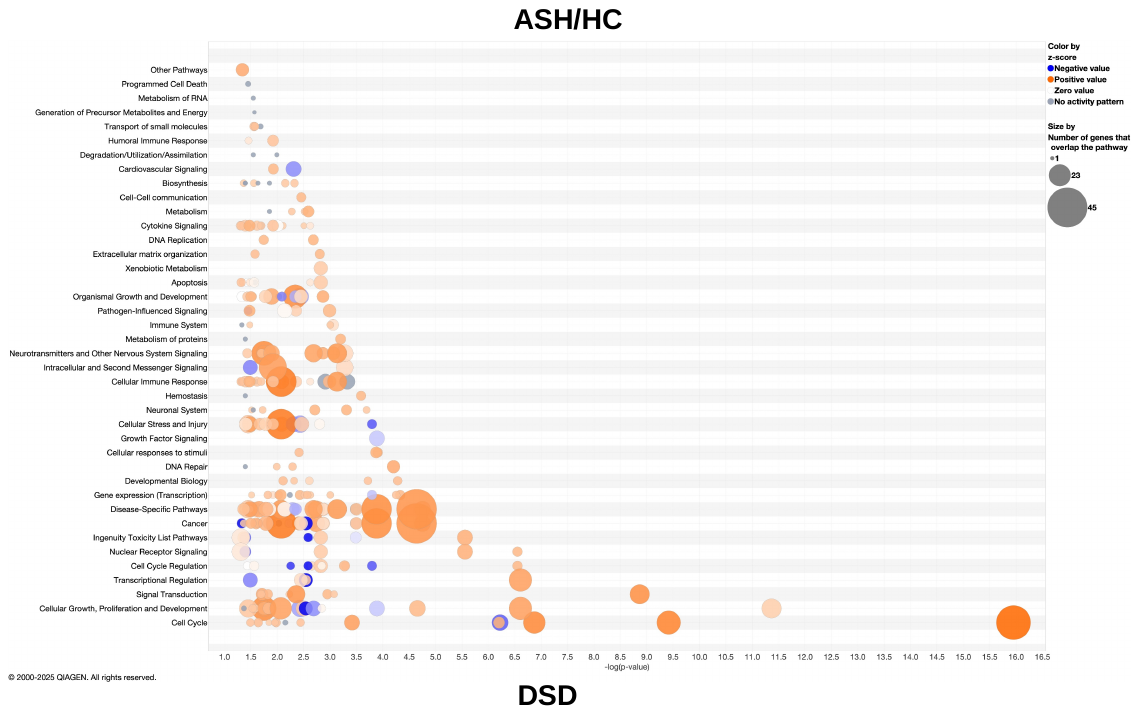

Figure S2

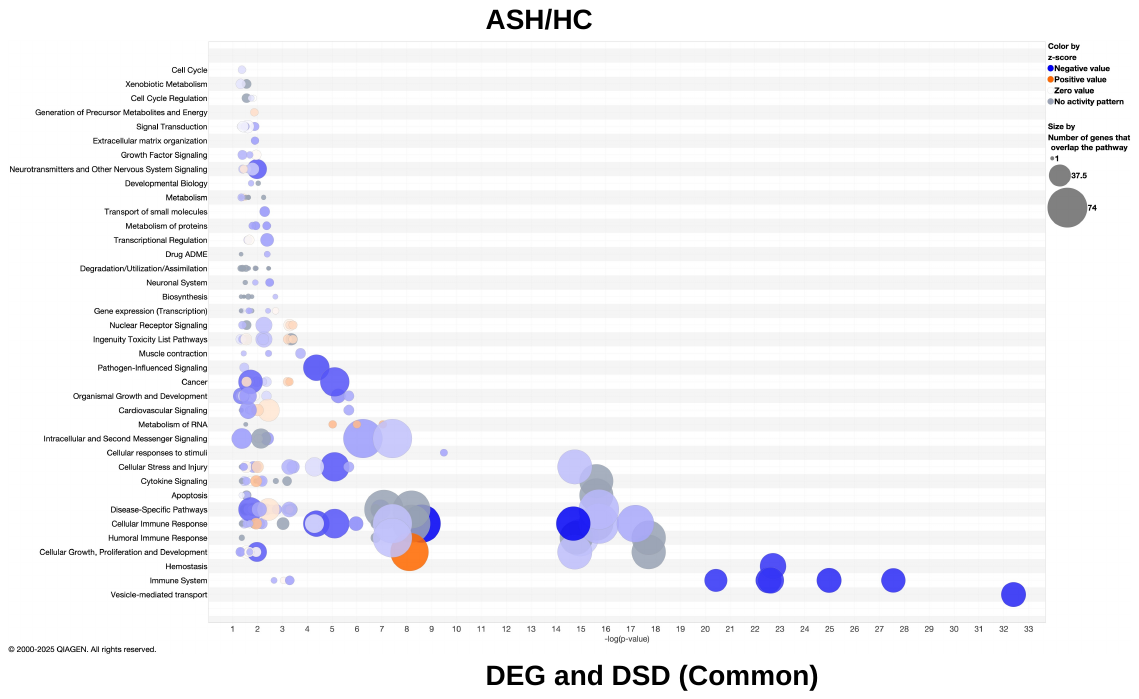

Figure S2

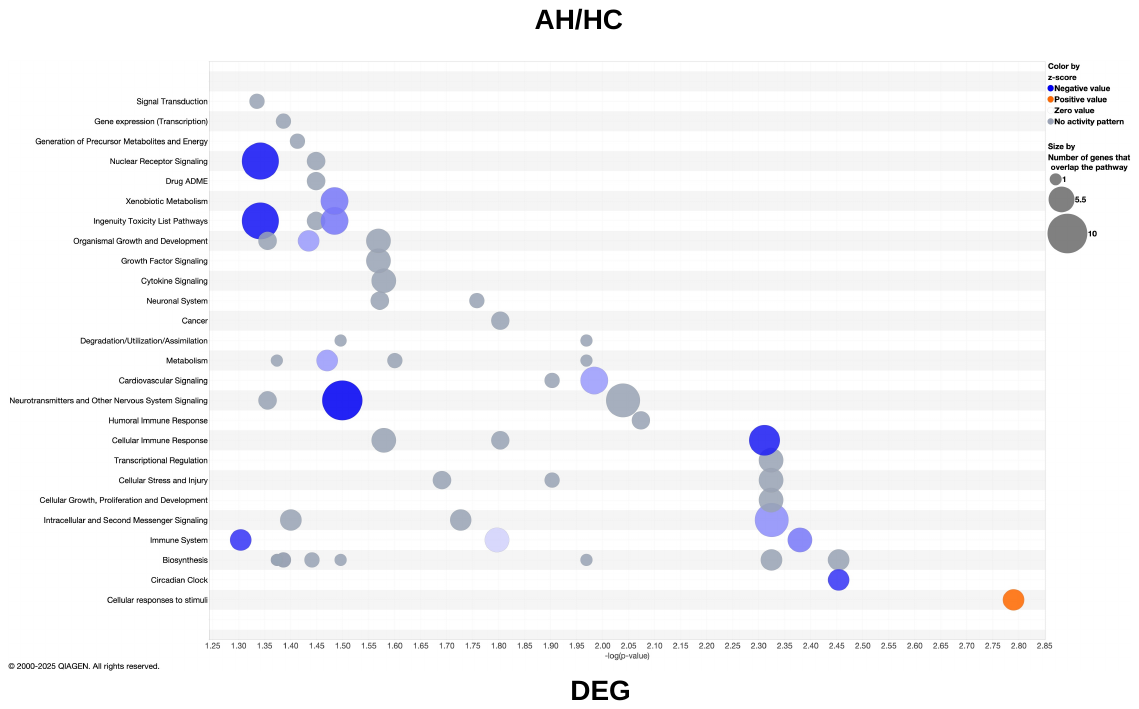

Figure S3

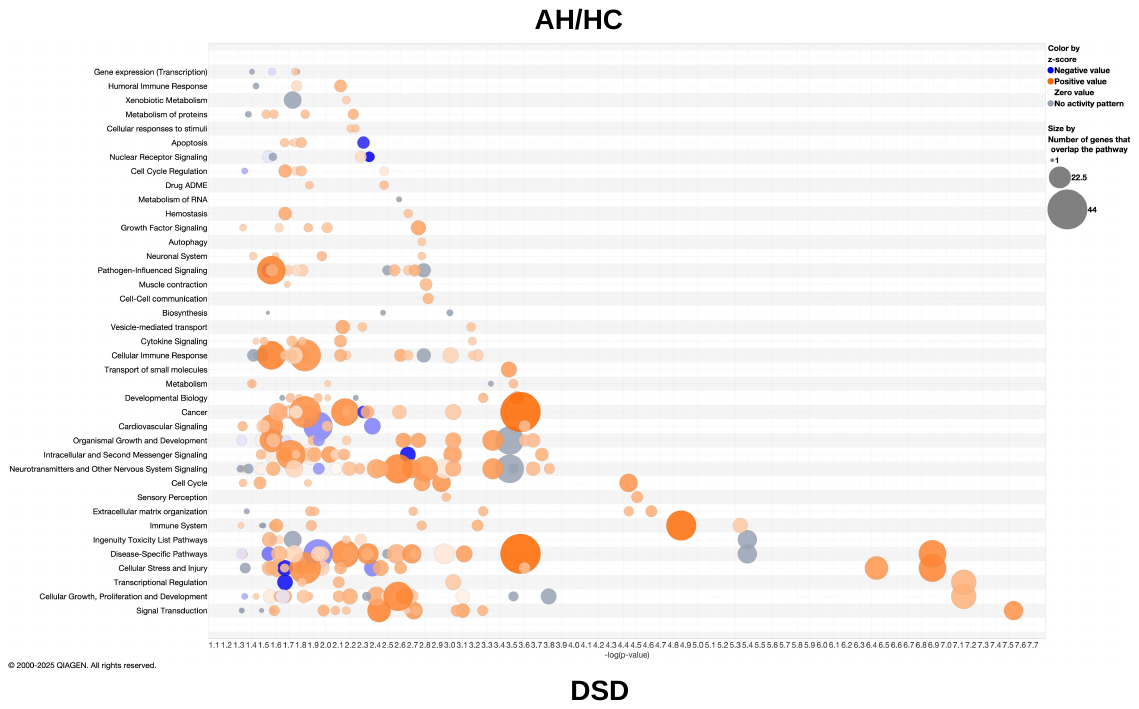

Figure S3

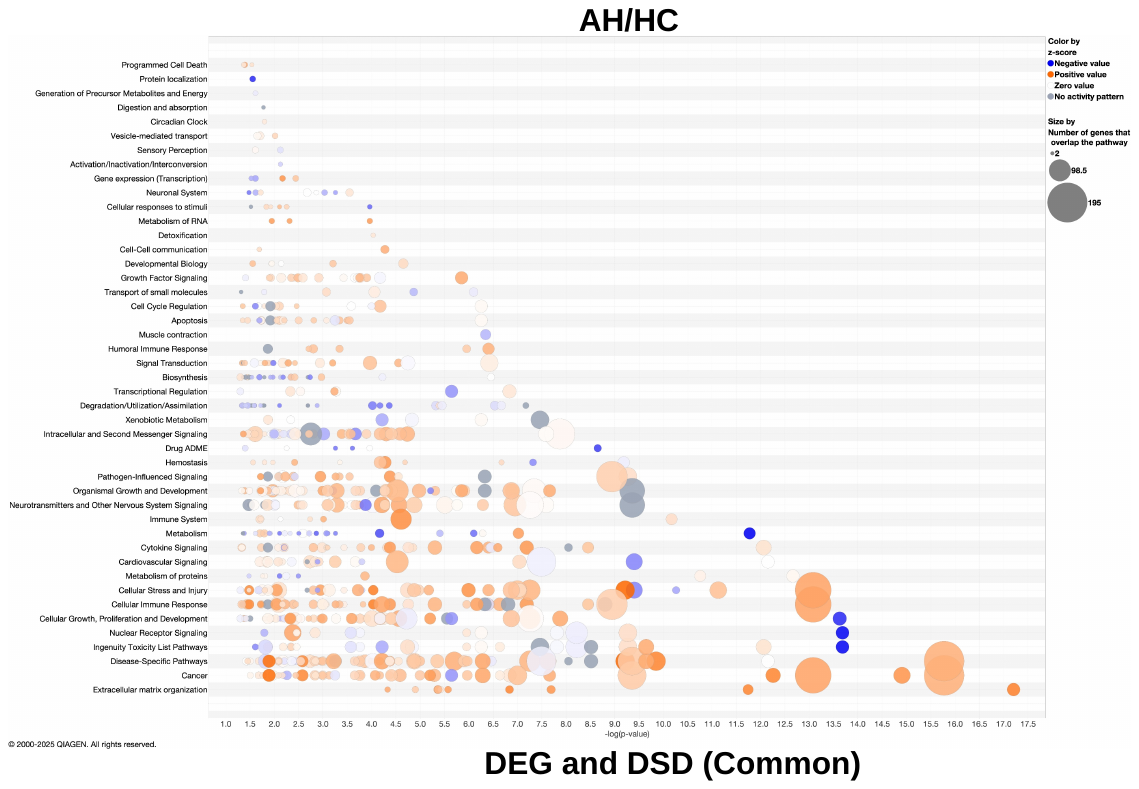

Figure S3

Supplemental Figure legends

Figure S1(A-C): Variability represented by the percentage of CV (coefficient of variation, that is standard deviation divided by mean) of three abundance distributions ASH vs HC (S1), AH vs HC (S2) and AH vs ASH (S3) for genes exclusive to DEG and DSD and common and shared by both DEG and DSD analysis.

Figure S2, S3: Canonical Pathway bubble chart. Canonical Pathway scores plotted as pathway category based on gene expression level for the DEG and DSD exclusive and common pathways in the ASH vs. HC and AH vs. HC group. The log of the p-value for each pathway is plotted on the x-axis versus the pathway categories plotted along the y-axis. By default, the coloring relates to the pathway's z-score, and the bubble size relates to the number of dataset genes that overlap each pathway, as shown in the legend.
